## Supplementary Data 1 for "Regulation of an iron-dependent repressor by tryptophan availability attenuates transcription of the tryptophan salvage genes in *Chlamydia trachomatis*"

> Escherichia coli strain K12

MKAIFVLKGWWRTS

RBS: AAAGGGTA

Length: 14

>Yersinia pestis strain FDAARGOS_601

MKTSLISLLRWWHISLSRAM

RBS: AAAGAGAG

Length: 20

>Vibrio cholerae strain 10432-62

MLQEFNPNHKPNFSPADAELAWWRTWTSSWWAHVYF

RBS: TGTTAGGG

Length: 36

>Pseudoalteromonas haloplanktis TAC125

MNNNLTLTHRWWRLI

RBS: ATTAGAGG

Length: 15

>Shewanella amazonensis SB2B

MNPIIASFINWWWHFPNTRVV

RBS: AAGATAAA

Length: 21

>Pseudomonas aeruginosa PAO1

Excluded due to lack of obvious trpL upstream of either trpE or trpG

>Bordetella parapertussis Bpp01

MNARPSNQMRNASWRWWRLSSGWR

RBS: AGACACTG

Length: 24

>Acidovorax citrulli AAC00-1

MTCSASFNVAKWWRFS

RBS: CAACTGGC

Length: 16

>Brucella melitensis bv. 1 str. 16M

MLNANSTSMNISRIIVINGWWWAR

RBS: TTGCAAGG

Length: 24

>Ochrobactrum anthropic ATCC 49188

MNISRNIVINGWWWAR

RBS: TAACAGTA

Length: 16

>Rhizobium etli CFN 42

MIKSLNIAVWWWAR

RBS: AGGCTAAC

Length: 14

>Agrobacterium tumefaciens strain Ach5

MNIVSMNIANWWWSSFTRP

RBS: AGGCTAAC

Length: 19

>Corynebacterium glutamicum stain ATCC 13032

Excluded due to lack of obvious trpL upstream of either trpE

>Streptomyces coelicolor A3(2)

MFAHSTRNWWWTAHPAAH

RBS: AGGGTCGG

Length: 18

>Chlamydia trachomatis L2 434/Bu

MHALLMNKYSVLAVLVRKYSCSMPCKSAFQADCFQDIQKFILLQRAWLSFESWRLSTWR

RBS: GATTGAAA

Length: 59

>Deinococcus radiodurans R1 – trpGD*

MSARTLRLIWWPRLT

RBS: No good candidate

Length: 15
