## Supplementary Note 1 for "Regulation of an iron-dependent repressor by tryptophan availability attenuates transcription of the tryptophan salvage genes in *Chlamydia trachomatis*"

### Figure 1

Nick Pokorzynski

4/14/2020

```
#Fig 1b
setwd("~/Documents/Carabeo Lab/R Code/Paper 2/Figure 1")
library(ggplot2)
library(ggpubr)

## Warning: package 'ggpubr' was built under R version 3.5.2

## Loading required package: magrittr

gdna<-read.csv("Fig1b.csv")
gdna$Treatment2<-factor(gdna$Treatment, levels=c("Mock", "Bpd", "Trp"))
gdna$ID2<-factor(gdna$ID, levels=c("24 hpi", "18+6 Bpd", "18+6 Trp-", "24
Bpd", "24 Trp-"))
gdnaaov<-aov(Value~ID, data=gdna)
summary(gdnaaov)

##             Df    Sum Sq   Mean Sq F value   Pr(>F)    
## ID           4 1.125e+10 2.813e+09   91.35 7.87e-08 ***
## Residuals   10 3.079e+08 3.079e+07                     
## ---
## Signif. codes:  0 '***' 0.001 '**' 0.01 '*' 0.05 '.' 0.1 ' ' 1

TukeyHSD(x=gdnaaov)

##   Tukey multiple comparisons of means
##     95% family-wise confidence level
## 
## Fit: aov(formula = Value ~ ID, data = gdna)
## 
## $ID
##                          diff       lwr       upr     p adj
## 18+6 Trp--18+6 Bpd   1933.792 -12977.87  16845.46 0.9919733
## 24 Bpd-18+6 Bpd    -43816.909 -58728.57 -28905.24 0.0000166
## 24 hpi-18+6 Bpd     28494.208  13582.54  43405.87 0.0006667
## 24 Trp--18+6 Bpd   -39745.907 -54657.57 -24834.24 0.0000399
## 24 Bpd-18+6 Trp-   -45750.700 -60662.36 -30839.04 0.0000112
## 24 hpi-18+6 Trp-    26560.416  11648.75  41472.08 0.0011596
## 24 Trp--18+6 Trp-  -41679.698 -56591.36 -26768.03 0.0000261
## 24 hpi-24 Bpd       72311.117  57399.45  87222.78 0.0000002
## 24 Trp--24 Bpd       4071.002 -10840.66  18982.67 0.8911029
## 24 Trp--24 hpi     -68240.115 -83151.78 -53328.45 0.0000003
```

```
ggbarplot(gdna, x="ID2", y="Log10_Value", add="mean_sd", fill="Treatment2",
palette=c("lightsteelblue4", "lightsteelblue1", "lightsteelblue3")) +
scale_y_continuous(expand=expand_scale(mult=c(0,0.1))) + ylab(label="Log10
Genome Equivalents/ng gDNA") + rremove("legend") +
theme(axis.text.x=element_text(angle=45, hjust=1)) + rremove("xlab") +
geom_exec(geomfunc=geom_point, data=gdna, x="ID2", y="Log10_Value",
colour="black") + geom_signif(comparisons=list(c("18+6 Bpd", "18+6 Trp-"),
c("24 hpi", "18+6 Bpd"), c("24 hpi", "18+6 Trp-"), c("24 Bpd", "24 Trp-"),
c("24 hpi", "24 Bpd"), c("24 hpi", "24 Trp-")), annotation=c("ns", "***",
"**", "ns", "***", "***"), y_position=c(5, 5.5, 6, 4, 6.5, 7)) +
theme(line=element_line(size=1), colour="black"),
panel.border=element_rect(colour="black", fill="NA", size=1))
```

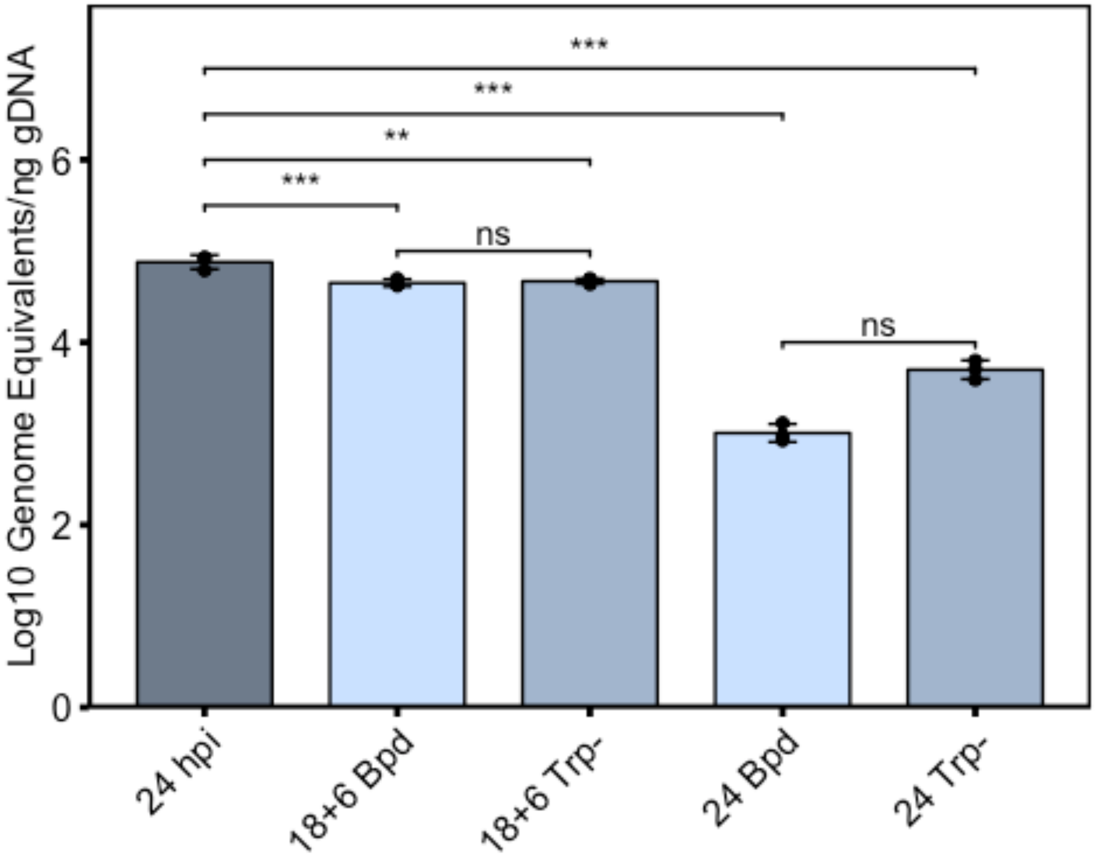


```
#Fig 1c
setwd("~/Documents/Carabeo Lab/R Code/Paper 2/Figure 1")
library(ggplot2)
library(ggpubr)
rt<-read.csv("Fig1c.csv")
rt$Treatment2<-factor(rt$Treatment, levels=c("Mock", "Bpd", "Trp"))
rt$ID2<-factor(rt$ID, levels=c("24 hpi", "18+6 Bpd", "18+6 Trp-", "24 Bpd",
"24 Trp-"))
omcB<-subset(rt, Gene%in%c("omcB"))
```

```
omcBaov<-aov(Log2_Value~ID, data=omcB)
summary(omcBaov)

##             Df Sum Sq Mean Sq F value   Pr(>F)    
## ID           4 24.309   6.077   34.09 8.42e-06 ***
## Residuals   10  1.783   0.178                     
## ---
## Signif. codes:  0 '***' 0.001 '**' 0.01 '*' 0.05 '.' 0.1 ' ' 1

TukeyHSD(x=omcBaov)

##   Tukey multiple comparisons of means
##     95% family-wise confidence level
## 
## Fit: aov(formula = Log2_Value ~ ID, data = omcB)
## 
## $ID
##                          diff        lwr        upr     p adj
## 18+6 Trp--18+6 Bpd -0.4241685 -1.5587295  0.7103924 0.7354420
## 24 Bpd-18+6 Bpd    -2.9528710 -4.0874320 -1.8183101 0.0000492
## 24 hpi-18+6 Bpd    -0.3230207 -1.4575816  0.8115403 0.8761407
## 24 Trp--18+6 Bpd   -2.7005289 -3.8350899 -1.5659680 0.0001072
## 24 Bpd-18+6 Trp-   -2.5287025 -3.6632634 -1.3941415 0.0001879
## 24 hpi-18+6 Trp-    0.1011479 -1.0334131  1.2357088 0.9980944
## 24 Trp--18+6 Trp-  -2.2763604 -3.4109213 -1.1417994 0.0004497
## 24 hpi-24 Bpd       2.6298503  1.4952894  3.7644113 0.0001347
## 24 Trp--24 Bpd      0.2523421 -0.8822189  1.3869030 0.9439768
## 24 Trp--24 hpi     -2.3775082 -3.5120692 -1.2429473 0.0003147

ggbarplot(omcB, x="ID2", y="Log2_Value", add="mean_sd", fill="Treatment2",
palette=c("lightsteelblue4", "lightsteelblue1", "lightsteelblue3"),
title="omcB") + scale_y_continuous(expand=expand_scale(mult=c(0,0.1))) +
ylab(label="Log2 Transcript Expression") + rremove("legend") +
theme(axis.text.x=element_text(angle=45, hjust=1)) + rremove("xlab") +
geom_exec(geomfunc=geom_point, data=omcB, x="ID2", y="Log2_Value",
colour="black") + geom_signif(comparisons=list(c("24 hpi", "18+6 Bpd"), c("24
hpi", "18+6 Trp-"), c("24 hpi", "24 Bpd"), c("24 hpi", "24 Trp-")),
annotation=c("ns", "ns", "***", "***"), y_position=c(10, 11, 12, 13)) +
theme(line=element_line(size=1, colour="black"),
panel.border=element_rect(colour="black", fill="NA", size=1)) +
theme(plot.title=element_text(hjust='0.5', face="italic"))
```

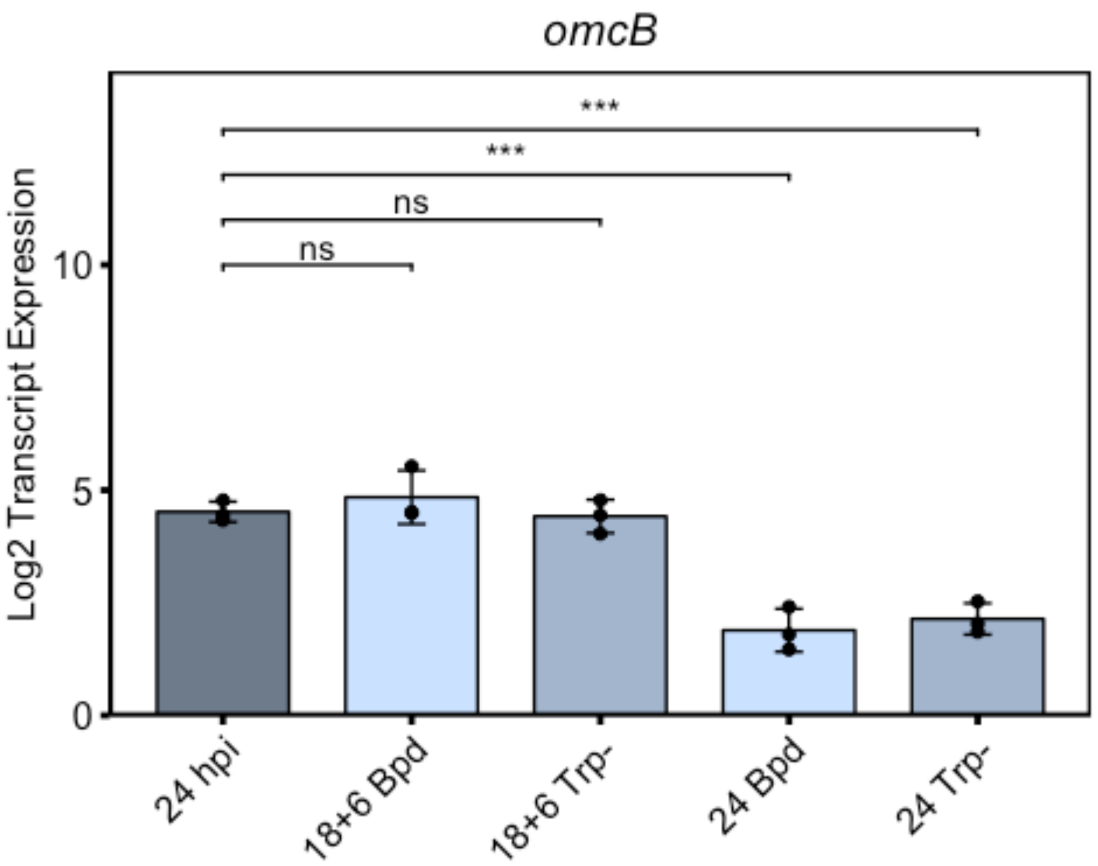


```
euo<-subset(rt, Gene%in%c("euo"))
euoaov<-aov(Log2_Value~ID, data=euo)
summary(euoaov)
```

```
##              Df Sum Sq Mean Sq F value  Pr(>F)
## ID            4  9.869   2.467   11.86 0.00124 **
## Residuals     9  1.872   0.208
## ---
## Signif. codes:  0 '***' 0.001 '**' 0.01 '*' 0.05 '.' 0.1 ' ' 1
```

```
TukeyHSD(x=euoaov)
```

```
##   Tukey multiple comparisons of means
##     95% family-wise confidence level
##
## Fit: aov(formula = Log2_Value ~ ID, data = euo)
##
## $ID
##                           diff        lwr        upr     p adj
## 18+6 Trp--18+6 Bpd  0.2648537 -0.9874213  1.5171286 0.9487041
## 24 Bpd-18+6 Bpd     1.7954179  0.5431430  3.0476928 0.0063579
## 24 hpi-18+6 Bpd    -0.5728762 -1.8251512  0.6793987 0.5655833
## 24 Trp--18+6 Bpd    0.9935047 -0.4065813  2.3935906 0.2035951
## 24 Bpd-18+6 Trp-    1.5305642  0.2782893  2.7828391 0.0169807
```

```
## 24 hpi-18+6 Trp-    -0.8377299 -2.0900048  0.4145450 0.2448921
## 24 Trp--18+6 Trp-    0.7286510 -0.6714349  2.1287369 0.4533450
## 24 hpi-24 Bpd       -2.3682941 -3.6205690 -1.1160192 0.0009306
## 24 Trp--24 Bpd      -0.8019132 -2.2019991  0.5981727 0.3696696
## 24 Trp--24 hpi       1.5663809  0.1662950  2.9664668 0.0279517
```

```
ggbarplot(euo, x="ID2", y="Log2_Value", add="mean_sd", fill="Treatment2",
palette=c("lightsteelblue4", "lightsteelblue1", "lightsteelblue3"),
title="euo") + scale_y_continuous(expand=expand_scale(mult=c(0,0.1))) +
ylab(label="Log2 Transcript Expression") + rremove("legend") +
theme(axis.text.x=element_text(angle=45, hjust=1)) + rremove("xlab") +
geom_exec(geomfunc=geom_point, data=euo, x="ID2", y="Log2_Value",
colour="black") + geom_signif(comparisons=list(c("24 hpi", "18+6 Bpd"), c("24
hpi", "18+6 Trp-"), c("24 hpi", "24 Bpd"), c("24 hpi", "24 Trp-")),
annotation=c("ns", "ns", "***", "*"), y_position=c(10, 11, 12, 13)) +
theme(line=element_line(size=1, colour="black"),
panel.border=element_rect(colour="black", fill="NA", size=1)) +
theme(plot.title=element_text(hjust='0.5', face="italic"))
```

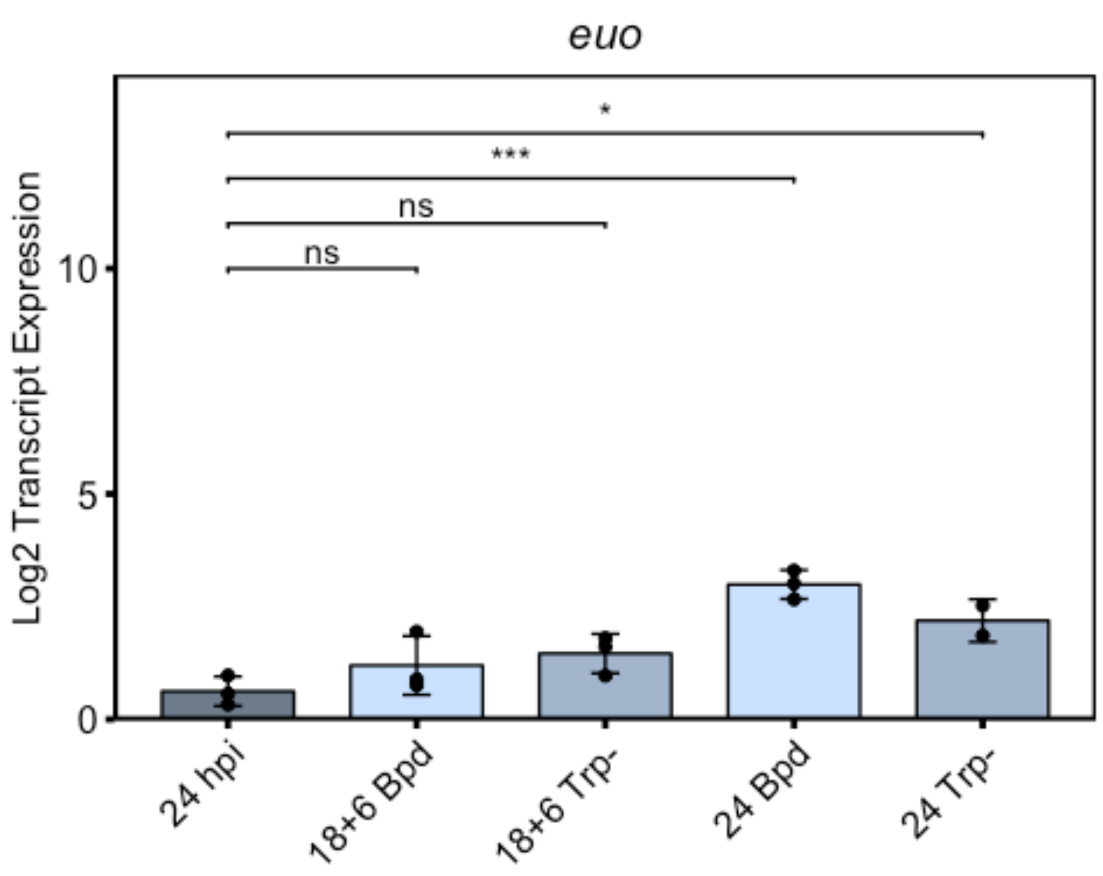


```
#Fig 1d
setwd("~/Documents/Carabeo Lab/R Code/Paper 2/Figure 1")
library(ggplot2)
library(ggpubr)
```

```
rt<-read.csv("Fig1d.csv")
rt$Treatment2<-factor(rt$Treatment, levels=c("Mock", "Bpd", "Trp"))
rt$ID2<-factor(rt$ID, levels=c("24 hpi", "18+6 Bpd", "18+6 Trp-", "24 Bpd",
"24 Trp-"))
trpR<-subset(rt, Gene%in%c("trpR"))
trpRaov<-aov(Log2_Value~ID, data=trpR)
summary(trpRaov)
```

```
##             Df Sum Sq Mean Sq F value   Pr(>F)    
## ID           4 129.78   32.45   83.58 1.21e-07 ***
## Residuals   10   3.88    0.39                     
## ---
## Signif. codes:  0 '***' 0.001 '**' 0.01 '*' 0.05 '.' 0.1 ' ' 1
```

```
TukeyHSD(x=trpRaov)
```

```
##   Tukey multiple comparisons of means
##     95% family-wise confidence level
## 
## Fit: aov(formula = Log2_Value ~ ID, data = trpR)
## 
## $ID
##                           diff       lwr        upr     p adj
## 18+6 Trp--18+6 Bpd  5.5735426  3.899239  7.2478460 0.0000053
## 24 Bpd-18+6 Bpd     2.7807179  1.106414  4.4550214 0.0019772
## 24 hpi-18+6 Bpd    -0.7725651 -2.446869  0.9017384 0.5741283
## 24 Trp--18+6 Bpd    6.6700711  4.995768  8.3443746 0.0000010
## 24 Bpd-18+6 Trp-   -2.7928247 -4.467128 -1.1185212 0.0019138
## 24 hpi-18+6 Trp-   -6.3461077 -8.020411 -4.6718042 0.0000016
## 24 Trp--18+6 Trp-   1.0965285 -0.577775  2.7708320 0.2703491
## 24 hpi-24 Bpd      -3.5532830 -5.227586 -1.8789795 0.0002833
## 24 Trp--24 Bpd      3.8893532  2.215050  5.5636567 0.0001322
## 24 Trp--24 hpi      7.4426362  5.768333  9.1169397 0.0000004
```

```
ggbarplot(trpR, x="ID2", y="Log2_Value", add="mean_sd", fill="Treatment2",
palette=c("lightsteelblue4", "lightsteelblue1", "lightsteelblue3"),
title="trpR") + scale_y_continuous(expand=expand_scale(mult=c(0,0.1))) +
ylab(label="Log2 Transcript Expression") + rremove("legend") +
theme(axis.text.x=element_text(angle=45, hjust=1)) + rremove("xlab") +
geom_exec(geomfunc=geom_point, data=trpR, x="ID2", y="Log2_Value",
colour="black") + geom_signif(comparisons=list(c("24 hpi", "18+6 Bpd"), c("24
hpi", "18+6 Trp-"), c("24 hpi", "24 Bpd"), c("24 hpi", "24 Trp-"), c("18+6
Bpd", "18+6 Trp-"), c("24 Bpd", "24 Trp-")), annotation=c("ns", "***", "***",
"***", "***", "***"), y_position=c(5, 11, 12, 13, 9.5, 10.5)) +
theme(line=element_line(size=1, colour="black"),
panel.border=element_rect(colour="black", fill="NA", size=1)) +
theme(plot.title=element_text(hjust='0.5', face="italic"))
```

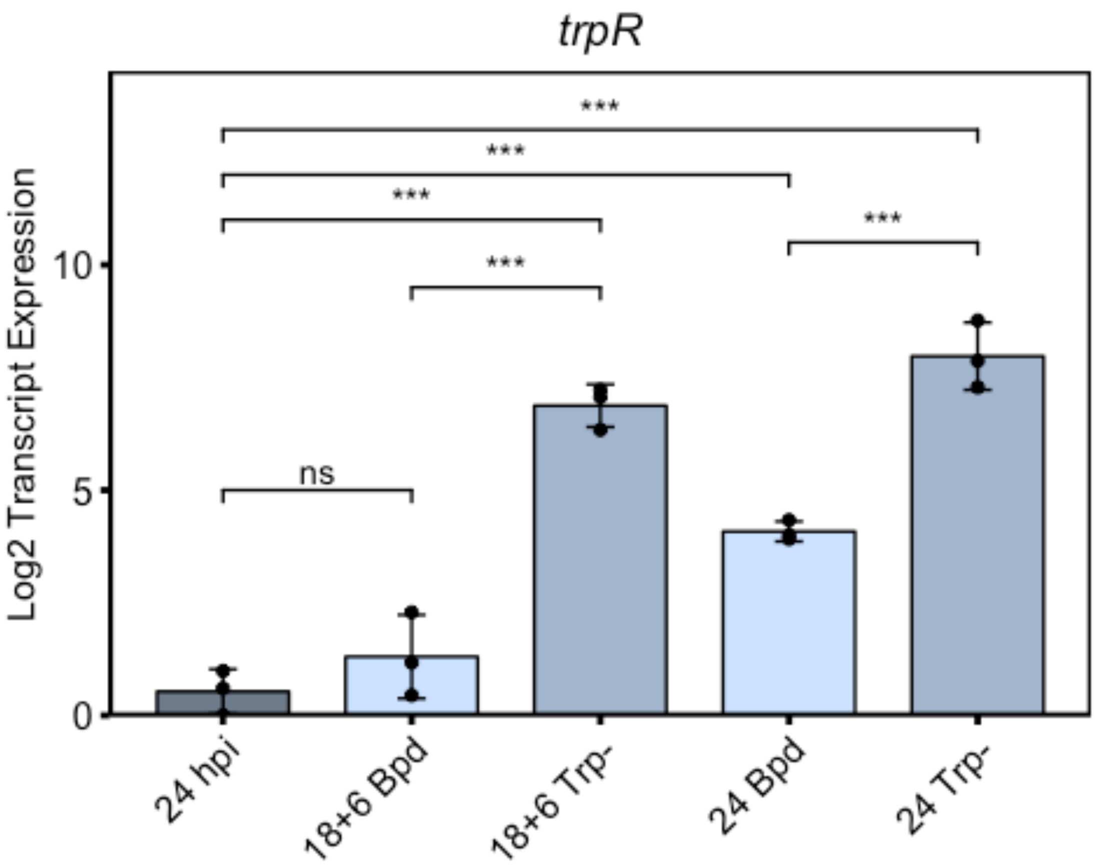


```r
trpB<-subset(rt, Gene%in%c("trpB"))
trpBaov<-aov(Log2_Value~ID, data=trpB)
summary(trpBaov)
```

```
##              Df Sum Sq Mean Sq F value   Pr(>F)    
## ID            4 161.80   40.45   184.5 2.53e-09 ***
## Residuals    10   2.19    0.22                     
## ---
## Signif. codes:  0 '***' 0.001 '**' 0.01 '*' 0.05 '.' 0.1 ' ' 1
```

```r
TukeyHSD(x=trpBaov)
```

```
##   Tukey multiple comparisons of means
##     95% family-wise confidence level
## 
## Fit: aov(formula = Log2_Value ~ ID, data = trpB)
## 
## $ID
##                        diff        lwr        upr     p adj
## 18+6 Trp--18+6 Bpd  5.463509  4.2054354  6.7215830 0.0000004
## 24 Bpd-18+6 Bpd     2.805007  1.5469330  4.0630805 0.0001873
## 24 hpi-18+6 Bpd    -2.247987 -3.5060611 -0.9899136 0.0011313
## 24 Trp--18+6 Bpd    6.532982  5.2749084  7.7910560 0.0000001
## 24 Bpd-18+6 Trp-   -2.658502 -3.9165762 -1.4004287 0.0002936
```

```
## 24 hpi-18+6 Trp-   -7.711497 -8.9695704 -6.4534228 0.0000000
## 24 Trp--18+6 Trp-   1.069473 -0.1886008  2.3275467 0.1068194
## 24 hpi-24 Bpd      -5.052994 -6.3110679 -3.7949203 0.0000009
## 24 Trp--24 Bpd      3.727975  2.4699016  4.9860492 0.0000154
## 24 Trp--24 hpi      8.780970  7.5228958 10.0390433 0.0000000
```

```
ggbarplot(trpB, x="ID2", y="Log2_Value", add="mean_sd", fill="Treatment2",
palette=c("lightsteelblue4", "lightsteelblue1", "lightsteelblue3"),
title="trpB") + scale_y_continuous(expand=expand_scale(mult=c(0,0.1))) +
ylab(label="Log2 Transcript Expression") + rremove("legend") +
theme(axis.text.x=element_text(angle=45, hjust=1)) + rremove("xlab") +
geom_exec(geomfunc=geom_point, data=trpB, x="ID2", y="Log2_Value",
colour="black") + geom_signif(comparisons=list(c("24 hpi", "18+6 Bpd"), c("24
hpi", "18+6 Trp-"), c("24 hpi", "24 Bpd"), c("24 hpi", "24 Trp-"), c("18+6
Bpd", "18+6 Trp-"), c("24 Bpd", "24 Trp-")), annotation=c("**", "***", "***",
"***", "***", "***"), y_position=c(5, 11, 12, 13, 9.5, 10.5)) +
theme(line=element_line(size=1, colour="black"),
panel.border=element_rect(colour="black", fill="NA", size=1)) +
theme(plot.title=element_text(hjust='0.5', face="italic"))
```

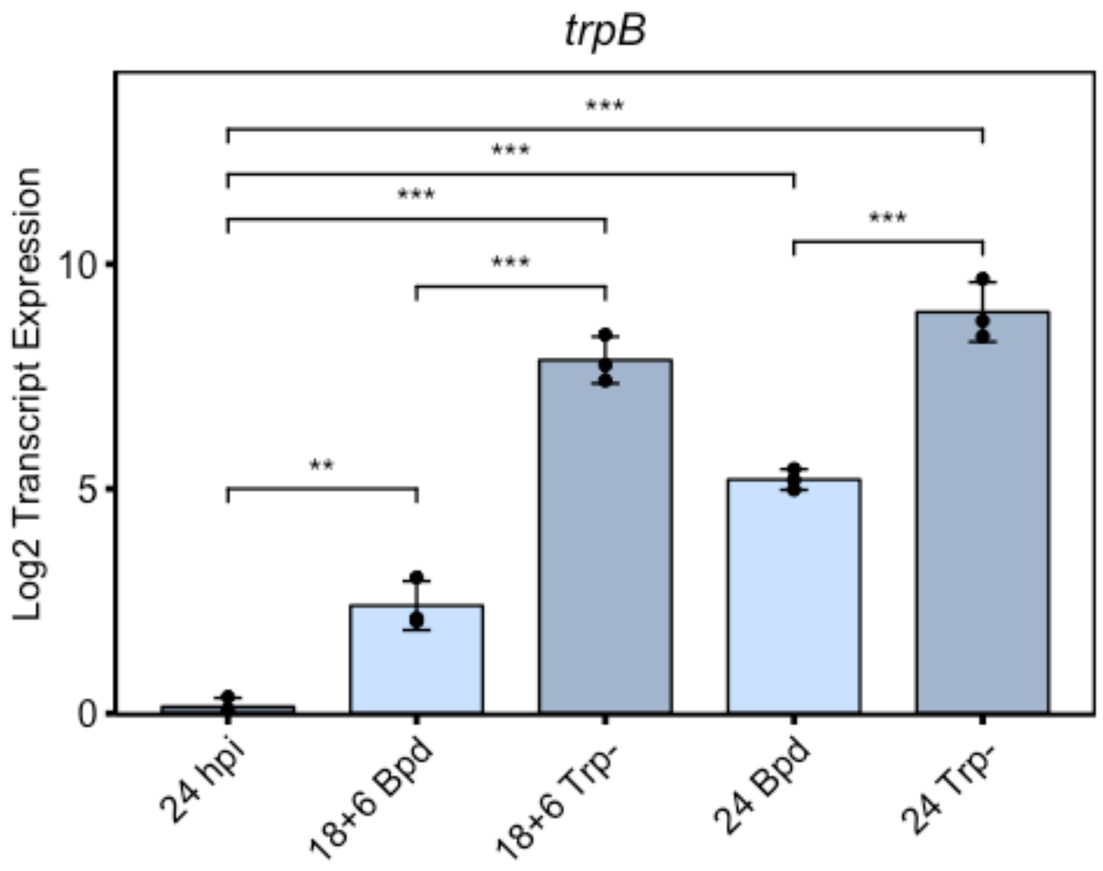


```
trpA<-subset(rt, Gene%in%c("trpA"))
trpAaov<-aov(Log2_Value~ID, data=trpA)
summary(trpAaov)
```

```
##              Df Sum Sq Mean Sq F value   Pr(>F)
## ID            4 137.12   34.28   242.2 6.62e-10 ***
## Residuals    10   1.42    0.14
## ---
## Signif. codes:  0 '***' 0.001 '**' 0.01 '*' 0.05 '.' 0.1 ' ' 1
```

```
TukeyHSD(x=trpAaov)
```

```
##   Tukey multiple comparisons of means
##     95% family-wise confidence level
##
## Fit: aov(formula = Log2_Value ~ ID, data = trpA)
##
## $ID
##                             diff        lwr        upr     p adj
## 18+6 Trp--18+6 Bpd  5.034743  4.0237649  6.0457202 0.0000001
## 24 Bpd-18+6 Bpd     2.924907  1.9139291  3.9358844 0.0000192
## 24 hpi-18+6 Bpd    -1.829587 -2.8405650 -0.8186097 0.0010246
## 24 Trp--18+6 Bpd    6.256004  5.2450268  7.2669821 0.0000000
## 24 Bpd-18+6 Trp-   -2.109836 -3.1208135 -1.0988581 0.0003256
## 24 hpi-18+6 Trp-   -6.864330 -7.8753076 -5.8533522 0.0000000
## 24 Trp--18+6 Trp-   1.221262  0.2102842  2.2322395 0.0173451
## 24 hpi-24 Bpd      -4.754494 -5.7654718 -3.7435165 0.0000002
## 24 Trp--24 Bpd      3.331098  2.3201200  4.3420753 0.0000058
## 24 Trp--24 hpi      8.085592  7.0746141  9.0965694 0.0000000
```

```
ggbarplot(trpA, x="ID2", y="Log2_Value", add="mean_sd", fill="Treatment2",
palette=c("lightsteelblue4", "lightsteelblue1", "lightsteelblue3"),
title="trpA") + scale_y_continuous(expand=expand_scale(mult=c(0,0.1))) +
ylab(label="Log2 Transcript Expression") + rremove("legend") +
theme(axis.text.x=element_text(angle=45, hjust=1)) + rremove("xlab") +
geom_exec(geomfunc=geom_point, data=trpA, x="ID2", y="Log2_Value",
colour="black") + geom_signif(comparisons=list(c("24 hpi", "18+6 Bpd"), c("24
hpi", "18+6 Trp-"), c("24 hpi", "24 Bpd"), c("24 hpi", "24 Trp-"), c("18+6
Bpd", "18+6 Trp-"), c("24 Bpd", "24 Trp-")), annotation=c("***", "***",
"***", "***", "***", "***"), y_position=c(5, 11, 12, 13, 9.5, 10.5)) +
theme(line=element_line(size=1, colour="black"),
panel.border=element_rect(colour="black", fill="NA", size=1)) +
theme(plot.title=element_text(hjust='0.5', face="italic"))
```

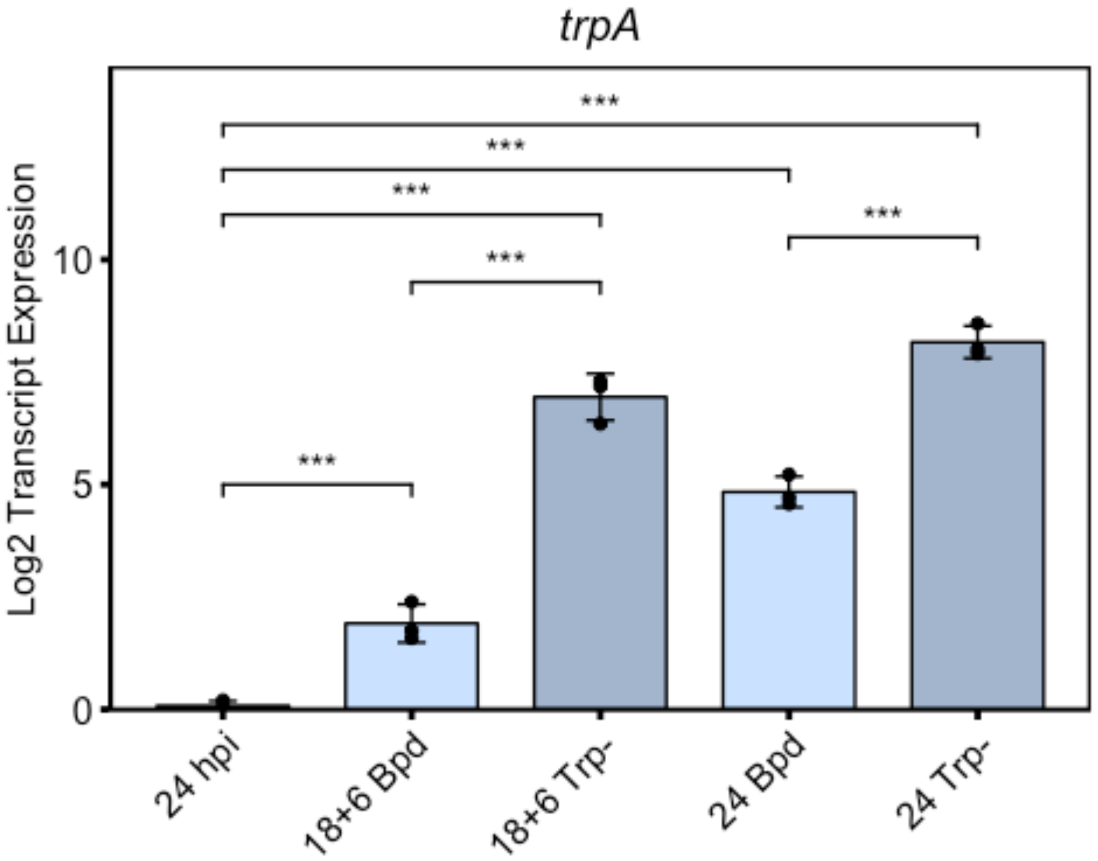


```
#Fig 1e
setwd("~/Documents/Carabeo Lab/R Code/Paper 2/Figure 1")
library(ggplot2)
library(ggpubr)
rt<-read.csv("Fig1e.csv")
rt$ID2<-factor(rt$ID, levels=c("24 hpi", "18+6 Bpd", "18+6 Trp-", "18+6 Bpd +
Trp-"))
rt$Treatment2<-factor(rt$Treatment, levels=c("Mock", "Bpd", "Trp", "Bpd +
Trp"))
trpB<-subset(rt, Gene%in%c("trpB"))
trpBaov<-aov(Log2_Value~ID, data=trpB)
summary(trpBaov)

##             Df Sum Sq Mean Sq F value   Pr(>F)
## ID           3 137.70   45.90   96.91 1.25e-06 ***
## Residuals    8   3.79    0.47
## ---
## Signif. codes:  0 '***' 0.001 '**' 0.01 '*' 0.05 '.' 0.1 ' ' 1

TukeyHSD(x=trpBaov)

##   Tukey multiple comparisons of means
##     95% family-wise confidence level
##
```

```
## Fit: aov(formula = Log2_Value ~ ID, data = trpB)
##
## $ID
##                                  diff        lwr       upr     p adj
## 18+6 Bpd + Trp--18+6 Bpd    4.0587587   2.259271  5.858247 0.0004136
## 18+6 Trp--18+6 Bpd          4.5597172   2.760229  6.359205 0.0001818
## 24 hpi-18+6 Bpd            -3.7999644  -5.599453 -2.000476 0.0006517
## 18+6 Trp--18+6 Bpd + Trp-   0.5009585  -1.298530  2.300447 0.8095902
## 24 hpi-18+6 Bpd + Trp-     -7.8587231  -9.658211 -6.059235 0.0000032
## 24 hpi-18+6 Trp-           -8.3596816 -10.159170 -6.560193 0.0000020
```

```
ggbarplot(trpB, x="ID2", y="Log2_Value", add="mean_sd", fill="Treatment2",
palette=c("darkseagreen4", "darkseagreen", "darkseagreen3", "darkseagreen1"),
title="trpB") + scale_y_continuous(expand=expand_scale(mult=c(0,0.1))) +
ylab(label="Log2 Transcript Expression") + rremove("legend") +
theme(axis.text.x=element_text(angle=45, hjust=1)) + rremove("xlab") +
geom_exec(geomfunc=geom_point, data=trpB, x="ID2", y="Log2_Value",
colour="black") + geom_signif(comparisons=list(c("18+6 Trp-", "18+6 Bpd +
Trp-"), c("24 hpi", "18+6 Bpd"), c("24 hpi", "18+6 Trp-"), c("24 hpi", "18+6
Bpd + Trp-")), annotation=c("ns", "***", "***", "***"), y_position=c(10, 11,
12, 13)) + theme(line=element_line(size=1, colour="black"),
panel.border=element_rect(colour="black", fill="NA", size=1)) +
theme(plot.title=element_text(hjust='0.5', face="italic"))
```

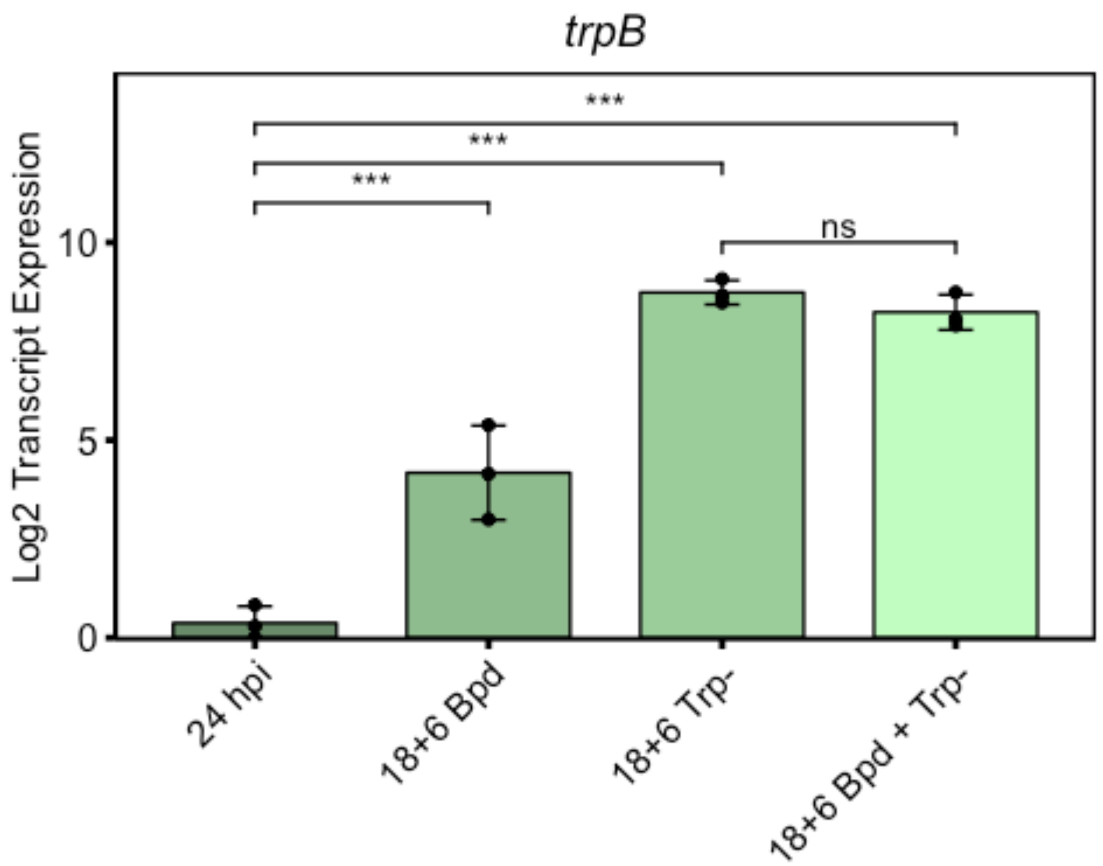

```
trpA<-subset(rt, Gene%in%c("trpA"))
trpAaov<-aov(Log2_Value~ID, data=trpA)
summary(trpAaov)
```

```
##             Df Sum Sq Mean Sq F value   Pr(>F)    
## ID           3 122.47   40.82   62.67 6.72e-06 ***
## Residuals    8   5.21    0.65                     
## ---
## Signif. codes:  0 '***' 0.001 '**' 0.01 '*' 0.05 '.' 0.1 ' ' 1
```

```
TukeyHSD(x=trpAaov)
```

```
##   Tukey multiple comparisons of means
##     95% family-wise confidence level
## 
## Fit: aov(formula = Log2_Value ~ ID, data = trpA)
## 
## $ID
##                                diff        lwr       upr     p adj
## 18+6 Bpd + Trp--18+6 Bpd   3.4860728   1.375730  5.596416 0.0032648
## 18+6 Trp--18+6 Bpd         4.1035767   1.993233  6.213920 0.0011373
## 24 hpi-18+6 Bpd           -3.9026910  -6.013034 -1.792348 0.0015844
## 18+6 Trp--18+6 Bpd + Trp-  0.6175038  -1.492839  2.727847 0.7867219
## 24 hpi-18+6 Bpd + Trp-    -7.3887638  -9.499107 -5.278421 0.0000168
## 24 hpi-18+6 Trp-          -8.0062677 -10.116611 -5.895924 0.0000092
```

```
ggbarplot(trpA, x="ID2", y="Log2_Value", add="mean_sd", fill="Treatment2",
palette=c("darkseagreen4", "darkseagreen", "darkseagreen3", "darkseagreen1"),
title="trpA") + scale_y_continuous(expand=expand_scale(mult=c(0,0.1))) +
ylab(label="Log2 Transcript Expression") + rremove("legend") +
theme(axis.text.x=element_text(angle=45, hjust=1)) + rremove("xlab") +
geom_exec(geomfunc=geom_point, data=trpA, x="ID2", y="Log2_Value",
colour="black") + geom_signif(comparisons=list(c("18+6 Trp-", "18+6 Bpd +
Trp-"), c("24 hpi", "18+6 Bpd"), c("24 hpi", "18+6 Trp-"), c("24 hpi", "18+6
Bpd + Trp-")), annotation=c("ns", "**", "***", "***"), y_position=c(10, 11,
12, 13)) + theme(line=element_line(size=1, colour="black"),
panel.border=element_rect(colour="black", fill="NA", size=1)) +
theme(plot.title=element_text(hjust='0.5', face="italic"))
```

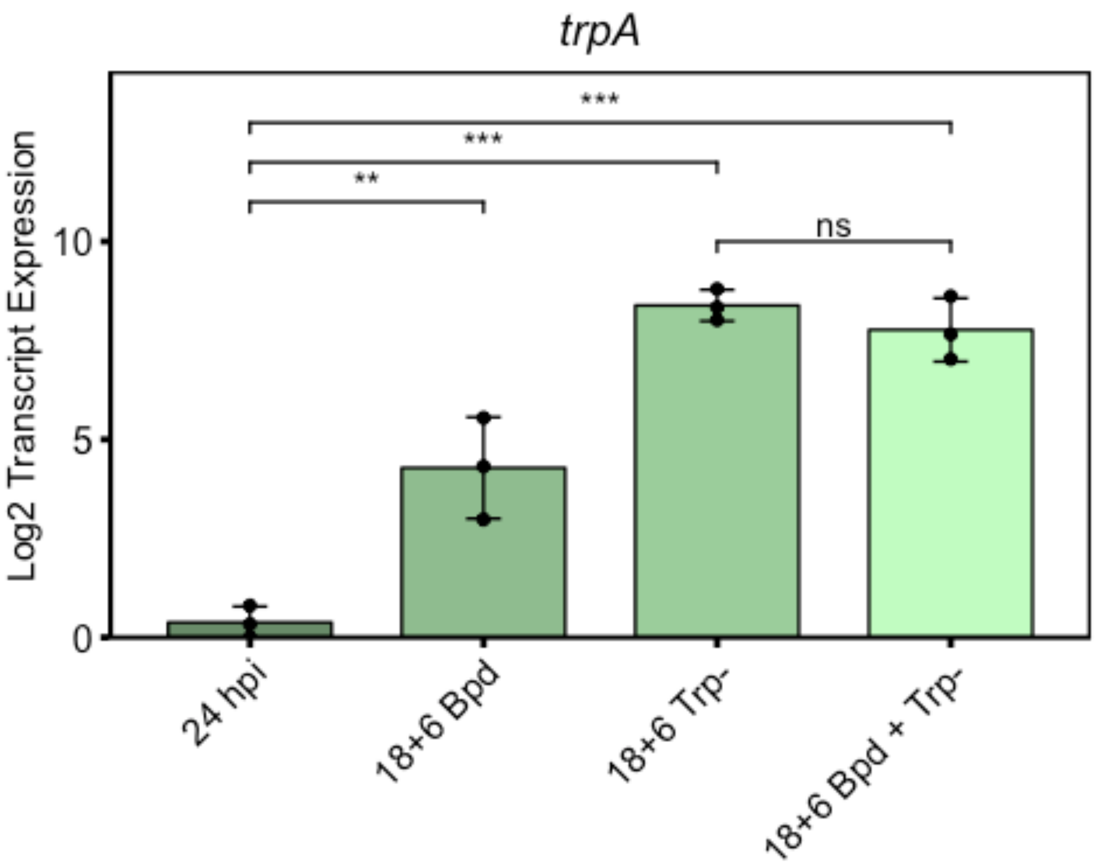

trpA
Log2 Transcript Expression
0
5
10
***
***
**
ns
24 hpi
18+6 Bpd
18+6 Trp-
18+6 Bpd + Trp-

### Figure 2

Nick Pokorzynski

4/15/2020

```
#Fig2e
setwd("~/Documents/Carabeo Lab/R Code/Paper 2/Figure 2")
library(ggplot2)
library(ggpubr)

## Warning: package 'ggpubr' was built under R version 3.5.2

## Loading required package: magrittr

sum<-read.csv("Fig2e.csv")
www<-subset(sum, ID%in%c("WWW"))
wilcox.test(Mean~Treatment, data=www)

##
##  Wilcoxon rank sum test
##
## data:  Mean by Treatment
## W = 643, p-value = 0.004114
## alternative hypothesis: true location shift is not equal to 0

yyf<-subset(sum, ID%in%c("YYF"))
wilcox.test(Mean~Treatment, data=yyf)

##
##  Wilcoxon rank sum test
##
## data:  Mean by Treatment
## W = 774, p-value = 0.05495
## alternative hypothesis: true location shift is not equal to 0

sum$Treatment2<-factor(sum$Treatment, levels=c("Trp+", "Trp-"))
ggviolin(sum, x="Treatment2", y="Mean", add='median', facet.by='ID',
fill="Treatment2", palette=c("dodgerblue4", "dodgerblue1")) +
ylab(label="MFI") + rremove('xlab') +
stat_compare_means(comparisons=list(c("Trp+", "Trp-")), label="p.signif",
method="wilcox.test", label.y=1800) +
scale_y_continuous(expand=expand_scale(mult=c(0.1,0.1))) + rremove('legend')
```

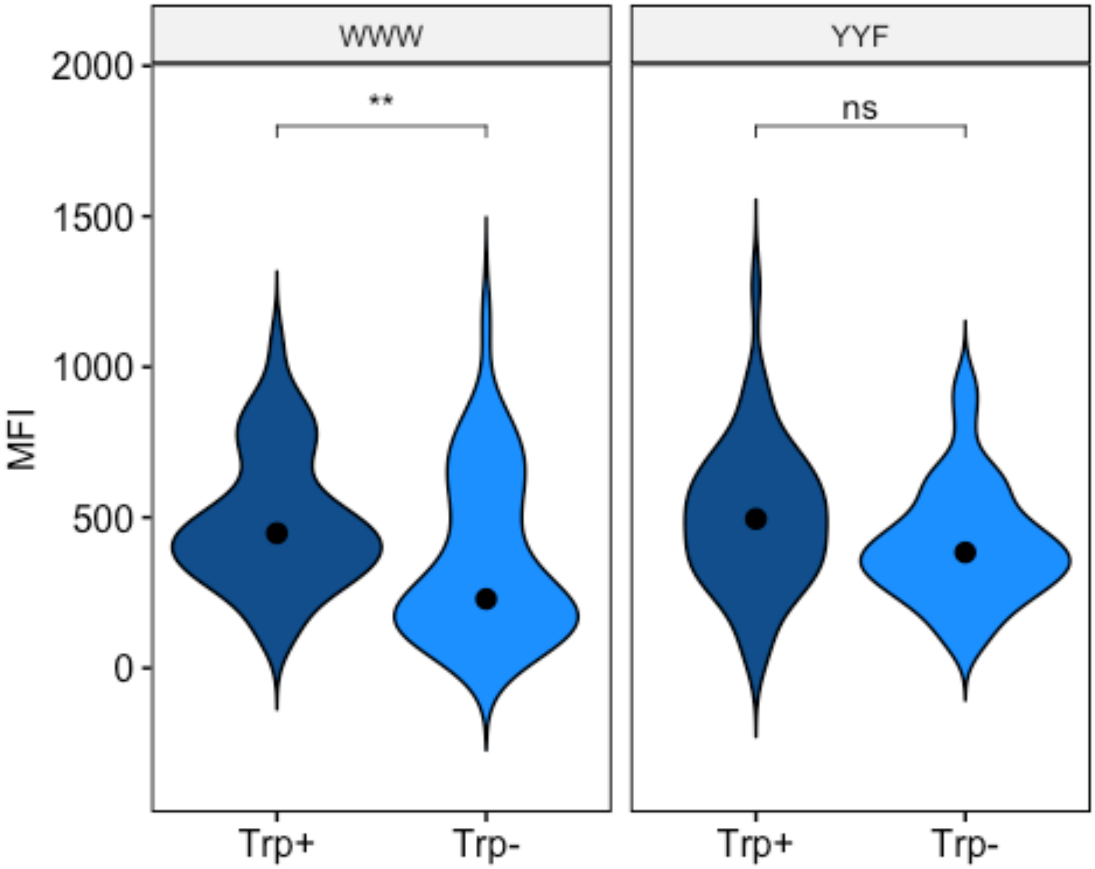
WWW
YYF
**
ns
2000
1500
1000
500
0
MFI
Trp+
Trp-
Trp+
Trp-

### Figure 3

Nick Pokorzynski

4/15/2020

```
#Fig3b
setwd("~/Documents/Carabeo Lab/R Code/Paper 2/Figure 3")
library(ggplot2)
library(ggpubr)

## Warning: package 'ggpubr' was built under R version 3.5.2

## Loading required package: magrittr

fig3<-read.csv("Fig3b,d.csv")
an<-subset(fig3, Treatment%in%c("AN3365"))
t.test(Value~ID, data=an)

##
##  Welch Two Sample t-test
##
## data:  Value by ID
## t = 1.0952, df = 3.6107, p-value = 0.3411
## alternative hypothesis: true difference in means is not equal to 0
## 95 percent confidence interval:
##  -45.58437 100.94842
## sample estimates:
## mean in group WWW mean in group YYF
##         100.00000          72.31797

ggbarplot(an, x="ID", y="Value", add="mean_sd", fill="ID",
palette=c("orchid4", "orchid1"), title="AN3365") +
scale_y_continuous(expand=expand_scale(mult=c(0,0.1))) +
ylab(label="Normalized Inhibition (%)") + rremove("legend") +
theme(axis.text.x=element_text(angle=45, hjust=1)) + rremove("xlab") +
geom_exec(geomfunc=geom_point, data=an, x="ID", y="Value", colour="black") +
geom_signif(comparisons=list(c("WWW", "YYF")), annotation="ns",
y_position=150) + theme(line=element_line(size=1, colour="black"),
panel.border=element_rect(colour="black", fill="NA", size=1)) +
theme(plot.title=element_text(hjust='0.5'))
```

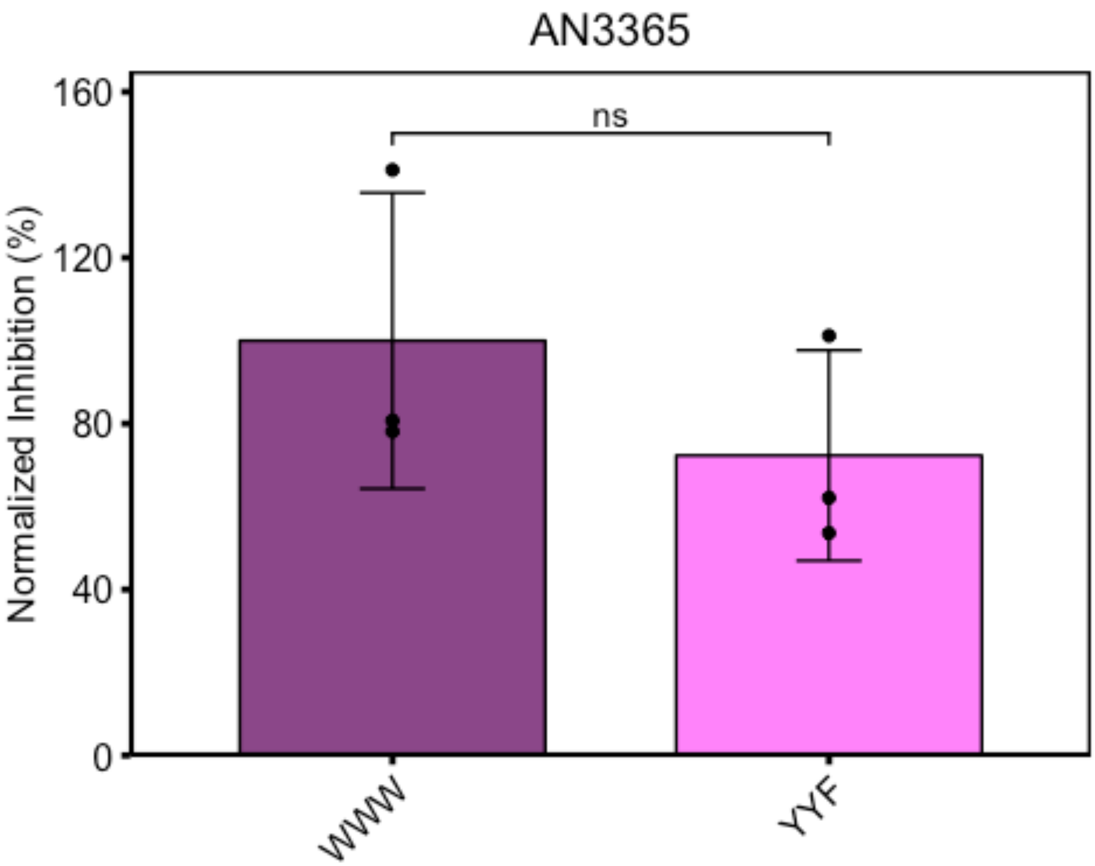


```r
#Fig3d
ind<-subset(fig3, Treatment%in%c("Indolmycin"))
t.test(Value~ID, data=ind)

##
##  Welch Two Sample t-test
##
## data:  Value by ID
## t = 9.1152, df = 5.1416, p-value = 0.0002297
## alternative hypothesis: true difference in means is not equal to 0
## 95 percent confidence interval:
##  48.58699 86.31901
## sample estimates:
## mean in group WWW mean in group YYF
##           100.000            32.547

ggbarplot(ind, x="ID", y="Value", add="mean_sd", fill="ID",
palette=c("mediumpurple4", "mediumpurple1"), title="Indolmycin") +
scale_y_continuous(expand=expand_scale(mult=c(0,0.1))) +
ylab(label="Normalized Inhibition (%)") + rremove("legend") +
theme(axis.text.x=element_text(angle=45, hjust=1)) + rremove("xlab") +
geom_exec(geomfunc=geom_point, data=ind, x="ID", y="Value", colour="black") +
geom_signif(comparisons=list(c("WWW", "YYF")), annotation="***",
```

```
y_position=150) + theme(line=element_line(size=1, colour="black"),
panel.border=element_rect(colour="black", fill="NA", size=1)) +
theme(plot.title=element_text(hjust='0.5'))
```

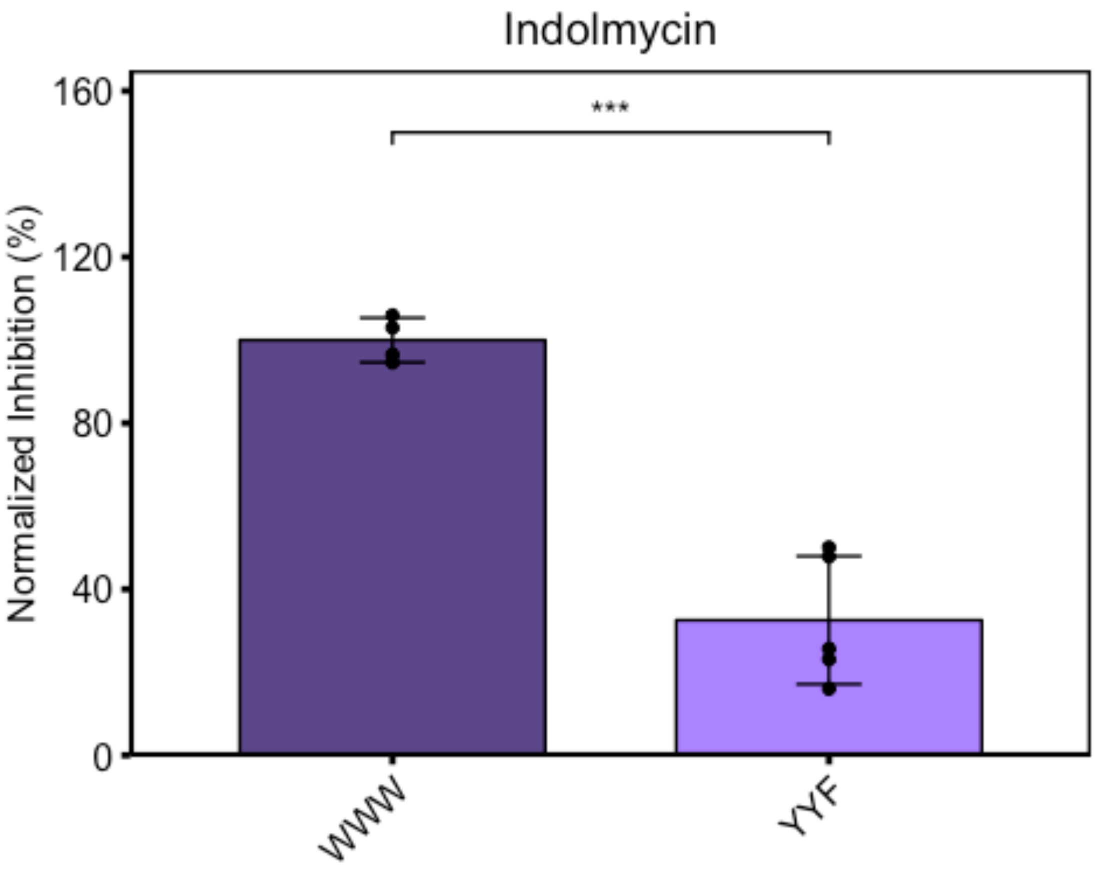

### Figure 4

Nick Pokorzynski

4/16/2020

```
#Fig4b
setwd("~/Documents/Carabeo Lab/R Code/Paper 2/Figure 4")
library(ggplot2)
library(ggpubr)

## Warning: package 'ggpubr' was built under R version 3.5.2

## Loading required package: magrittr

x<-read.csv("Fig4b.csv")
x$Treatment2<-factor(x$Treatment, levels=c("UTD", "Ind", "IFNg"))
#subset values for each gene for statistical analysis
CTL0174<-subset(x, Gene%in%c("CTL0174"))
GroEL_1<-subset(x, Gene%in%c("GroEL_1"))
OmpA<-subset(x, Gene%in%c("OmpA"))
YtgCR<-subset(x, Gene%in%c("YtgCR"))
#assign analysis of variance to variable
ctlaov<-aov(Value~Treatment, data=CTL0174)
#summarize results of aov
summary(ctlaov)

##             Df Sum Sq Mean Sq F value Pr(>F)
## Treatment    2 0.2265  0.1132   0.781  0.499
## Residuals    6 0.8694  0.1449

#compute Tukey's Honestly Significant Differences multiple comparisons test
TukeyHSD(x=ctlaov)

##   Tukey multiple comparisons of means
##     95% family-wise confidence level
##
## Fit: aov(formula = Value ~ Treatment, data = CTL0174)
##
## $Treatment
##                diff        lwr       upr     p adj
## Ind-IFNg  0.2611761 -0.6924453 1.2147975 0.6938791
## UTD-IFNg -0.1185586 -1.0721800 0.8350629 0.9239101
## UTD-Ind  -0.3797347 -1.3333561 0.5738868 0.4842284

#repeat for each gene
groaov<-aov(Value~Treatment, data=GroEL_1)
summary(groaov)
```

```
##              Df Sum Sq Mean Sq F value Pr(>F)
## Treatment     2 0.2181 0.10903   4.515 0.0636 .
## Residuals     6 0.1449 0.02415
## ---
## Signif. codes:  0 '***' 0.001 '**' 0.01 '*' 0.05 '.' 0.1 ' ' 1
```

```
TukeyHSD(x=groaov)
```

```
##   Tukey multiple comparisons of means
##     95% family-wise confidence level
##
## Fit: aov(formula = Value ~ Treatment, data = GroEL_1)
##
## $Treatment
##                  diff        lwr        upr     p adj
## Ind-IFNg  0.03800001 -0.3513097 0.42730977 0.9521470
## UTD-IFNg -0.30955084 -0.6988606 0.07975892 0.1102394
## UTD-Ind  -0.34755085 -0.7368606 0.04175891 0.0753717
```

```
ompaov<-aov(Value~Treatment, data=OmpA)
summary(ompaov)
```

```
##              Df  Sum Sq  Mean Sq F value Pr(>F)
## Treatment     2 0.01127 0.005633   2.916   0.13
## Residuals     6 0.01159 0.001931
```

```
TukeyHSD(x=ompaov)
```

```
##   Tukey multiple comparisons of means
##     95% family-wise confidence level
##
## Fit: aov(formula = Value ~ Treatment, data = OmpA)
##
## $Treatment
##                  diff         lwr       upr     p adj
## Ind-IFNg -0.00453640 -0.11463572 0.1055629 0.9912403
## UTD-IFNg  0.07268089 -0.03741843 0.1827802 0.1868391
## UTD-Ind   0.07721729 -0.03288203 0.1873166 0.1591450
```

```
ytgcaov<-aov(Value~Treatment, data=YtgCR)
summary(ytgcaov)
```

```
##              Df Sum Sq Mean Sq F value  Pr(>F)
## Treatment     2 0.3728 0.18640    25.5 0.00117 **
## Residuals     6 0.0438 0.00731
## ---
## Signif. codes:  0 '***' 0.001 '**' 0.01 '*' 0.05 '.' 0.1 ' ' 1
```

```
TukeyHSD(x=ytgcaov)
```

```
##   Tukey multiple comparisons of means
##     95% family-wise confidence level
```

```
##
## Fit: aov(formula = Value ~ Treatment, data = YtgCR)
##
## $Treatment
##                 diff         lwr        upr     p adj
## Ind-IFNg -0.1831521 -0.39732083 0.03101665 0.0871991
## UTD-IFNg  0.3099715  0.09580276 0.52414024 0.0103810
## UTD-Ind   0.4931236  0.27895485 0.70729232 0.0009831

#Plot each gene individually w/ annotated significance for TukeyHSD
#CTL0174
ggbarplot(CTL0174, x="Treatment2", y="Value", add="mean_sd", color="black",
fill="Treatment2", palette=c("steelblue4", "steelblue3", "steelblue2"),
title="CTL0174") + scale_y_continuous(limits=c(0, 2.2),
expand=expand_scale(mult=c(0,0.1))) + ylab(label="3'-to-5' Ratio") +
rremove("legend") + theme(axis.text.x=element_text(angle=45, hjust=1)) +
rremove("xlab") + geom_exec(geomfunc=geom_point, data=CTL0174,
x="Treatment2", y="Value", colour="black") +
geom_signif(comparisons=list(c("UTD", "Ind"), c("UTD", "IFNg"), c("Ind",
"IFNg")), annotation=c("ns", "ns", "ns"), y_position = c(1.7, 1.9, 2.1)) +
theme(line=element_line(size=1, colour="black"),
panel.border=element_rect(colour="black", fill="NA", size=1)) +
theme(plot.title=element_text(hjust='0.5'))
```

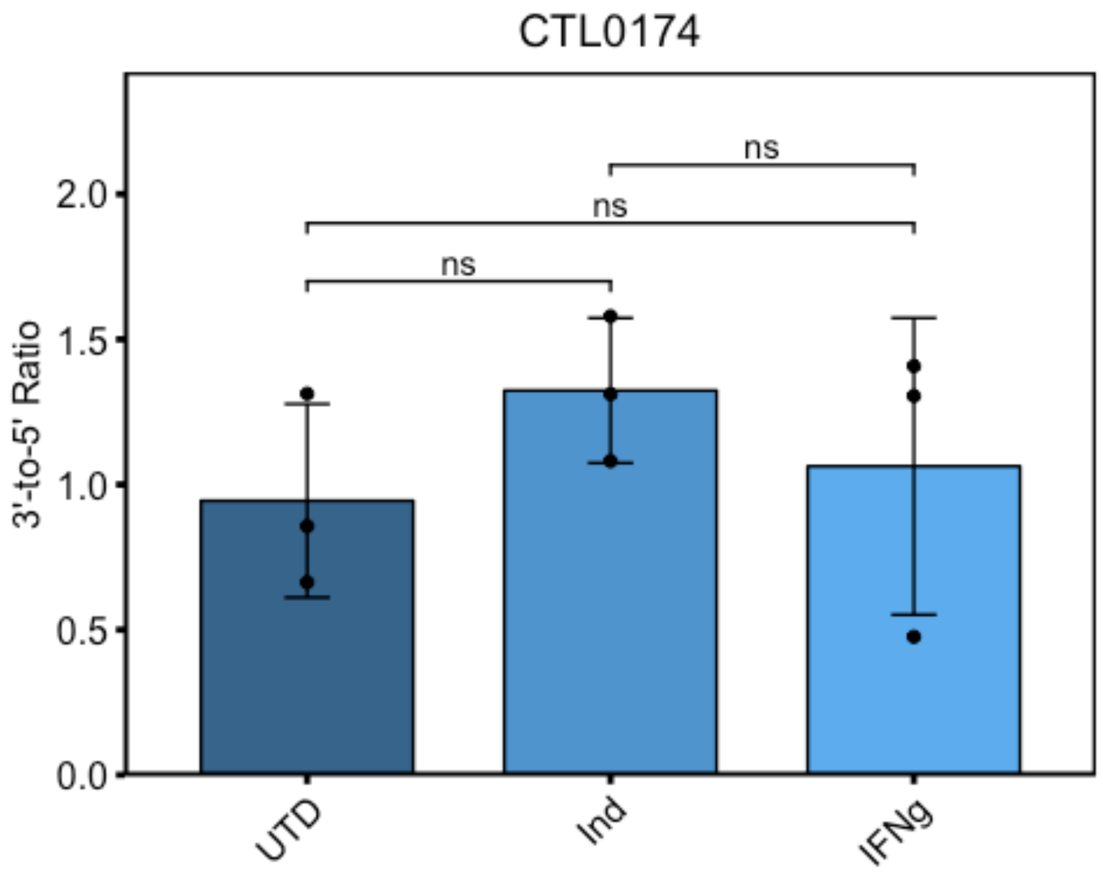

```
#YtgCR
ggbarplot(YtgCR, x="Treatment2", y="Value", add="mean_sd", color="black",
fill="Treatment2", palette=c("steelblue4", "steelblue3", "steelblue2"),
title="YtgCR") + scale_y_continuous(limits=c(0, 2.2),
expand=expand_scale(mult=c(0,0.1))) + ylab(label="3'-to-5' Ratio") +
rremove("legend") + theme(axis.text.x=element_text(angle=45, hjust=1)) +
rremove("xlab") + geom_exec(geomfunc=geom_point, data=YtgCR, x="Treatment2",
y="Value", colour="black") + geom_signif(comparisons=list(c("UTD", "Ind"),
c("UTD", "IFNg"), c("Ind", "IFNg")), annotation=c("***", "*", "ns"),
y_position = c(1.7, 1.9, 2.1)) + theme(line=element_line(size=1,
colour="black"), panel.border=element_rect(colour="black", fill="NA",
size=1)) + theme(plot.title=element_text(hjust='0.5'))
```

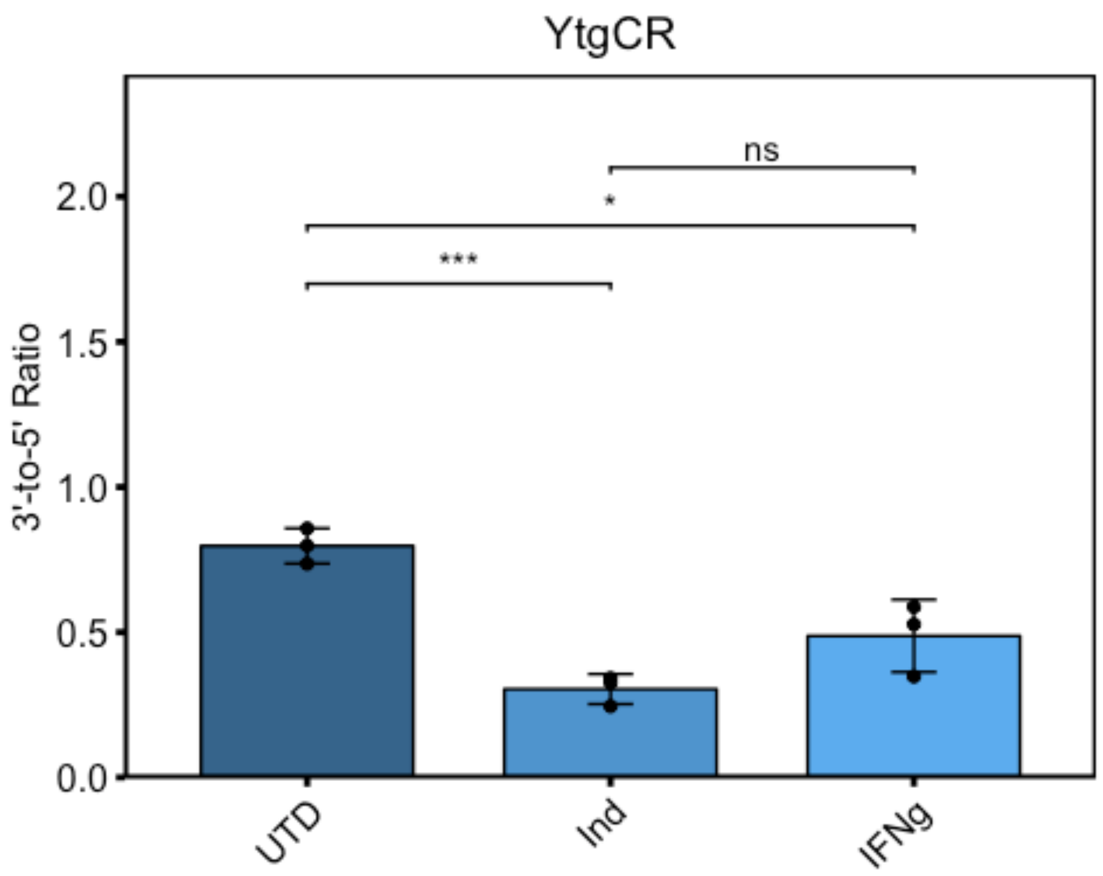


```
#OmpA
ggbarplot(OmpA, x="Treatment2", y="Value", add="mean_sd", color="black",
fill="Treatment2", palette=c("steelblue4", "steelblue3", "steelblue2"),
title="OmpA") + scale_y_continuous(limits=c(0, 2.2),
expand=expand_scale(mult=c(0,0.1))) + ylab(label="3'-to-5' Ratio") +
rremove("legend") + theme(axis.text.x=element_text(angle=45, hjust=1)) +
rremove("xlab") + geom_exec(geomfunc=geom_point, data=OmpA, x="Treatment2",
y="Value", colour="black") + geom_signif(comparisons=list(c("UTD", "Ind"),
c("UTD", "IFNg"), c("Ind", "IFNg")), annotation=c("ns", "ns", "ns"),
y_position = c(1.7, 1.9, 2.1)) + theme(line=element_line(size=1,
```

```
colour="black"), panel.border=element_rect(colour="black", fill="NA",
size=1)) + theme(plot.title=element_text(hjust='0.5'))
```

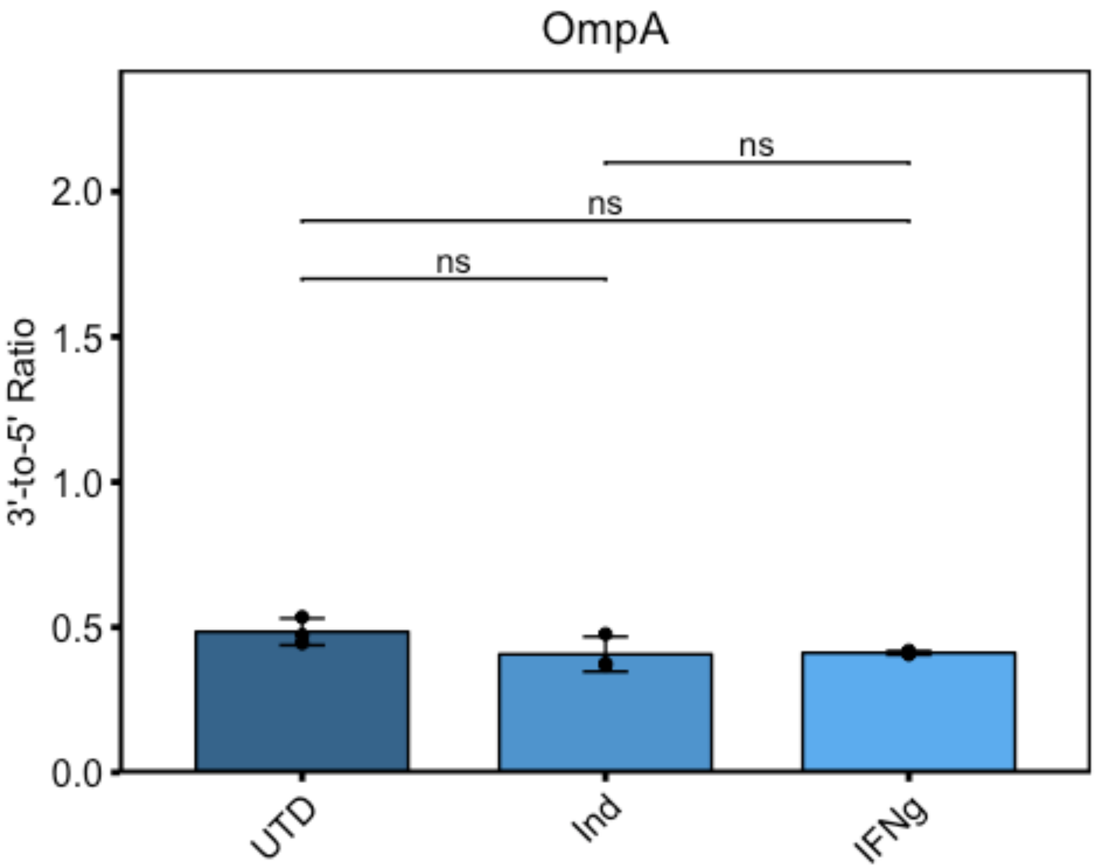


```
#GroEL_1
ggbarplot(GroEL_1, x="Treatment2", y="Value", add="mean_sd", color="black",
fill="Treatment2", palette=c("steelblue4", "steelblue3", "steelblue2"),
title="GroEL_1") + scale_y_continuous(limits=c(0, 2.2),
expand=expand_scale(mult=c(0,0.1))) + ylab(label="3'-to-5' Ratio") +
rremove("legend") + theme(axis.text.x=element_text(angle=45, hjust=1)) +
rremove("xlab") + geom_exec(geomfunc=geom_point, data=GroEL_1,
x="Treatment2", y="Value", colour="black") +
geom_signif(comparisons=list(c("UTD", "Ind"), c("UTD", "IFNg"), c("Ind",
"IFNg")), annotation=c("ns", "ns", "ns"), y_position = c(1.7, 1.9, 2.1)) +
theme(line=element_line(size=1, colour="black"),
panel.border=element_rect(colour="black", fill="NA", size=1)) +
theme(plot.title=element_text(hjust='0.5'))
```

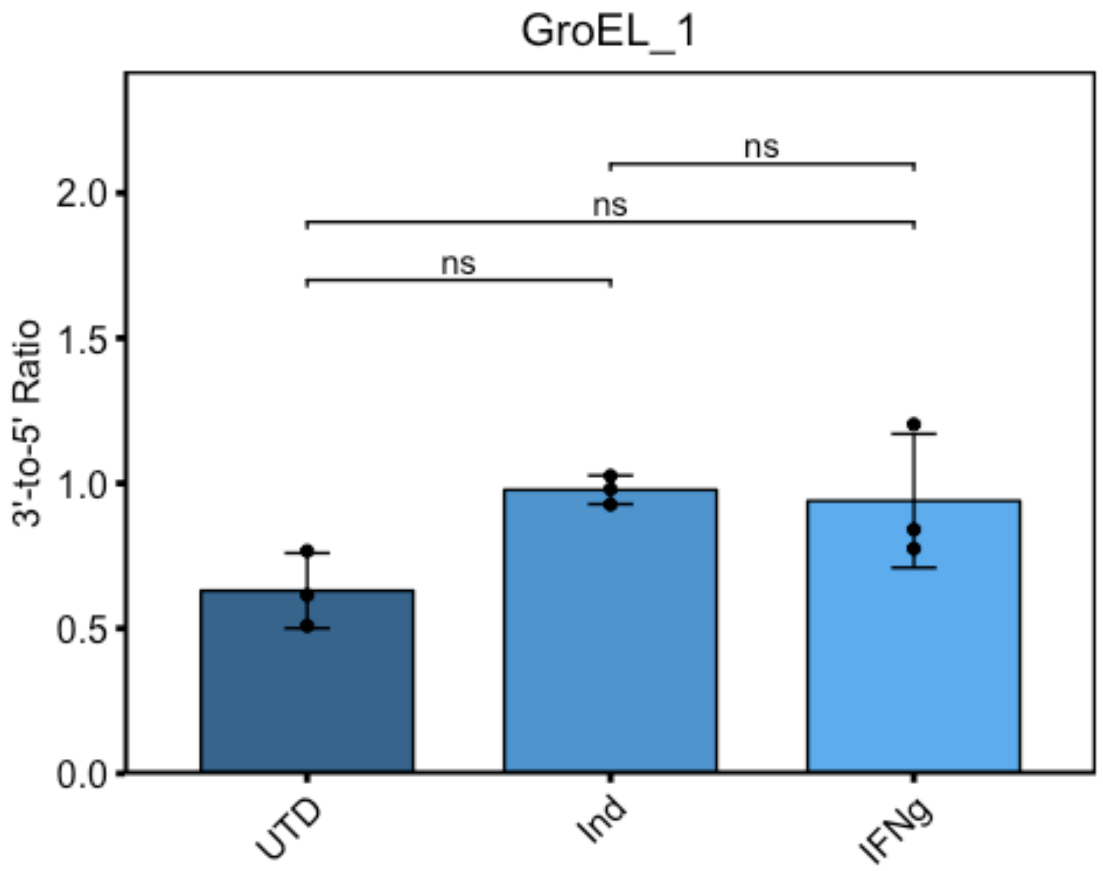


```
#Fig4c
setwd("~/Documents/Carabeo Lab/R Code/Paper 2/Figure 4")
library(ggplot2)
library(ggpubr)
x<-read.csv("Fig4c.csv")
x$Treatment2<-factor(x$Treatment, levels=c("Trp - ", "Ind", "Ind+Bic"))
#subset values for each gene for TukeyHSD analysis
CTL0174<-subset(x, Gene%in%c("CTL0174"))
GroEL_1<-subset(x, Gene%in%c("GroEL_1"))
OmpA<-subset(x, Gene%in%c("OmpA"))
YtgCR<-subset(x, Gene%in%c("YtgCR"))
YtgD<-subset(x, Gene%in%c("YtgD:YtgA"))
#compute analysis of variance and assign to variable
ctlaov<-aov(Value~Treatment, data=CTL0174)
#call aov results
summary(ctlaov)

##             Df Sum Sq Mean Sq F value Pr(>F)
## Treatment    2 0.4795 0.23976   3.244  0.111
## Residuals    6 0.4434 0.07391

#compute Tukey HSD
TukeyHSD(x=ctlaov)
```

```
##   Tukey multiple comparisons of means
##     95% family-wise confidence level
##
## Fit: aov(formula = Value ~ Treatment, data = CTL0174)
##
## $Treatment
##                      diff        lwr       upr     p adj
## Ind+Bic-Ind     0.3254333 -0.3556281 1.0064947 0.3696610
## Trp - -Ind     -0.2377000 -0.9187614 0.4433614 0.5641991
## Trp - -Ind+Bic -0.5631333 -1.2441947 0.1179281 0.0973815
```

```
#repeat for each gene
groaov<-aov(Value~Treatment, data=GroEL_1)
summary(groaov)
```

```
##             Df  Sum Sq Mean Sq F value Pr(>F)
## Treatment    2 0.09841 0.04920   4.999 0.0528 .
## Residuals    6 0.05905 0.00984
## ---
## Signif. codes:  0 '***' 0.001 '**' 0.01 '*' 0.05 '.' 0.1 ' ' 1
```

```
TukeyHSD(x=groaov)
```

```
##   Tukey multiple comparisons of means
##     95% family-wise confidence level
##
## Fit: aov(formula = Value ~ Treatment, data = GroEL_1)
##
## $Treatment
##                      diff         lwr       upr     p adj
## Ind+Bic-Ind    0.20250454 -0.04603820 0.4510473 0.1020872
## Trp - -Ind     0.23707278 -0.01146996 0.4856155 0.0595836
## Trp - -Ind+Bic 0.03456824 -0.21397450 0.2831110 0.9060032
```

```
ompaov<-aov(Value~Treatment, data=OmpA)
summary(ompaov)
```

```
##             Df  Sum Sq Mean Sq F value Pr(>F)
## Treatment    2 0.08267 0.04134   10.26 0.0116 *
## Residuals    6 0.02417 0.00403
## ---
## Signif. codes:  0 '***' 0.001 '**' 0.01 '*' 0.05 '.' 0.1 ' ' 1
```

```
TukeyHSD(x=ompaov)
```

```
##   Tukey multiple comparisons of means
##     95% family-wise confidence level
##
## Fit: aov(formula = Value ~ Treatment, data = OmpA)
##
## $Treatment
##                        diff         lwr       upr     p adj
```

```
## Ind+Bic-Ind    -0.009250502 -0.16824376 0.1497428 0.9826353
## Trp - -Ind      0.198529035  0.03953578 0.3575223 0.0202366
## Trp - -Ind+Bic  0.207779537  0.04878628 0.3667728 0.0165567
```

```
ytgcaov<-aov(Value~Treatment, data=YtgCR)
summary(ytgcaov)
```

```
##             Df  Sum Sq Mean Sq F value   Pr(>F)    
## Treatment    2 0.16647 0.08324   38.67 0.000373 ***
## Residuals    6 0.01292 0.00215                     
## ---
## Signif. codes:  0 '***' 0.001 '**' 0.01 '*' 0.05 '.' 0.1 ' ' 1
```

```
TukeyHSD(x=ytgcaov)
```

```
##   Tukey multiple comparisons of means
##     95% family-wise confidence level
## 
## Fit: aov(formula = Value ~ Treatment, data = YtgCR)
## 
## $Treatment
##                       diff        lwr       upr     p adj
## Ind+Bic-Ind    0.007685001 -0.1085461 0.1239162 0.9776511
## Trp - -Ind     0.292272885  0.1760417 0.4085040 0.0006086
## Trp - -Ind+Bic 0.284587884  0.1683567 0.4008190 0.0007042
```

```
ytgdaov<-aov(Value~Treatment, data=YtgD)
summary(ytgdaov)
```

```
##             Df  Sum Sq  Mean Sq F value   Pr(>F)    
## Treatment    2 0.03191 0.015956   60.43 0.000106 ***
## Residuals    6 0.00158 0.000264                     
## ---
## Signif. codes:  0 '***' 0.001 '**' 0.01 '*' 0.05 '.' 0.1 ' ' 1
```

```
TukeyHSD(x=ytgdaov)
```

```
##   Tukey multiple comparisons of means
##     95% family-wise confidence level
## 
## Fit: aov(formula = Value ~ Treatment, data = YtgD)
## 
## $Treatment
##                      diff         lwr        upr     p adj
## Ind+Bic-Ind    0.04431689 0.003608391 0.08502539 0.0359205
## Trp - -Ind     0.14250572 0.101797224 0.18321422 0.0000949
## Trp - -Ind+Bic 0.09818883 0.057480334 0.13889733 0.0007642
```

```
#Plot each gene individually w/ annotated significance for TukeyHSD
#CTL0174
ggbarplot(CTL0174, x="Treatment2", y="Value", add="mean_sd", color="black",
fill="Treatment2", palette=c("darkorchid4", "darkorchid3", "darkorchid2"),
```

```
title="CTL0174") + scale_y_continuous(limits=c(0, 2.2),
expand=expand_scale(mult=c(0,0.1))) + ylab(label="3'-to-5' Ratio") +
rremove("legend") + theme(axis.text.x=element_text(angle=45, hjust=1)) +
rremove("xlab") + geom_exec(geomfunc=geom_point, data=CTL0174,
x="Treatment2", y="Value", colour="black") +
geom_signif(comparisons=list(c("Trp - ", "Ind"), c("Trp - ", "Ind+Bic"),
c("Ind", "Ind+Bic")), annotation=c("ns", "ns", "ns"), y_position = c(1.7,
1.9, 2.1)) + theme(line=element_line(size=1, colour="black"),
panel.border=element_rect(colour="black", fill="NA", size=1)) +
theme(plot.title=element_text(hjust='0.5'))
```

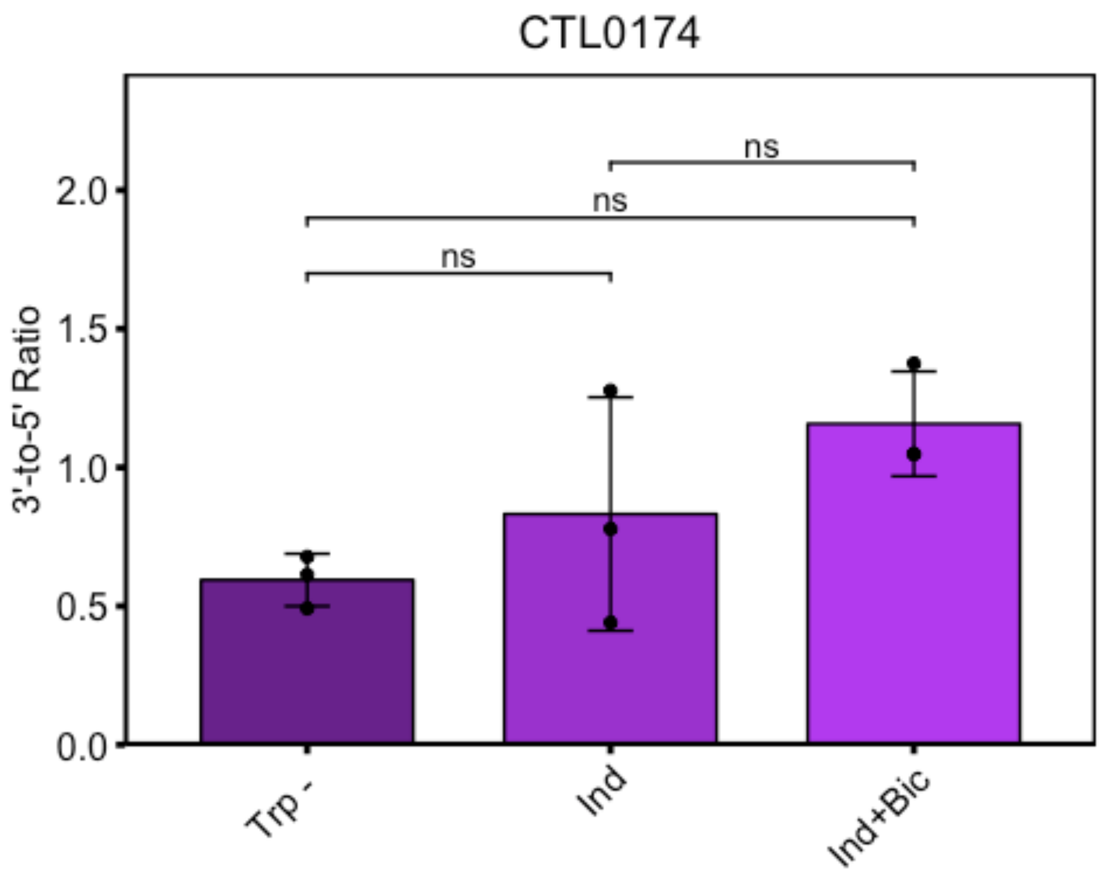


```
#YtgCR
ggbarplot(YtgCR, x="Treatment2", y="Value", add="mean_sd", color="black",
fill="Treatment2", palette=c("darkorchid4", "darkorchid3", "darkorchid2"),
title="YtgCR") + scale_y_continuous(limits=c(0, 2.2),
expand=expand_scale(mult=c(0,0.1))) + ylab(label="3'-to-5' Ratio") +
rremove("legend") + theme(axis.text.x=element_text(angle=45, hjust=1)) +
rremove("xlab") + geom_exec(geomfunc=geom_point, data=YtgCR, x="Treatment2",
y="Value", colour="black") + geom_signif(comparisons=list(c("Trp - ", "Ind"),
c("Trp - ", "Ind+Bic"), c("Ind", "Ind+Bic")), annotation=c("***", "***",
"ns"), y_position = c(1.7, 1.9, 2.1)) + theme(line=element_line(size=1,
colour="black"), panel.border=element_rect(colour="black", fill="NA",
size=1)) + theme(plot.title=element_text(hjust='0.5'))
```

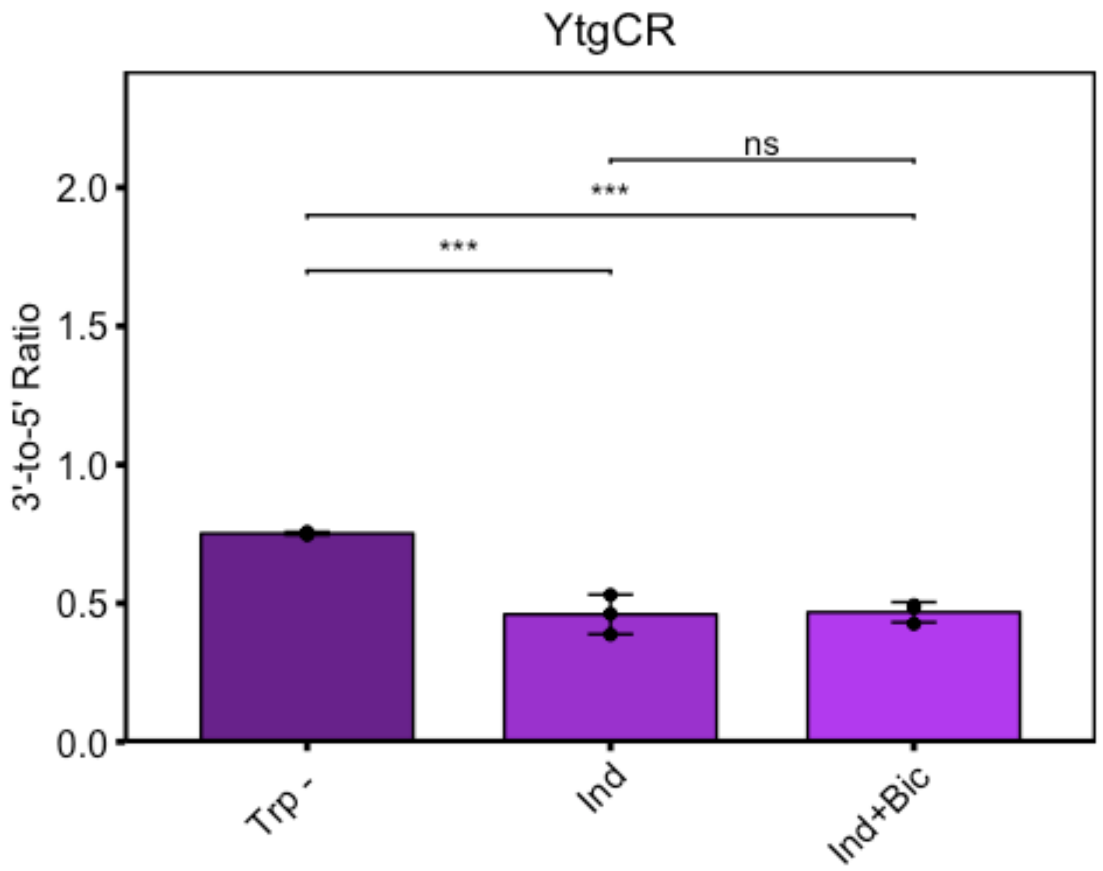


```
#GroEL_1
ggbarplot(GroEL_1, x="Treatment2", y="Value", add="mean_sd", color="black",
fill="Treatment2", palette=c("darkorchid4", "darkorchid3", "darkorchid2"),
title="GroEL_1") + scale_y_continuous(limits=c(0, 2.2),
expand=expand_scale(mult=c(0,0.1))) + ylab(label="3'-to-5' Ratio") +
rremove("legend") + theme(axis.text.x=element_text(angle=45, hjust=1)) +
rremove("xlab") + geom_exec(geomfunc=geom_point, data=GroEL_1,
x="Treatment2", y="Value", colour="black") +
geom_signif(comparisons=list(c("Trp - ", "Ind"), c("Trp - ", "Ind+Bic"),
c("Ind", "Ind+Bic")), annotation=c("ns", "ns", "ns"), y_position = c(1.7,
1.9, 2.1)) + theme(line=element_line(size=1, colour="black"),
panel.border=element_rect(colour="black", fill="NA", size=1)) +
theme(plot.title=element_text(hjust='0.5'))
```

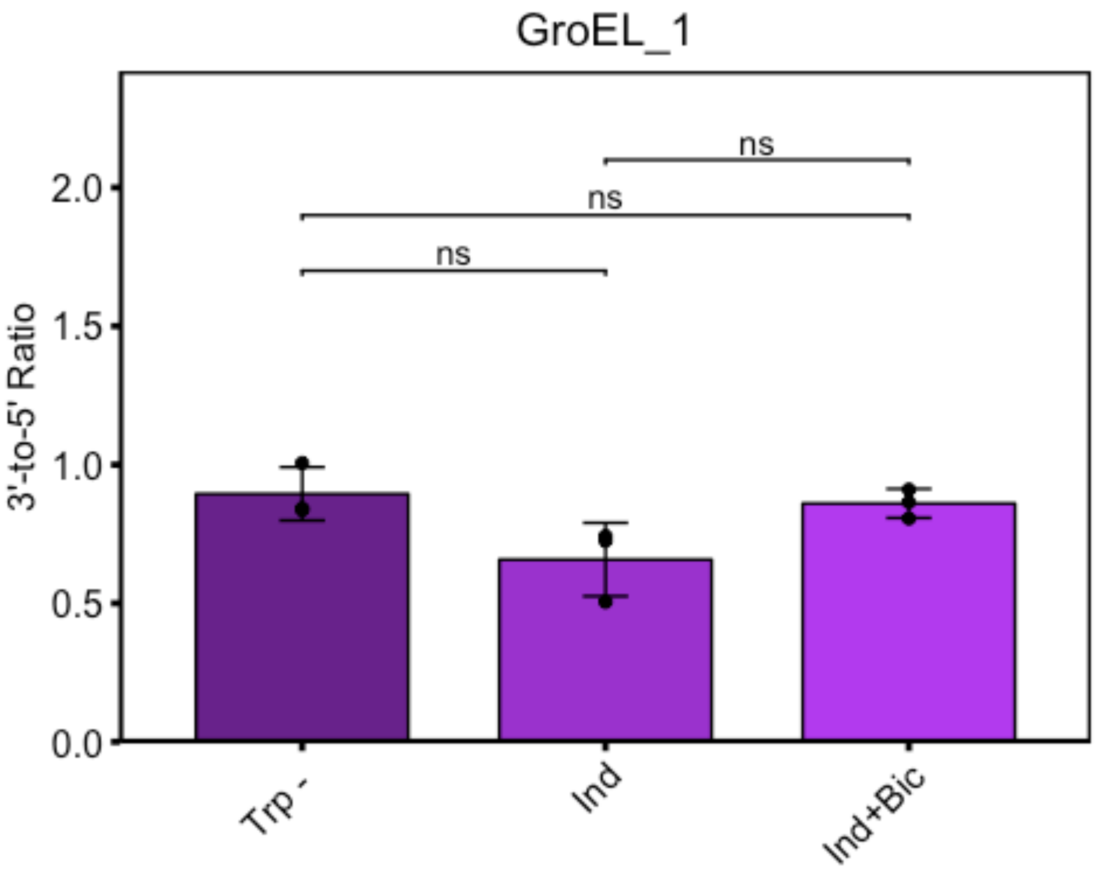


```
#OmpA
ggbarplot(OmpA, x="Treatment2", y="Value", add="mean_sd", color="black",
fill="Treatment2", palette=c("darkorchid4", "darkorchid3", "darkorchid2"),
title="OmpA") + scale_y_continuous(limits=c(0, 2.2),
expand=expand_scale(mult=c(0,0.1))) + ylab(label="3'-to-5' Ratio") +
rremove("legend") + theme(axis.text.x=element_text(angle=45, hjust=1)) +
rremove("xlab") + geom_exec(geomfunc=geom_point, data=OmpA, x="Treatment2",
y="Value", colour="black") + geom_signif(comparisons=list(c("Trp - ", "Ind"),
c("Trp - ", "Ind+Bic"), c("Ind", "Ind+Bic")), annotation=c("*", "*", "ns"),
y_position = c(1.7, 1.9, 2.1)) + theme(line=element_line(size=1,
colour="black"), panel.border=element_rect(colour="black", fill="NA",
size=1)) + theme(plot.title=element_text(hjust='0.5'))
```

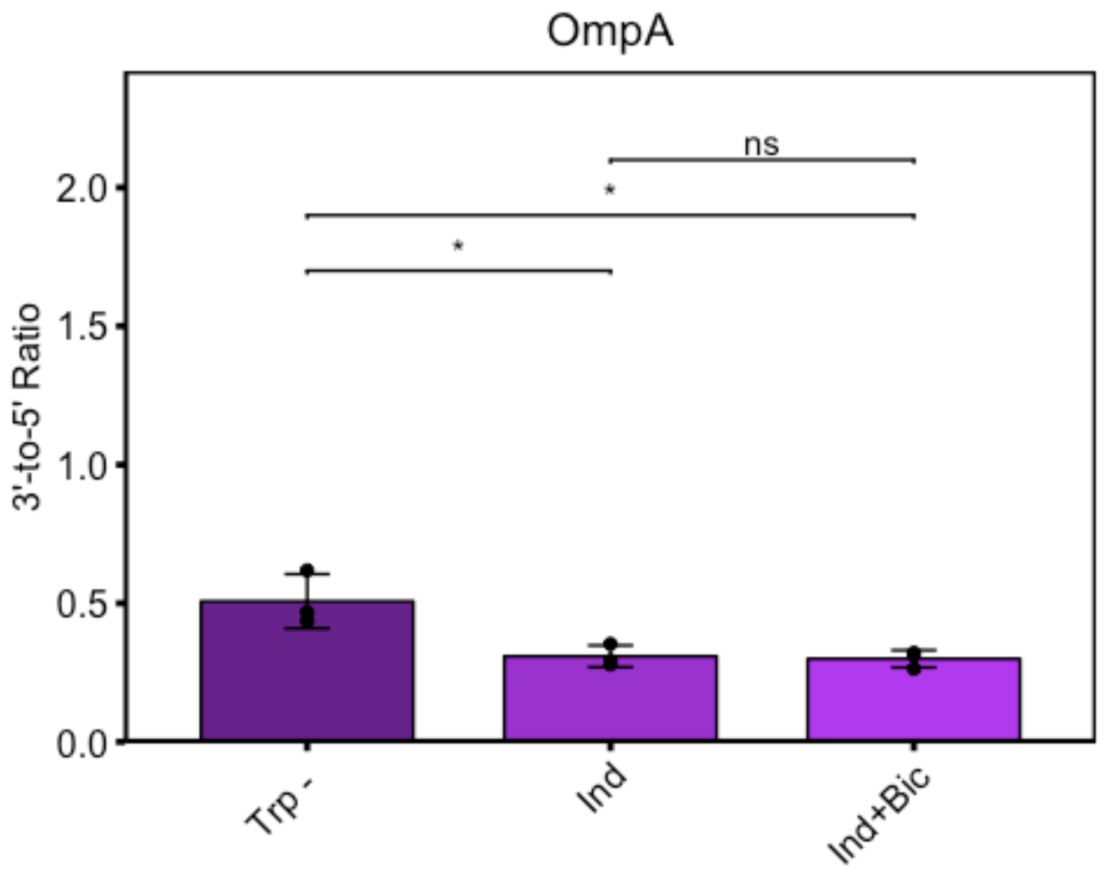


```
#YtgD:YtgA
ggbarplot(YtgD, x="Treatment2", y="Value", add="mean_sd", color="black",
fill="Treatment2", palette=c("darkorchid4", "darkorchid3", "darkorchid2"),
title="YtgD:YtgA") + scale_y_continuous(limits=c(0, 2.2),
expand=expand_scale(mult=c(0,0.1))) + ylab(label="3'-to-5' Ratio") +
rremove("legend") + theme(axis.text.x=element_text(angle=45, hjust=1)) +
rremove("xlab") + geom_exec(geomfunc=geom_point, data=YtgD, x="Treatment2",
y="Value", colour="black") + geom_signif(comparisons=list(c("Trp - ", "Ind"),
c("Trp - ", "Ind+Bic"), c("Ind", "Ind+Bic")), annotation=c("***", "***",
"*"), y_position = c(1.7, 1.9, 2.1)) + theme(line=element_line(size=1,
colour="black"), panel.border=element_rect(colour="black", fill="NA",
size=1)) + theme(plot.title=element_text(hjust='0.5'))
```

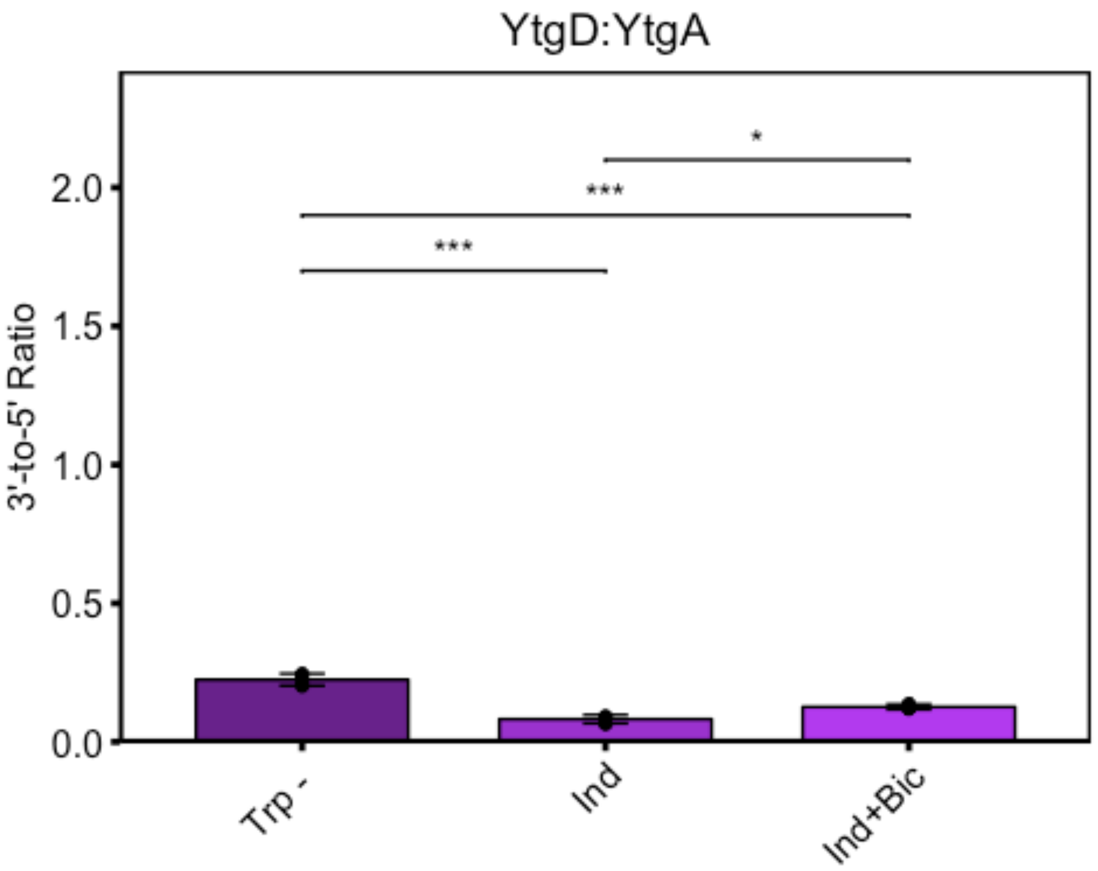

YtgD:YtgA
3'-to-5' Ratio
2.0
1.5
1.0
0.5
0.0
*
***
***
Trp -
Ind
Ind+Bic

### Figure 5

Nick Pokorzynski

4/16/2020

```
#Fig5b
setwd("~/Documents/Carabeo Lab/R Code/Paper 2/Figure 5")
library(ggplot2)
library(ggpubr)
```

```
## Warning: package 'ggpubr' was built under R version 3.5.2
```

```
## Loading required package: magrittr
```

```
bpd<-read.csv("Fig5c.csv")
p<-ggdensity(bpd, x="Position", fill="Promoter", y="..scaled..",
position="stack", palette=c("dodgerblue3"), title="24 Bpd", color="Promoter")
+ scale_y_continuous(name="Count Density",
expand=expand_scale(mult=c(0,0.1))) +
theme(panel.border=element_rect(colour="black", fill="NA", size=1),
axis.text=element_text(color='black')) + xlab(label="C. trachomatis L2 434/Bu
Genome Position") + scale_x_continuous(limits=c(511300, 512100)) +
theme(plot.title=element_text(hjust='0.5'))
ggpar(p, legend=c("right"))
```

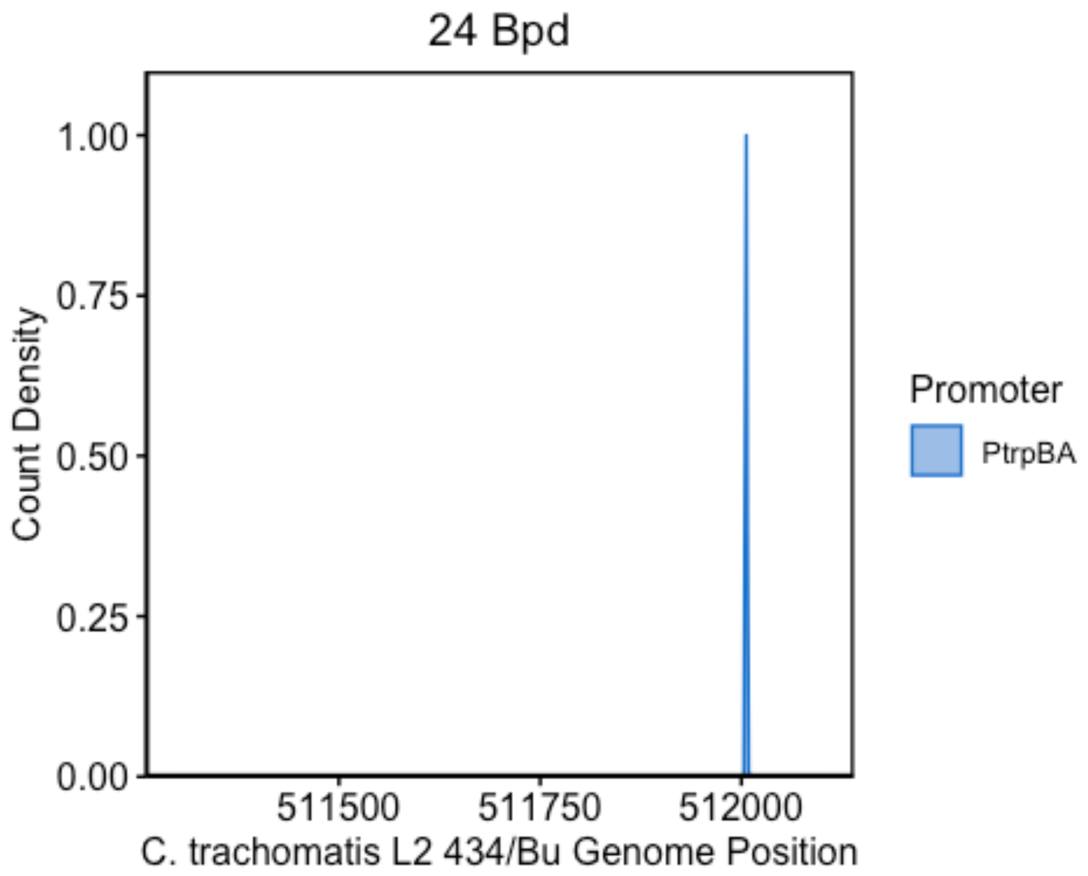


```
#Fig5c
setwd("~/Documents/Carabeo Lab/R Code/Paper 2/Figure 5")
library(ggplot2)
library(ggpubr)
trp<-read.csv("Fig5d.csv")
p<-ggdensity(trp, x="Position", y="..scaled..", fill="Promoter",
position="stack", palette=c("violetred4", "violetred1"), title="24 Trp-",
color="Promoter") + scale_y_continuous(name="Count Density",
expand=expand_scale(mult=c(0,0.1))) +
theme(panel.border=element_rect(colour="black", fill="NA", size=1),
axis.text=element_text(color='black')) + xlab(label="C. trachomatis L2 434/Bu
Genome Position") + scale_x_continuous(limits=c(511300, 512100)) +
theme(plot.title=element_text(hjust='0.5'))
ggpar(p, legend=c("right"))
```

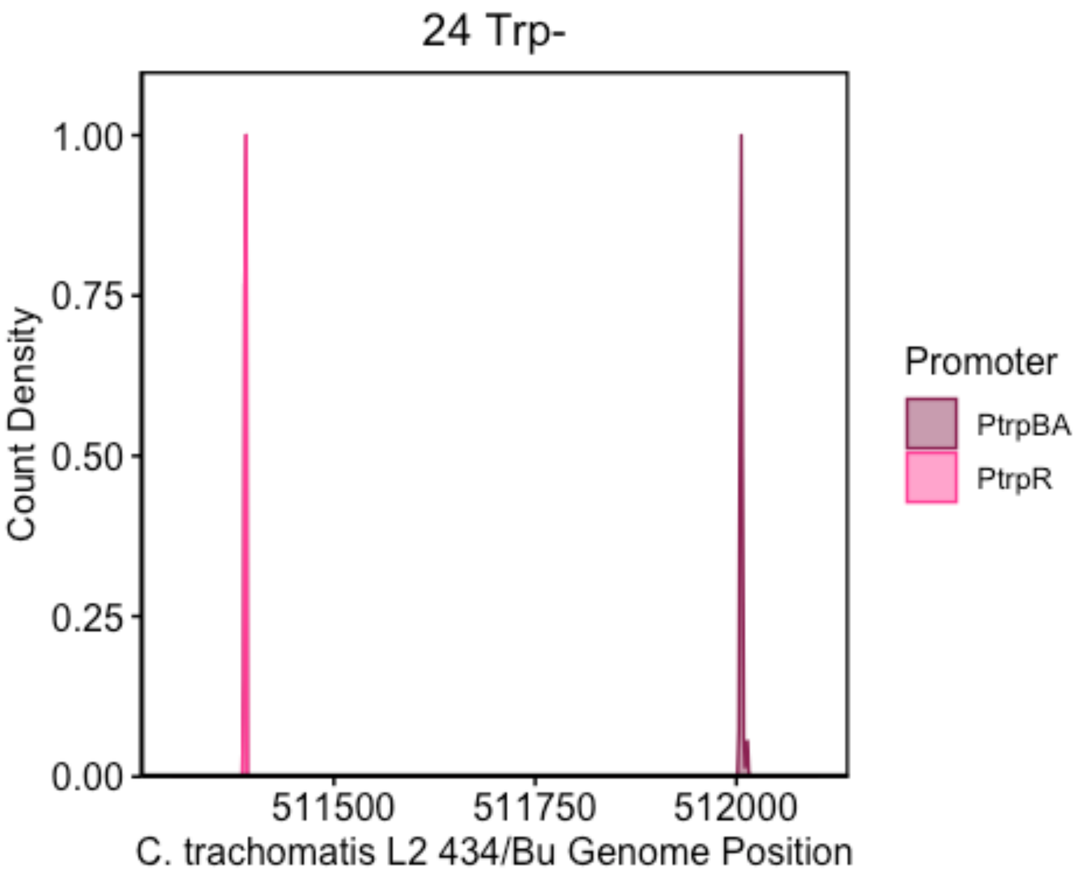


```
#Fig5d
setwd("~/Documents/Carabeo Lab/R Code/Paper 2/Figure 5")
library(ggplot2)
library(ggpubr)
library(ggsignif)

## Warning: package 'ggsignif' was built under R version 3.5.2

p<-read.csv("Fig5a.csv")
paov<-aov(Value~ID, data=p)
summary(paov)

##             Df Sum Sq Mean Sq F value  Pr(>F)
## ID           4   4.80   1.200   6.977 0.00598 **
## Residuals   10   1.72   0.172
## ---
## Signif. codes:  0 '***' 0.001 '**' 0.01 '*' 0.05 '.' 0.1 ' ' 1

TukeyHSD(x=paov)

##   Tukey multiple comparisons of means
##     95% family-wise confidence level
##
## Fit: aov(formula = Value ~ ID, data = p)
```

```
##
## $ID
##                          diff        lwr        upr     p adj
## 18+6Trp--18+6Bpd -0.22341281 -1.3379178  0.8910922 0.9607634
## 24 hpi-18+6Bpd   -1.52440288 -2.6389079 -0.4098979 0.0078456
## 24Bpd-18+6Bpd    -0.03770382 -1.1522088  1.0768012 0.9999586
## 24Trp--18+6Bpd   -0.28049509 -1.3950001  0.8340099 0.9157907
## 24 hpi-18+6Trp-  -1.30099007 -2.4154951 -0.1864851 0.0213024
## 24Bpd-18+6Trp-    0.18570898 -0.9287960  1.3002140 0.9796766
## 24Trp--18+6Trp-  -0.05708228 -1.1715873  1.0574227 0.9997845
## 24Bpd-24 hpi      1.48669905  0.3721940  2.6012041 0.0092608
## 24Trp--24 hpi     1.24390779  0.1294028  2.3584128 0.0276339
## 24Trp--24Bpd     -0.24279127 -1.3572963  0.8717137 0.9477743

p$ID2<-factor(p$ID, levels=c("24 hpi", "18+6Bpd", "18+6Trp-", "24Bpd",
"24Trp-"))
p$Treatment2<-factor(p$Treatment, levels=c("Mock", "BPD", "Trp"))
ggbarplot(p, x="ID2", y="Value", add="mean_sd", fill="Treatment2",
palette=c("lightsteelblue4", "lightsteelblue1", "lightsteelblue3")) +
scale_y_continuous(expand=expand_scale(mult=c(0,0.1))) + ylab(label="PtrpBA
Activity (trpB:trpR)") + rremove("legend") +
theme(axis.text.x=element_text(angle=45, hjust=1)) + rremove("xlab") +
geom_exec(geomfunc=geom_point, data=p, x="ID2", y="Value", colour="black") +
geom_signif(comparisons=list(c("24 hpi", "18+6Bpd"), c("24 hpi", "18+6Trp-"),
c("24 hpi", "24Bpd"), c("24 hpi", "24Trp-")), annotation=c("**", "*", "**",
"*"), y_position=c(3.5, 4, 4.5, 5)) + theme(line=element_line(size=1,
colour="black"), panel.border=element_rect(colour="black", fill="NA",
size=1)) + theme(plot.title=element_text(hjust='0.5')) +
geom_hline(yintercept=1, linetype=2)
```

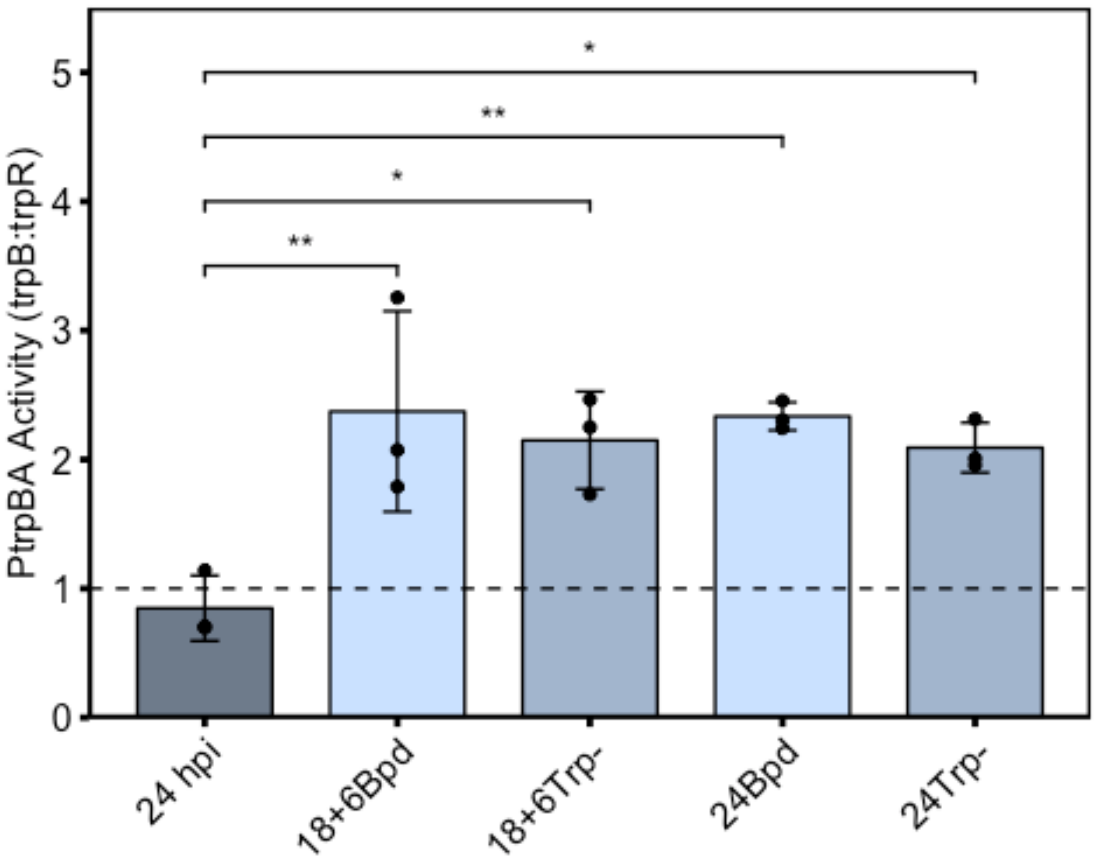

PtrpBA Activity (trpB:trpR)
0
1
2
3
4
5
**
*
**
*
24 hpi
18+6Bpd
18+6Trp-
24Bpd
24Trp-

### Supplementary Figure 1

Nick Pokorzynski

4/14/2020

```
#Supplementary Figure 1
setwd("~/Documents/Carabeo Lab/R Code/Paper 2/Supplementary Figure 1")
library(ggplot2)
library(ggpubr)

## Warning: package 'ggpubr' was built under R version 3.5.2

## Loading required package: magrittr

rt<-read.csv("SuppFig1.csv")
rt$Treatment2<-factor(rt$Treatment, levels=c("Mock", "Bpd", "Trp"))
rt$ID2<-factor(rt$ID, levels=c("24 hpi", "18+6 Bpd", "18+6 Trp-", "24 Bpd",
"24 Trp-"))
ytgA<-subset(rt, Gene%in%c("ytgA"))
ytgAaov<-aov(Log2_Value~ID, data=ytgA)
summary(ytgAaov)

##             Df Sum Sq Mean Sq F value   Pr(>F)    
## ID           4 10.438  2.6095   17.14 0.000179 ***
## Residuals   10  1.522  0.1522                     
## ---
## Signif. codes:  0 '***' 0.001 '**' 0.01 '*' 0.05 '.' 0.1 ' ' 1

TukeyHSD(x=ytgAaov)

##   Tukey multiple comparisons of means
##     95% family-wise confidence level
## 
## Fit: aov(formula = Log2_Value ~ ID, data = ytgA)
## 
## $ID
##                           diff         lwr        upr     p adj
## 18+6 Trp--18+6 Bpd  0.2946426 -0.75370006  1.3429852 0.8809175
## 24 Bpd-18+6 Bpd     1.7021401  0.65379748  2.7504827 0.0023406
## 24 hpi-18+6 Bpd    -0.5739096 -1.62225219  0.4744330 0.4230307
## 24 Trp--18+6 Bpd    1.2722600  0.22391735  2.3206026 0.0168636
## 24 Bpd-18+6 Trp-    1.4074975  0.35915492  2.4558402 0.0088758
## 24 hpi-18+6 Trp-   -0.8685521 -1.91689475  0.1797905 0.1189154
## 24 Trp--18+6 Trp-   0.9776174 -0.07072521  2.0259600 0.0705137
## 24 hpi-24 Bpd      -2.2760497 -3.32439229 -1.2277071 0.0002343
## 24 Trp--24 Bpd     -0.4298801 -1.47822275  0.6184625 0.6696059
## 24 Trp--24 hpi      1.8461695  0.79782692  2.8945122 0.0012662
```

```
ggbarplot(ytgA, x="ID2", y="Log2_Value", add="mean_sd", fill="Treatment2",
palette=c("lightsteelblue4", "lightsteelblue1", "lightsteelblue3"),
title="ytgA") + scale_y_continuous(expand=expand_scale(mult=c(0,0.1))) +
ylab(label="Log2 Transcript Expression") + rremove("legend") +
theme(axis.text.x=element_text(angle=45, hjust=1)) + rremove("xlab") +
geom_exec(geomfunc=geom_point, data=ytgA, x="ID2", y="Log2_Value",
colour="black") + geom_signif(comparisons=list(c("24 hpi", "18+6 Bpd"), c("24
hpi", "18+6 Trp-"), c("24 hpi", "24 Bpd"), c("24 hpi", "24 Trp-"), c("18+6
Bpd", "18+6 Trp-"), c("24 Bpd", "24 Trp-")), annotation=c("ns", "ns", "***",
"**", "ns", "ns"), y_position=c(5, 11, 12, 13, 9.5, 10.5)) +
theme(line=element_line(size=1, colour="black"),
panel.border=element_rect(colour="black", fill="NA", size=1)) +
theme(plot.title=element_text(hjust='0.5', face="italic"))
```

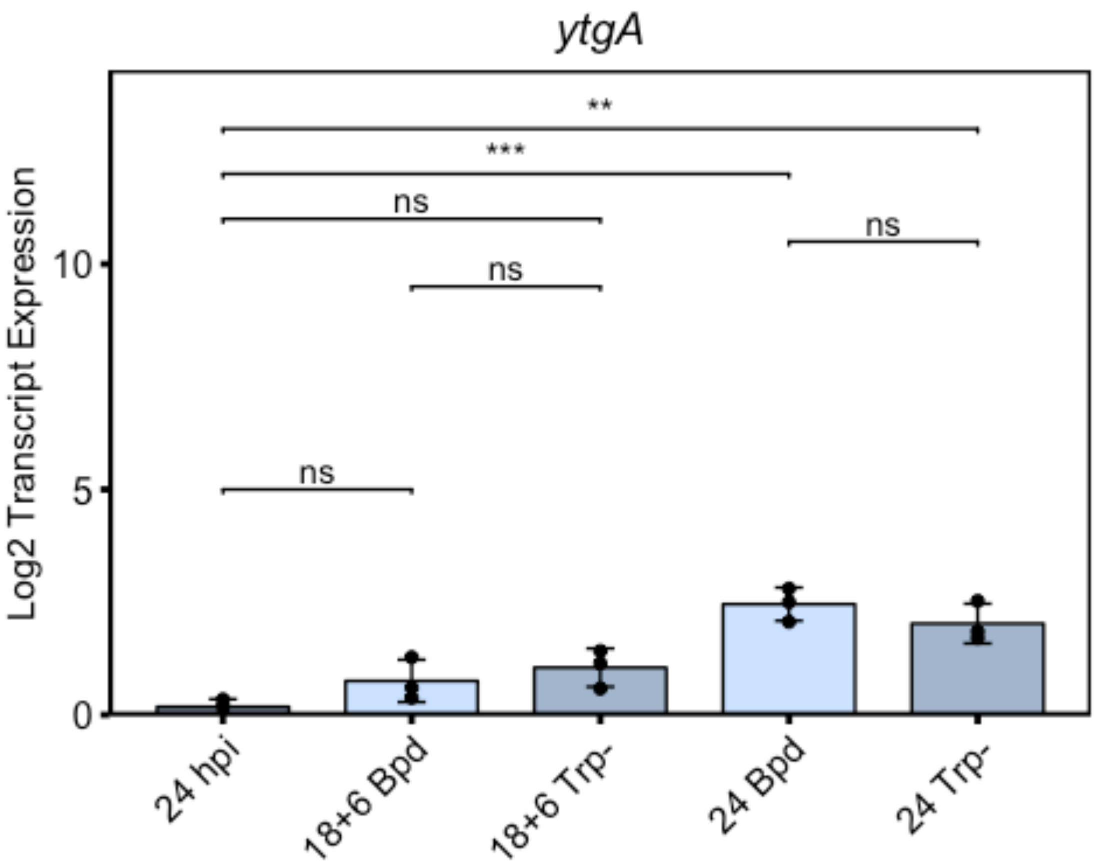


```
nrdA<-subset(rt, Gene%in%c("nrdA"))
nrdAaov<-aov(Log2_Value~ID, data=nrdA)
summary(nrdAaov)

##              Df Sum Sq Mean Sq F value  Pr(>F)
## ID            4 14.120   3.530   14.07 0.00041 ***
## Residuals    10  2.509   0.251
## ---
## Signif. codes:  0 '***' 0.001 '**' 0.01 '*' 0.05 '.' 0.1 ' ' 1
```

```
TukeyHSD(x=nrdAaov)

##   Tukey multiple comparisons of means
##     95% family-wise confidence level
##
## Fit: aov(formula = Log2_Value ~ ID, data = nrdA)
##
## $ID
##                           diff        lwr        upr     p adj
## 18+6 Trp--18+6 Bpd -0.22968355 -1.5756635  1.1162964 0.9778460
## 24 Bpd-18+6 Bpd    -0.19865523 -1.5446352  1.1473247 0.9869792
## 24 hpi-18+6 Bpd    -2.40435462 -3.7503345 -1.0583747 0.0011339
## 24 Trp--18+6 Bpd    0.31625533 -1.0297246  1.6622353 0.9327062
## 24 Bpd-18+6 Trp-    0.03102832 -1.3149516  1.3770083 0.9999910
## 24 hpi-18+6 Trp-   -2.17467106 -3.5206510 -0.8286911 0.0024274
## 24 Trp--18+6 Trp-   0.54593888 -0.8000411  1.8919188 0.6778278
## 24 hpi-24 Bpd      -2.20569939 -3.5516793 -0.8597195 0.0021852
## 24 Trp--24 Bpd      0.51491055 -0.8310694  1.8608905 0.7198936
## 24 Trp--24 hpi      2.72060994  1.3746300  4.0665899 0.0004234

ggbarplot(nrdA, x="ID2", y="Log2_Value", add="mean_sd", fill="Treatment2",
palette=c("lightsteelblue4", "lightsteelblue1", "lightsteelblue3"),
title="nrdA") + scale_y_continuous(expand=expand_scale(mult=c(0,0.1))) +
ylab(label="Log2 Transcript Expression") + rremove("legend") +
theme(axis.text.x=element_text(angle=45, hjust=1)) + rremove("xlab") +
geom_exec(geomfunc=geom_point, data=nrdA, x="ID2", y="Log2_Value",
colour="black") + geom_signif(comparisons=list(c("24 hpi", "18+6 Bpd"), c("24
hpi", "18+6 Trp-"), c("24 hpi", "24 Bpd"), c("24 hpi", "24 Trp-"), c("18+6
Bpd", "18+6 Trp-"), c("24 Bpd", "24 Trp-")), annotation=c("**", "**", "**",
"***", "ns", "ns"), y_position=c(5, 11, 12, 13, 9.5, 10.5)) +
theme(line=element_line(size=1, colour="black"),
panel.border=element_rect(colour="black", fill="NA", size=1)) +
theme(plot.title=element_text(hjust='0.5', face="italic"))
```

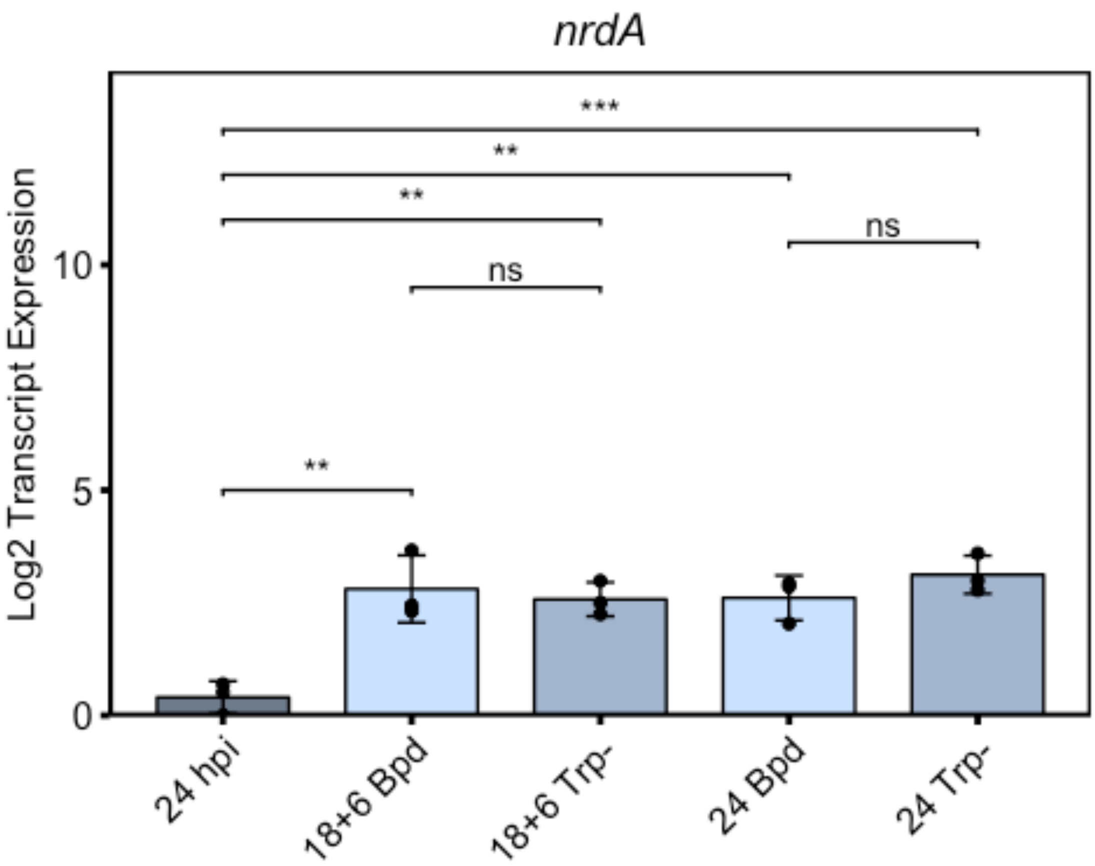


```
nrdB<-subset(rt, Gene%in%c("nrdB"))
nrdBaov<-aov(Log2_Value~ID, data=nrdB)
summary(nrdBaov)

##             Df Sum Sq Mean Sq F value   Pr(>F)
## ID           4 11.410  2.8526   11.81 0.000833 ***
## Residuals   10  2.415  0.2415
## ---
## Signif. codes:  0 '***' 0.001 '**' 0.01 '*' 0.05 '.' 0.1 ' ' 1

TukeyHSD(x=nrdBaov)

##   Tukey multiple comparisons of means
##     95% family-wise confidence level
##
## Fit: aov(formula = Log2_Value ~ ID, data = nrdB)
##
## $ID
##                          diff        lwr        upr     p adj
## 18+6 Trp--18+6 Bpd -0.1398678 -1.4602973  1.1805617 0.9962874
## 24 Bpd-18+6 Bpd     0.5476006 -0.7728289  1.8680302 0.6609772
## 24 hpi-18+6 Bpd    -1.7149568 -3.0353863 -0.3945273 0.0110174
## 24 Trp--18+6 Bpd    0.7744049 -0.5460246  2.0948344 0.3621862
## 24 Bpd-18+6 Trp-    0.6874684 -0.6329611  2.0078979 0.4679752
```

```
## 24 hpi-18+6 Trp-    -1.5750890 -2.8955185 -0.2546595 0.0187214
## 24 Trp--18+6 Trp-    0.9142727 -0.4061568  2.2347022 0.2282578
## 24 hpi-24 Bpd       -2.2625574 -3.5829870 -0.9421279 0.0015617
## 24 Trp--24 Bpd       0.2268043 -1.0936252  1.5472338 0.9773157
## 24 Trp--24 hpi       2.4893617  1.1689322  3.8097912 0.0007423
```

```
ggbarplot(nrdB, x="ID2", y="Log2_Value", add="mean_sd", fill="Treatment2",
palette=c("lightsteelblue4", "lightsteelblue1", "lightsteelblue3"),
title="nrdB") + scale_y_continuous(expand=expand_scale(mult=c(0,0.1))) +
ylab(label="Log2 Transcript Expression") + rremove("legend") +
theme(axis.text.x=element_text(angle=45, hjust=1)) + rremove("xlab") +
geom_exec(geomfunc=geom_point, data=nrdB, x="ID2", y="Log2_Value",
colour="black") + geom_signif(comparisons=list(c("24 hpi", "18+6 Bpd"), c("24
hpi", "18+6 Trp-"), c("24 hpi", "24 Bpd"), c("24 hpi", "24 Trp-"), c("18+6
Bpd", "18+6 Trp-"), c("24 Bpd", "24 Trp-")), annotation=c("*", "*", "**",
"***", "ns", "ns"), y_position=c(5, 11, 12, 13, 9.5, 10.5)) +
theme(line=element_line(size=1, colour="black"),
panel.border=element_rect(colour="black", fill="NA", size=1)) +
theme(plot.title=element_text(hjust='0.5', face="italic"))
```

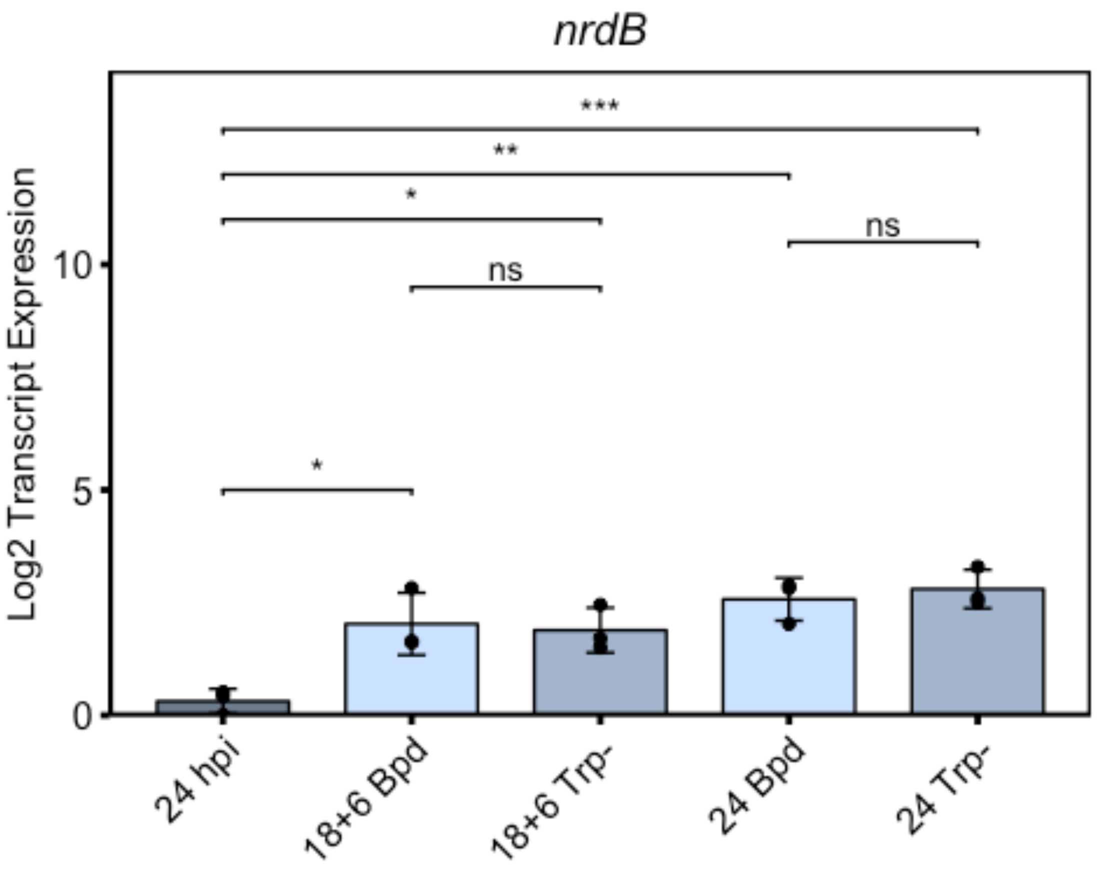


```
devB<-subset(rt, Gene%in%c("devB"))
devBaov<-aov(Log2_Value~ID, data=devB)
summary(devBaov)
```

```
##              Df Sum Sq Mean Sq F value   Pr(>F)    
## ID            4 18.238   4.559   13.38 0.000505 ***
## Residuals    10  3.409   0.341                     
## ---
## Signif. codes:  0 '***' 0.001 '**' 0.01 '*' 0.05 '.' 0.1 ' ' 1
```

```
TukeyHSD(x=devBaov)
```

```
##   Tukey multiple comparisons of means
##     95% family-wise confidence level
## 
## Fit: aov(formula = Log2_Value ~ ID, data = devB)
## 
## $ID
##                             diff          lwr        upr     p adj
## 18+6 Trp--18+6 Bpd  0.3283291 -1.240613727  1.8972719 0.9544718
## 24 Bpd-18+6 Bpd     2.4380769  0.869134054  4.0070197 0.0032271
## 24 hpi-18+6 Bpd    -0.3590856 -1.928028422  1.2098572 0.9383308
## 24 Trp--18+6 Bpd    1.8994723  0.330529493  3.4684151 0.0171135
## 24 Bpd-18+6 Trp-    2.1097478  0.540804962  3.6786906 0.0087847
## 24 hpi-18+6 Trp-   -0.6874147 -2.256357514  0.8815281 0.6173763
## 24 Trp--18+6 Trp-   1.5711432  0.002200401  3.1400860 0.0496430
## 24 hpi-24 Bpd      -2.7971625 -4.366105295 -1.2282197 0.0011513
## 24 Trp--24 Bpd     -0.5386046 -2.107547380  1.0303383 0.7881113
## 24 Trp--24 hpi      2.2585579  0.689615096  3.8275007 0.0055430
```

```
ggbarplot(devB, x="ID2", y="Log2_Value", add="mean_sd", fill="Treatment2",
palette=c("lightsteelblue4", "lightsteelblue1", "lightsteelblue3"),
title="devB") + scale_y_continuous(expand=expand_scale(mult=c(0,0.1))) +
ylab(label="Log2 Transcript Expression") + rremove("legend") +
theme(axis.text.x=element_text(angle=45, hjust=1)) + rremove("xlab") +
geom_exec(geomfunc=geom_point, data=devB, x="ID2", y="Log2_Value",
colour="black") + geom_signif(comparisons=list(c("24 hpi", "18+6 Bpd"), c("24
hpi", "18+6 Trp-"), c("24 hpi", "24 Bpd"), c("24 hpi", "24 Trp-"), c("18+6
Bpd", "18+6 Trp-"), c("24 Bpd", "24 Trp-")), annotation=c("ns", "ns", "**",
"**", "ns", "ns"), y_position=c(5, 11, 12, 13, 9.5, 10.5)) +
theme(line=element_line(size=1, colour="black"),
panel.border=element_rect(colour="black", fill="NA", size=1)) +
theme(plot.title=element_text(hjust='0.5', face="italic"))
```

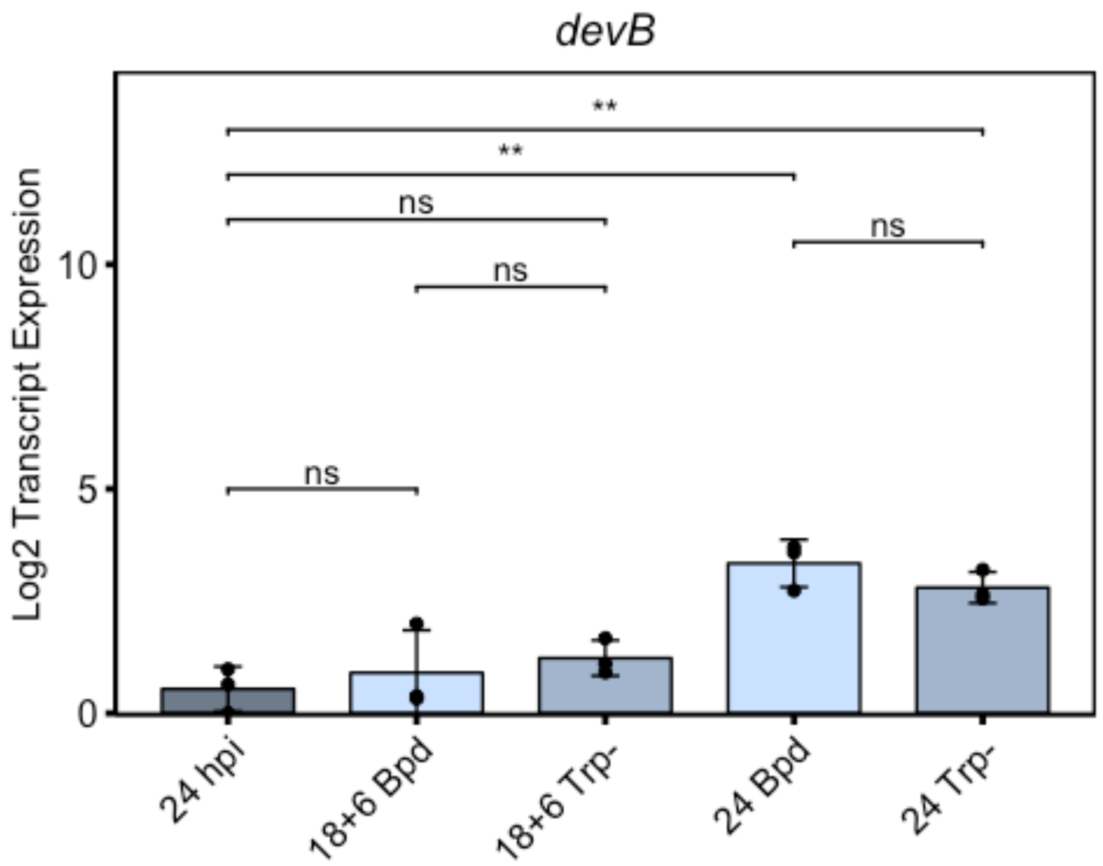


```
ahpC<-subset(rt, Gene%in%c("ahpC"))
ahpCaov<-aov(Log2_Value~ID, data=ahpC)
summary(ahpCaov)
```

```
##             Df Sum Sq Mean Sq F value   Pr(>F)    
## ID           4 23.427   5.857   27.47 2.26e-05 ***
## Residuals   10  2.132   0.213                     
## ---
## Signif. codes:  0 '***' 0.001 '**' 0.01 '*' 0.05 '.' 0.1 ' ' 1
```

```
TukeyHSD(x=ahpCaov)
```

```
##   Tukey multiple comparisons of means
##     95% family-wise confidence level
## 
## Fit: aov(formula = Log2_Value ~ ID, data = ahpC)
## 
## $ID
##                          diff         lwr       upr     p adj
## 18+6 Trp--18+6 Bpd  1.2734998  0.03270228 2.5142973 0.0436993
## 24 Bpd-18+6 Bpd     2.4408362  1.20003863 3.6816337 0.0005279
## 24 hpi-18+6 Bpd    -0.4424367 -1.68323427 0.7983608 0.7656610
## 24 Trp--18+6 Bpd    2.6585278  1.41773023 3.8993253 0.0002617
## 24 Bpd-18+6 Trp-    1.1673364 -0.07346118 2.4081339 0.0676162
```

```
## 24 hpi-18+6 Trp-   -1.7159366 -2.95673409 -0.4751390 0.0072873
## 24 Trp--18+6 Trp-   1.3850280  0.14423042  2.6258255 0.0276153
## 24 hpi-24 Bpd      -2.8832729 -4.12407044 -1.6424754 0.0001318
## 24 Trp--24 Bpd      0.2176916 -1.02310594  1.4584891 0.9755166
## 24 Trp--24 hpi      3.1009645  1.86016697  4.3417620 0.0000703
```

```
ggbarplot(ahpC, x="ID2", y="Log2_Value", add="mean_sd", fill="Treatment2",
palette=c("lightsteelblue4", "lightsteelblue1", "lightsteelblue3"),
title="ahpC") + scale_y_continuous(expand=expand_scale(mult=c(0,0.1))) +
ylab(label="Log2 Transcript Expression") + rremove("legend") +
theme(axis.text.x=element_text(angle=45, hjust=1)) + rremove("xlab") +
geom_exec(geomfunc=geom_point, data=ahpC, x="ID2", y="Log2_Value",
colour="black") + geom_signif(comparisons=list(c("24 hpi", "18+6 Bpd"), c("24
hpi", "18+6 Trp-"), c("24 hpi", "24 Bpd"), c("24 hpi", "24 Trp-"), c("18+6
Bpd", "18+6 Trp-"), c("24 Bpd", "24 Trp-")), annotation=c("ns", "**", "***",
"***", "*", "ns"), y_position=c(5, 11, 12, 13, 9.5, 10.5)) +
theme(line=element_line(size=1, colour="black"),
panel.border=element_rect(colour="black", fill="NA", size=1)) +
theme(plot.title=element_text(hjust='0.5', face="italic"))
```

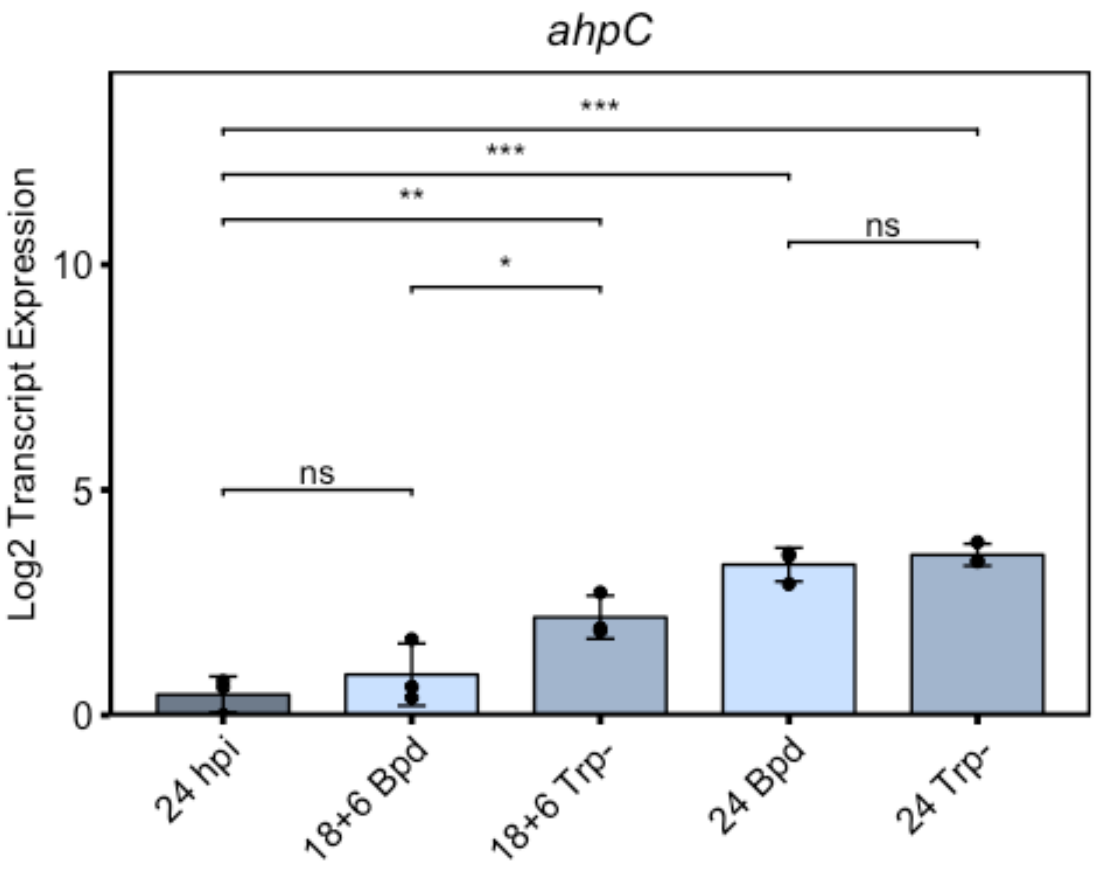

### Supplementary Figure 2

Nick Pokorzynski

4/16/2020

```
#Supplementary Fig 2b
setwd("~/Documents/Carabeo Lab/R Code/Paper 2/Supplementary Figure 2")
library(ggplot2)
library(ggpubr)

## Warning: package 'ggpubr' was built under R version 3.5.2

## Loading required package: magrittr

gdna<-read.csv("SuppFig2b.csv")
gdna$Treatment2<-factor(gdna$Treatment, levels=c("Mock", "Bpd", "Trp", "Bpd +
Trp"))
gdna$ID2<-factor(gdna$ID, levels=c("24 hpi", "18+6 Bpd", "18+6 Trp-", "18+6
Bpd + Trp-"))
gdnaaov<-aov(Value~ID, data=gdna)
summary(gdnaaov)

##             Df    Sum Sq   Mean Sq F value Pr(>F)
## ID           3 2.270e+09 756822407   5.096 0.0292 *
## Residuals    8 1.188e+09 148513214
## ---
## Signif. codes:  0 '***' 0.001 '**' 0.01 '*' 0.05 '.' 0.1 ' ' 1

TukeyHSD(x=gdnaaov)

##   Tukey multiple comparisons of means
##     95% family-wise confidence level
##
## Fit: aov(formula = Value ~ ID, data = gdna)
##
## $ID
##                                  diff         lwr      upr     p adj
## 18+6 Bpd + Trp--18+6 Bpd   -2128.0250 -33992.4427 29736.39 0.9962576
## 18+6 Trp--18+6 Bpd          -422.8413 -32287.2590 31441.58 0.9999700
## 24 hpi-18+6 Bpd            30862.6759  -1001.7418 62727.09 0.0576198
## 18+6 Trp--18+6 Bpd + Trp-   1705.1837 -30159.2340 33569.60 0.9980584
## 24 hpi-18+6 Bpd + Trp-     32990.7009   1126.2831 64855.12 0.0426520
## 24 hpi-18+6 Trp-           31285.5172   -578.9006 63149.93 0.0542688

ggbarplot(gdna, x="ID2", y="Log10_Value", add="mean_sd", fill="Treatment2",
palette=c("darkseagreen4", "darkseagreen", "darkseagreen3", "darkseagreen1"))
+ scale_y_continuous(expand=expand_scale(mult=c(0,0.1))) + ylab(label="Log10
Genome Equivalents/ng gDNA") + rremove("legend") +
```

```
theme(axis.text.x=element_text(angle=45, hjust=1)) + rremove("xlab") +
geom_exec(geomfunc=geom_point, data=gdna, x="ID2", y="Log10_Value",
colour="black") + geom_signif(comparisons=list(c("24 hpi", "18+6 Bpd"), c("24
hpi", "18+6 Trp-"), c("24 hpi", "18+6 Bpd + Trp-")), annotation=c("p =
0.0576", "p = 0.0543", "*"), y_position=c(5.5,6.5,7.5)) +
theme(line=element_line(size=1, colour="black"),
panel.border=element_rect(colour="black", fill="NA", size=1))
```

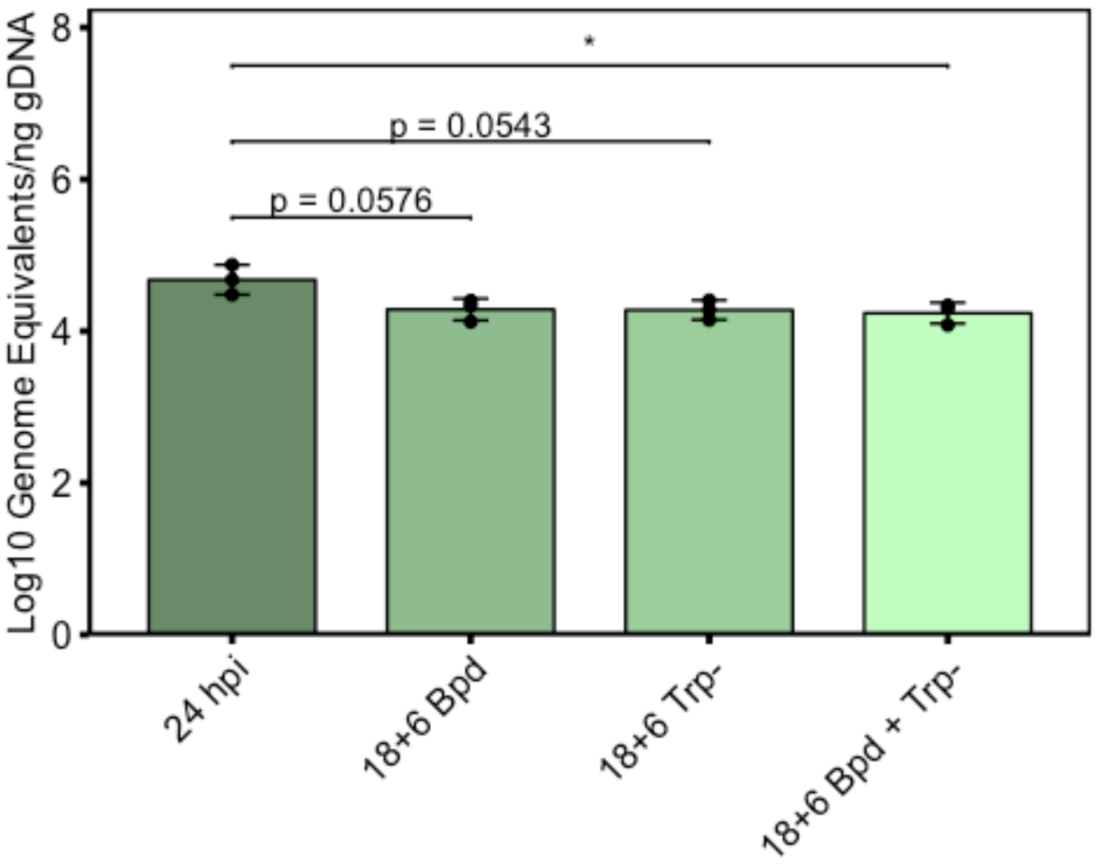


```
#Supplementary Fig 2c
setwd("~/Documents/Carabeo Lab/R Code/Paper 2/Supplementary Figure 2")
library(ggplot2)
library(ggpubr)
rt<-read.csv("SuppFig2c.csv")
rt$ID2<-factor(rt$ID, levels=c("24 hpi", "18+6 Bpd", "18+6 Trp-", "18+6 Bpd +
Trp-"))
rt$Treatment2<-factor(rt$Treatment, levels=c("Mock", "Bpd", "Trp", "Bpd +
Trp"))
#euo
euo<-subset(rt, Gene%in%c("euo"))
euoaov<-aov(Log2_Value~ID, data=euo)
summary(euoaov)
```

```
##              Df Sum Sq Mean Sq F value Pr(>F)
## ID            3  2.906  0.9687   1.956  0.199
## Residuals     8  3.962  0.4953
```

```
TukeyHSD(x=euoaov)
```

```
##   Tukey multiple comparisons of means
##     95% family-wise confidence level
##
## Fit: aov(formula = Log2_Value ~ ID, data = euo)
##
## $ID
##                                 diff       lwr       upr     p adj
## 18+6 Bpd + Trp--18+6 Bpd  -0.7211714 -2.561324 1.1189815 0.6128068
## 18+6 Trp--18+6 Bpd        -0.2126230 -2.052776 1.6275300 0.9814903
## 24 hpi-18+6 Bpd           -1.2732494 -3.113402 0.5669035 0.1985166
## 18+6 Trp--18+6 Bpd + Trp-  0.5085485 -1.331604 2.3487014 0.8127793
## 24 hpi-18+6 Bpd + Trp-    -0.5520780 -2.392231 1.2880749 0.7744874
## 24 hpi-18+6 Trp-          -1.0606265 -2.900779 0.7795265 0.3210705
```

```
ggbarplot(euo, x="ID2", y="Log2_Value", add="mean_sd", fill="Treatment2",
palette=c("darkseagreen4", "darkseagreen", "darkseagreen3", "darkseagreen1"),
title="euo") + scale_y_continuous(expand=expand_scale(mult=c(0,0.1))) +
ylab(label="Log2 Transcript Expression") + rremove("legend") +
theme(axis.text.x=element_text(angle=45, hjust=1)) + rremove("xlab") +
geom_exec(geomfunc=geom_point, data=euo, x="ID2", y="Log2_Value",
colour="black") + geom_signif(comparisons=list(c("18+6 Trp-", "18+6 Bpd +
Trp-"), c("24 hpi", "18+6 Bpd"), c("24 hpi", "18+6 Trp-"), c("24 hpi", "18+6
Bpd + Trp-")), annotation=c("ns", "ns", "ns", "ns"), y_position=c(10, 11, 12,
13)) + theme(line=element_line(size=1, colour="black"),
panel.border=element_rect(colour="black", fill="NA", size=1)) +
theme(plot.title=element_text(hjust='0.5', face="italic"))
```

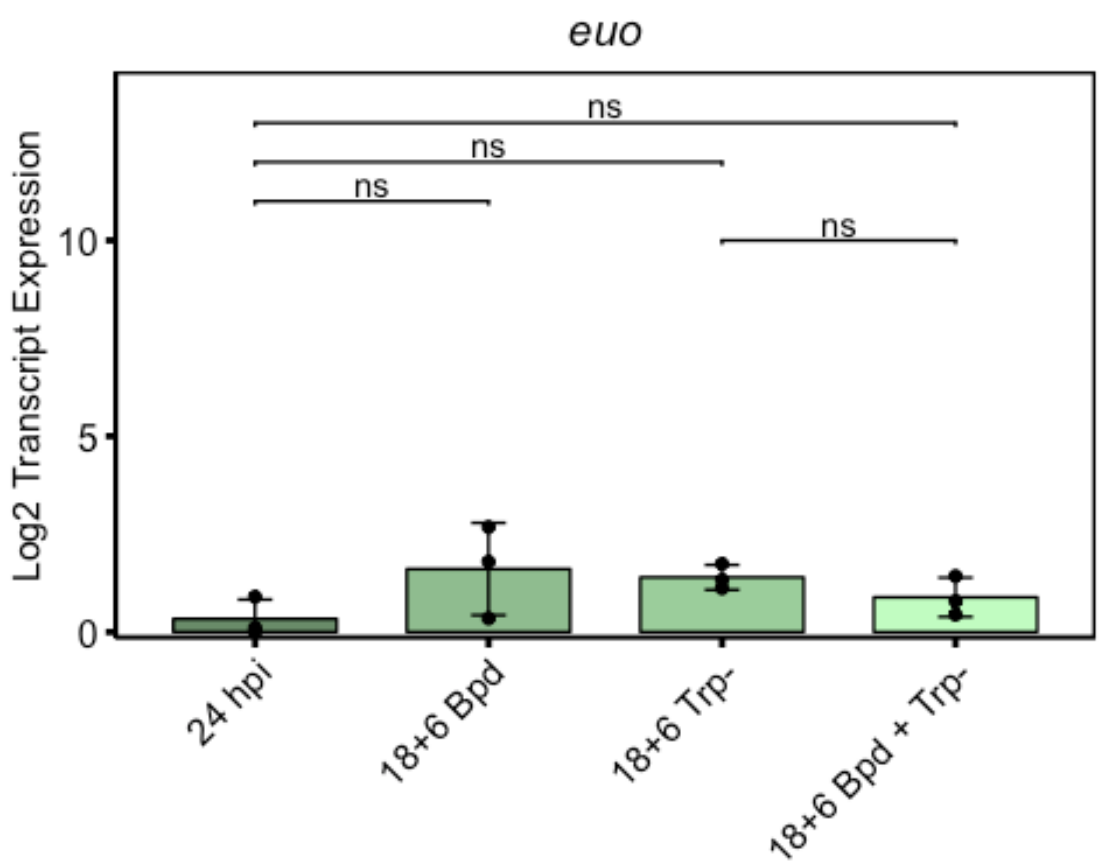


```
#omcB
omcB<-subset(rt, Gene%in%c("omcB"))
omcBaov<-aov(Log2_Value~ID, data=omcB)
summary(omcBaov)

##             Df Sum Sq Mean Sq F value Pr(>F)
## ID           3  5.173  1.7244   2.687  0.117
## Residuals    8  5.135  0.6418

TukeyHSD(x=omcBaov)

##   Tukey multiple comparisons of means
##     95% family-wise confidence level
##
## Fit: aov(formula = Log2_Value ~ ID, data = omcB)
##
## $ID
##                                  diff       lwr       upr     p adj
## 18+6 Bpd + Trp--18+6 Bpd   -1.3483693 -3.443151 0.7464121 0.2438009
## 18+6 Trp--18+6 Bpd         -1.7800503 -3.874832 0.3147311 0.0985941
## 24 hpi-18+6 Bpd            -1.0505709 -3.145352 1.0442105 0.4270064
## 18+6 Trp--18+6 Bpd + Trp- -0.4316810 -2.526462 1.6631005 0.9091369
## 24 hpi-18+6 Bpd + Trp-     0.2977984 -1.796983 2.3925798 0.9667096
## 24 hpi-18+6 Trp-           0.7294793 -1.365302 2.8242608 0.6910336
```

```
ggbarplot(omcB, x="ID2", y="Log2_Value", add="mean_sd", fill="Treatment2",
palette=c("darkseagreen4", "darkseagreen", "darkseagreen3", "darkseagreen1"),
title="omcB") + scale_y_continuous(expand=expand_scale(mult=c(0,0.1))) +
ylab(label="Log2 Transcript Expression") + rremove("legend") +
theme(axis.text.x=element_text(angle=45, hjust=1)) + rremove("xlab") +
geom_exec(geomfunc=geom_point, data=omcB, x="ID2", y="Log2_Value",
colour="black") + geom_signif(comparisons=list(c("18+6 Trp-", "18+6 Bpd +
Trp-"), c("24 hpi", "18+6 Bpd"), c("24 hpi", "18+6 Trp-"), c("24 hpi", "18+6
Bpd + Trp-")), annotation=c("ns", "ns", "ns", "ns"), y_position=c(10, 11, 12,
13)) + theme(line=element_line(size=1, colour="black"),
panel.border=element_rect(colour="black", fill="NA", size=1)) +
theme(plot.title=element_text(hjust='0.5', face="italic"))
```

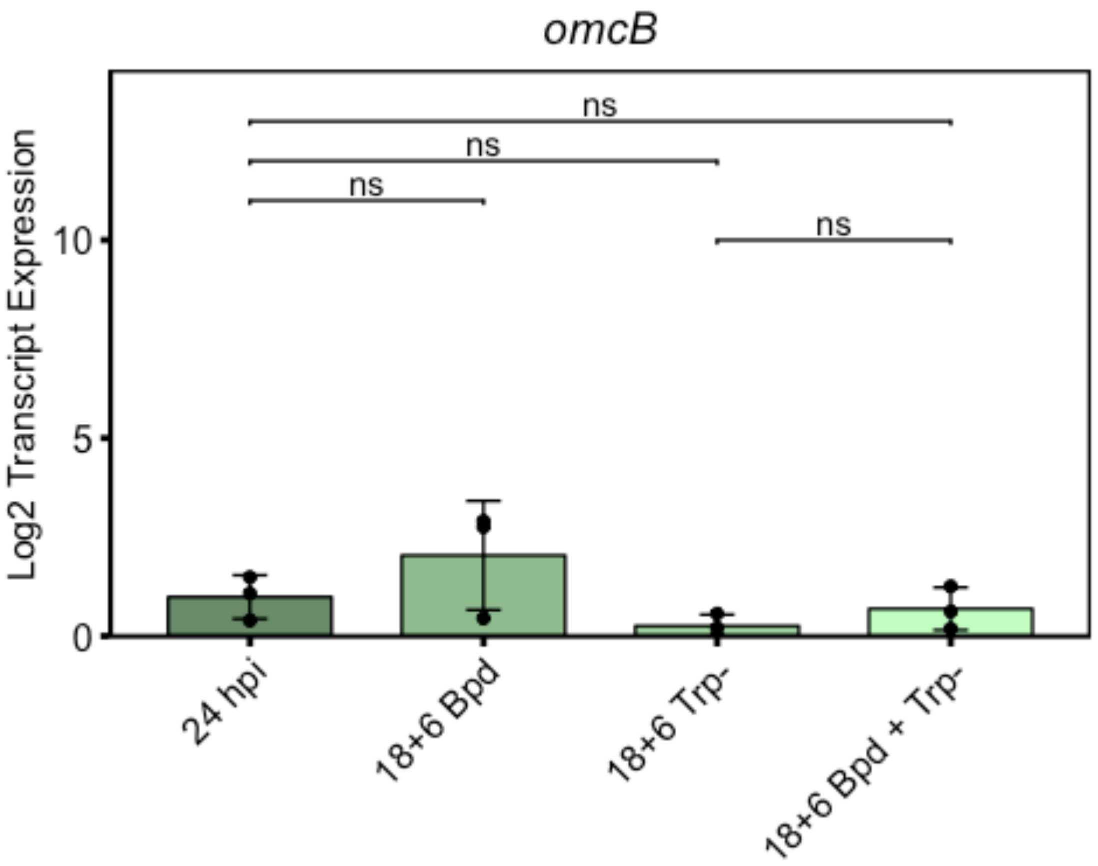

### Supplementary Figure 4

Nick Pokorzynski

4/16/2020

```
#Supplementary Figure 4a
setwd("~/Documents/Carabeo Lab/R Code/Paper 2/Supplementary Figure 4")
library(ggplot2)
library(ggpubr)

## Warning: package 'ggpubr' was built under R version 3.5.2

## Loading required package: magrittr

an<-read.csv("SuppFig4a.csv")
an$Condition2<-factor(an$Condition, levels=c("Mock", "AN"))
an$Vector2<-factor(an$Vector, levels=c("EV", "WWW", "YYF"))
anaov<-aov(OD~Condition+Vector+Condition:Vector, data=an)
summary(anaov)

##                  Df  Sum Sq Mean Sq F value   Pr(>F)    
## Condition         1 0.00020 0.00020   0.196    0.666    
## Vector            2 0.10589 0.05294  51.975 1.23e-06 ***
## Condition:Vector  2 0.08674 0.04337  42.573 3.55e-06 ***
## Residuals        12 0.01222 0.00102                     
## ---
## Signif. codes:  0 '***' 0.001 '**' 0.01 '*' 0.05 '.' 0.1 ' ' 1

TukeyHSD(x=anaov)

##   Tukey multiple comparisons of means
##     95% family-wise confidence level
## 
## Fit: aov(formula = OD ~ Condition + Vector + Condition:Vector, data = an)
## 
## $Condition
##                diff        lwr        upr     p adj
## Mock-AN 0.006666667 -0.0261149 0.03944823 0.6655779
## 
## $Vector
##                diff         lwr         upr     p adj
## WWW-EV  -0.16866667 -0.21782748 -0.11950585 0.0000026
## YYF-EV  -0.15600000 -0.20516081 -0.10683919 0.0000058
## YYF-WWW  0.01266667 -0.03649415  0.06182748 0.7751207
## 
## $`Condition:Vector`
##                           diff          lwr          upr     p adj
## Mock:EV-AN:EV      0.202666667  0.11513399  0.290199340 0.0000575
```

```
## AN:WWW-AN:EV      -0.026666667 -0.11419934  0.060866006 0.9012709
## Mock:WWW-AN:EV    -0.108000000 -0.19553267 -0.020467327 0.0132960
## AN:YYF-AN:EV      -0.004000000 -0.09153267  0.083532673 0.9999845
## Mock:YYF-AN:EV    -0.105333333 -0.19286601 -0.017800660 0.0157964
## AN:WWW-Mock:EV    -0.229333333 -0.31686601 -0.141800660 0.0000163
## Mock:WWW-Mock:EV  -0.310666667 -0.39819934 -0.223133994 0.0000006
## AN:YYF-Mock:EV    -0.206666667 -0.29419934 -0.119133994 0.0000472
## Mock:YYF-Mock:EV  -0.308000000 -0.39553267 -0.220467327 0.0000007
## Mock:WWW-AN:WWW   -0.081333333 -0.16886601  0.006199340 0.0742595
## AN:YYF-AN:WWW      0.022666667 -0.06486601  0.110199340 0.9468618
## Mock:YYF-AN:WWW   -0.078666667 -0.16619934  0.008866006 0.0878334
## AN:YYF-Mock:WWW    0.104000000  0.01646733  0.191532673 0.0172203
## Mock:YYF-Mock:WWW  0.002666667 -0.08486601  0.090199340 0.9999979
## Mock:YYF-AN:YYF   -0.101333333 -0.18886601 -0.013800660 0.0204689
```

```
p<-ggbarplot(an, x="Vector2", y="OD", add=c("mean_sd", "point"),
fill="Condition2", palette=c("orchid4", "orchid1"), title="AN3365", position
= position_dodge()) + scale_y_continuous(expand=expand_scale(mult=c(0,0.1)))
+ ylab(label="OD600") + rremove("xlab") + theme(line=element_line(size=1,
colour="black"), panel.border=element_rect(colour="black", fill="NA",
size=1)) + theme(plot.title=element_text(hjust='0.5')) +
geom_signif(y_position = c(0.9, 0.9, 0.9), xmin = c(0.75, 1.75, 2.75), xmax =
c(1.25, 2.25, 3.25), annotation = c("***", "ns", "*"), tip_length = .05) +
theme(legend.title=element_blank())
ggpar(p, legend=c("right"))
```

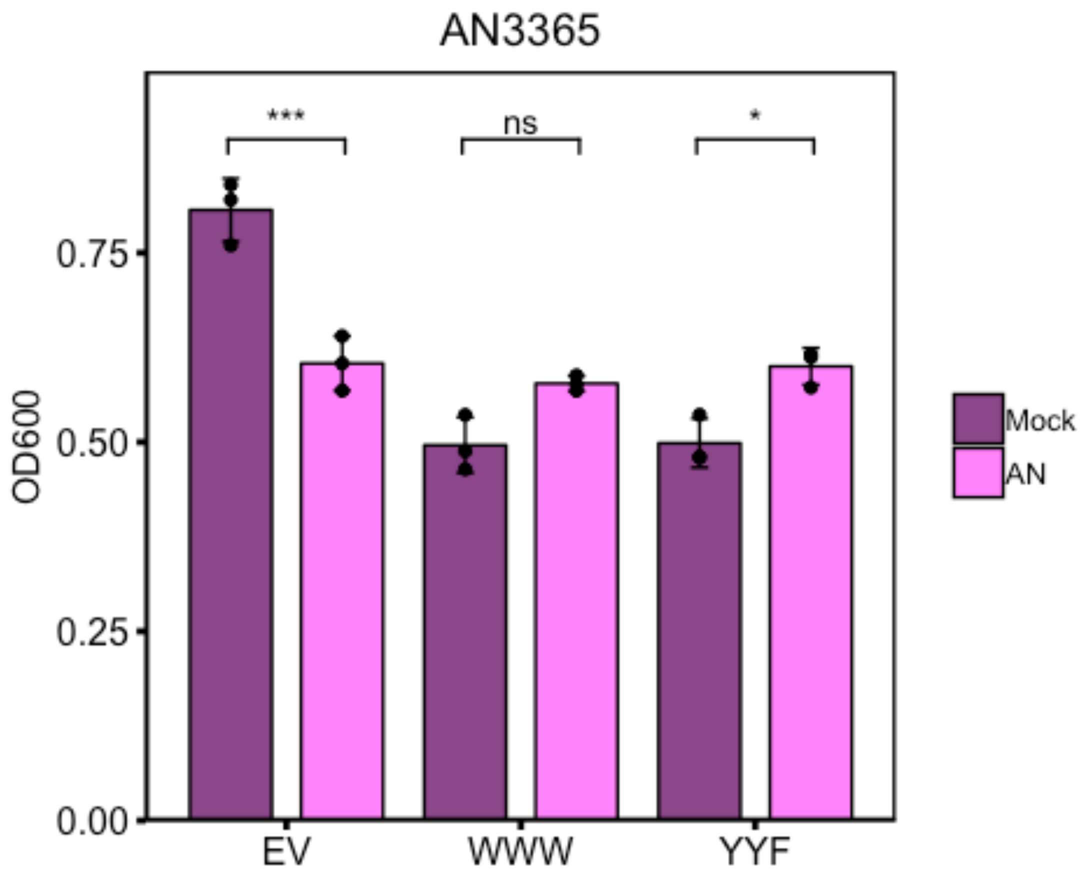


```
#Supplementary Figure 4b
setwd("~/Documents/Carabeo Lab/R Code/Paper 2/Supplementary Figure 4")
library(ggplot2)
library(ggpubr)
ind<-read.csv("SuppFig4b.csv")
ind$Condition2<-factor(ind$Condition, levels=c("Mock", "Ind"))
ind$Vector2<-factor(ind$Vector, levels=c("EV", "WWW", "YYF"))
indaov<-aov(OD~Condition+Vector+Condition:Vector, data=ind)
summary(indaov)

##                  Df Sum Sq Mean Sq F value   Pr(>F)
## Condition         1 0.0151 0.01505   3.875   0.0607 .
## Vector            2 0.5059 0.25297  65.126 2.01e-10 ***
## Condition:Vector  2 0.0484 0.02421   6.233   0.0066 **
## Residuals        24 0.0932 0.00388
## ---
## Signif. codes:  0 '***' 0.001 '**' 0.01 '*' 0.05 '.' 0.1 ' ' 1

TukeyHSD(x=indaov)

##   Tukey multiple comparisons of means
##     95% family-wise confidence level
##
## Fit: aov(formula = OD ~ Condition + Vector + Condition:Vector, data = ind)
```

```
##
## $Condition
##              diff        lwr         upr     p adj
## Mock-Ind -0.0448 -0.0917695 0.002169499 0.0606565
##
## $Vector
##                 diff        lwr        upr    p adj
## WWW-EV  -0.27400000 -0.3518210 -0.1961790 0.000000
## YYF-EV  -0.35166667 -0.4294877 -0.2738457 0.000000
## YYF-WWW -0.07766667 -0.1412072 -0.0141261 0.014593
##
## $`Condition:Vector`
##                           diff          lwr         upr     p adj
## Mock:EV-Ind:EV     0.081333333 -0.07600785  0.23867452 0.6074152
## Ind:WWW-Ind:EV    -0.167333333 -0.30359480 -0.03107187 0.0100624
## Mock:WWW-Ind:EV   -0.299333333 -0.43559480 -0.16307187 0.0000069
## Ind:YYF-Ind:EV    -0.300666667 -0.43692813 -0.16440520 0.0000064
## Mock:YYF-Ind:EV   -0.321333333 -0.45759480 -0.18507187 0.0000022
## Ind:WWW-Mock:EV   -0.248666667 -0.38492813 -0.11240520 0.0001093
## Mock:WWW-Mock:EV  -0.380666667 -0.51692813 -0.24440520 0.0000001
## Ind:YYF-Mock:EV   -0.382000000 -0.51826146 -0.24573854 0.0000001
## Mock:YYF-Mock:EV  -0.402666667 -0.53892813 -0.26640520 0.0000000
## Mock:WWW-Ind:WWW  -0.132000000 -0.24325702 -0.02074298 0.0136221
## Ind:YYF-Ind:WWW   -0.133333333 -0.24459035 -0.02207631 0.0124883
## Mock:YYF-Ind:WWW  -0.154000000 -0.26525702 -0.04274298 0.0031435
## Ind:YYF-Mock:WWW  -0.001333333 -0.11259035  0.10992369 1.0000000
## Mock:YYF-Mock:WWW -0.022000000 -0.13325702  0.08925702 0.9891382
## Mock:YYF-Ind:YYF  -0.020666667 -0.13192369  0.09059035 0.9918274

p<-ggbarplot(ind, x="Vector2", y="OD", add=c("mean_sd", "point"),
fill="Condition2", palette=c("mediumpurple4", "mediumpurple1"),
title="Indolmycin", position = position_dodge()) +
scale_y_continuous(expand=expand_scale(mult=c(0,0.1))) + ylab(label="OD600")
+ rremove("xlab") + theme(line=element_line(size=1, colour="black"),
panel.border=element_rect(colour="black", fill="NA", size=1)) +
theme(plot.title=element_text(hjust='0.5')) + geom_signif(y_position = c(1.1,
1.1, 1.1), xmin = c(0.75, 1.75, 2.75), xmax = c(1.25, 2.25, 3.25), annotation
= c("ns", "*", "ns"), tip_length = .05) + theme(legend.title=element_blank())
ggpar(p, legend=c("right"))
```

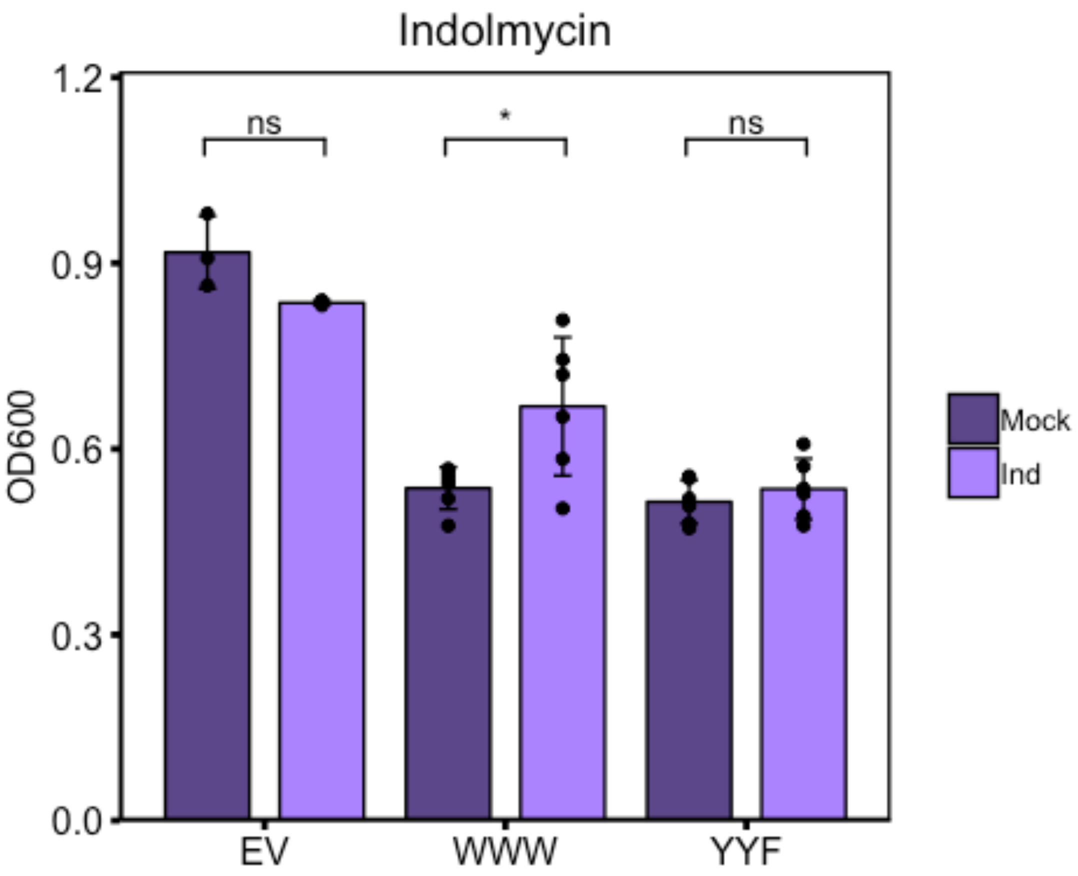
Indolmycin
OD600
1.2
0.9
0.6
0.3
0.0
ns
*
ns
EV
WWW
YYF
Mock
Ind

### Supplementary Figure 5

Nick Pokorzynski

4/17/2020

```
#Supplementary Figure 5
setwd("~/Documents/Carabeo Lab/R Code/Paper 2/Supplementary Figure 5")
library(ggplot2)
library(ggpubr)

## Warning: package 'ggpubr' was built under R version 3.5.2

## Loading required package: magrittr

x<-read.csv("SuppFig5.csv")
OmpA<-subset(x, Gene%in%c("OmpA"))
YtgCR<-subset(x, Gene%in%c("YtgCR"))
t.test(Value~Treatment, data=OmpA)

##
##  Welch Two Sample t-test
##
## data:  Value by Treatment
## t = 3.2866, df = 2.0179, p-value = 0.08046
## alternative hypothesis: true difference in means is not equal to 0
## 95 percent confidence interval:
##  -0.006064588  0.046760690
## sample estimates:
##   mean in group 24 hpi mean in group 24h -Trp
##              0.4101351              0.3897871

t.test(Value~Treatment, data=YtgCR)

##
##  Welch Two Sample t-test
##
## data:  Value by Treatment
## t = -0.51273, df = 2.9582, p-value = 0.644
## alternative hypothesis: true difference in means is not equal to 0
## 95 percent confidence interval:
##  -0.269057  0.194904
## sample estimates:
##   mean in group 24 hpi mean in group 24h -Trp
##              0.9873781              1.0244546

ggbarplot(YtgCR, x="Treatment", y="Value", add=c("mean_sd", "point"),
color="black", fill="Treatment", palette=c("steelblue4", "steelblue2"),
title="YtgCR") + scale_y_continuous(expand=expand_scale(mult=c(0,0.1))) +
```

```
ylab(label="3'-to-5' Ratio") + rremove("legend") +
theme(axis.text.x=element_text(angle=45, hjust=1)) + rremove("xlab") +
theme(line=element_line(size=1, colour="black"),
panel.border=element_rect(colour="black", fill="NA", size=1)) +
theme(plot.title=element_text(hjust='0.5')) +
geom_signif(comparisons=list(c("24 hpi", "24h -Trp")), annotation="ns",
y_position=1.25)
```

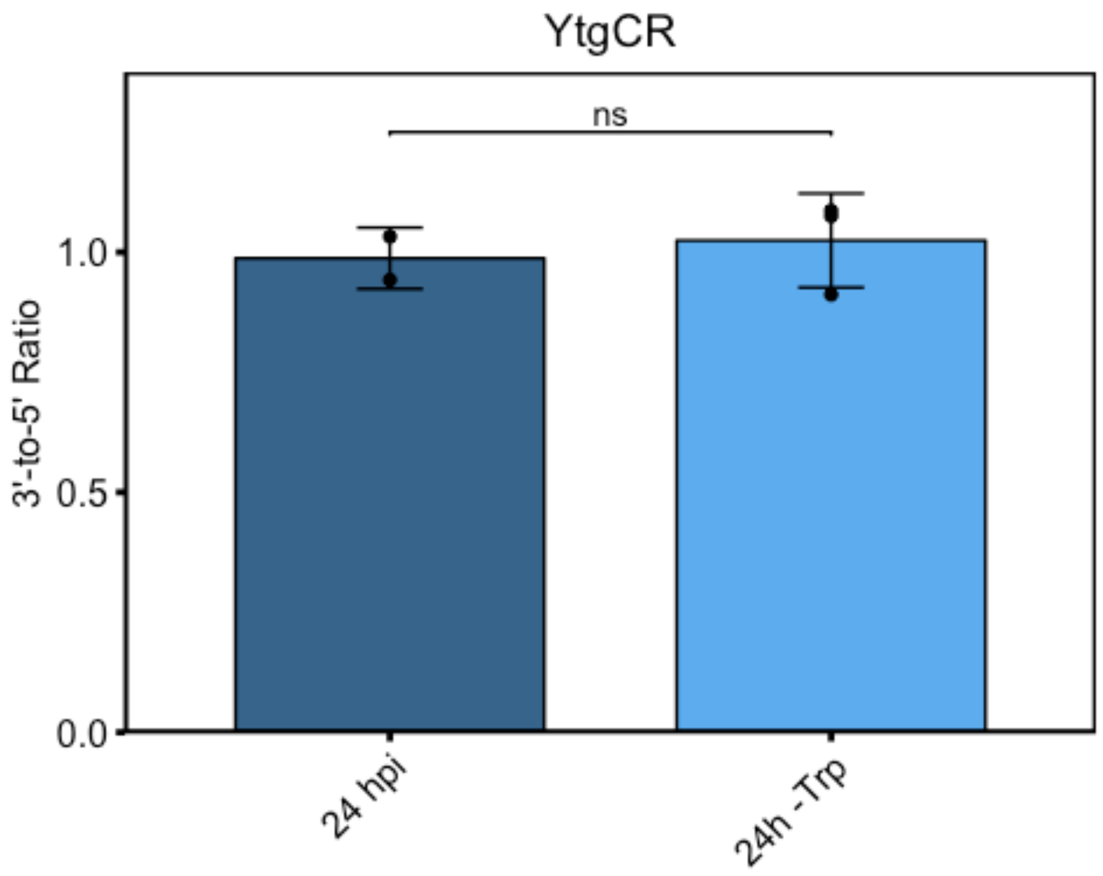


```
ggbarplot(OmpA, x="Treatment", y="Value", add=c("mean_sd", "point"),
color="black", fill="Treatment", palette=c("steelblue4", "steelblue2"),
title="OmpA") + scale_y_continuous(expand=expand_scale(mult=c(0,0.1))) +
ylab(label="3'-to-5' Ratio") + rremove("legend") +
theme(axis.text.x=element_text(angle=45, hjust=1)) + rremove("xlab") +
theme(line=element_line(size=1, colour="black"),
panel.border=element_rect(colour="black", fill="NA", size=1)) +
theme(plot.title=element_text(hjust='0.5')) +
geom_signif(comparisons=list(c("24 hpi", "24h -Trp")), annotation="ns",
y_position=1.25)
```

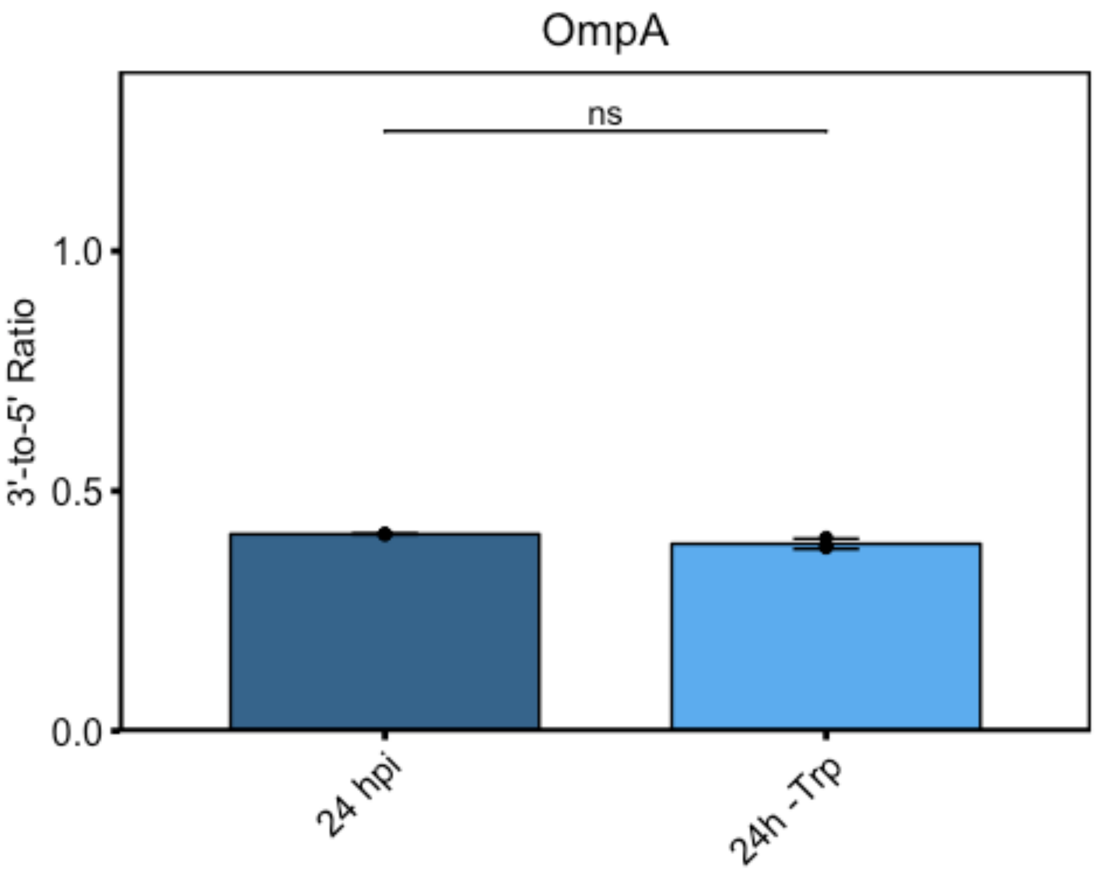
OmpA
ns
3'-to-5' Ratio
1.0
0.5
0.0
24 hpi
24h -Trp

### Supplementary Figure 6

Nick Pokorzynski

4/17/2020

```
#Supplementary Figure 6
setwd("~/Documents/Carabeo Lab/R Code/Paper 2/Supplementary Figure 6")
library(ggplot2)
library(ggpubr)

## Warning: package 'ggpubr' was built under R version 3.5.2

## Loading required package: magrittr

x<-read.csv("SuppFig6.csv")
#subset values for each gene for statistical analysis
CTL0174<-subset(x, Gene%in%c("CTL0174"))
GroEL_1<-subset(x, Gene%in%c("GroEL_1"))
OmpA<-subset(x, Gene%in%c("OmpA"))
YtgD<-subset(x, Gene%in%c("YtgD:YtgA"))
YtgCR<-subset(x, Gene%in%c("YtgCR"))
#compute t-test w/ Welch's correction for unequal variance
t.test(Value~Treatment, data=CTL0174)

##
##  Welch Two Sample t-test
##
## data:  Value by Treatment
## t = 5.5583, df = 3.4607, p-value = 0.00778
## alternative hypothesis: true difference in means is not equal to 0
## 95 percent confidence interval:
##  0.1221562 0.3996149
## sample estimates:
## mean in group Bic mean in group UTD
##         0.9588099         0.6979244

t.test(Value~Treatment, data=GroEL_1)

##
##  Welch Two Sample t-test
##
## data:  Value by Treatment
## t = 9.3409, df = 3.5373, p-value = 0.001289
## alternative hypothesis: true difference in means is not equal to 0
## 95 percent confidence interval:
##  0.1235900 0.2363191
## sample estimates:
```

```
## mean in group Bic mean in group UTD
##         0.8887219         0.7087673

t.test(Value~Treatment, data=OmpA)

##
##  Welch Two Sample t-test
##
## data:  Value by Treatment
## t = 2.1061, df = 3.8481, p-value = 0.1057
## alternative hypothesis: true difference in means is not equal to 0
## 95 percent confidence interval:
##  -0.01965513  0.13558747
## sample estimates:
## mean in group Bic mean in group UTD
##         0.5135750         0.4556089

t.test(Value~Treatment, data=YtgD)

##
##  Welch Two Sample t-test
##
## data:  Value by Treatment
## t = 8.5129, df = 2.8011, p-value = 0.004398
## alternative hypothesis: true difference in means is not equal to 0
## 95 percent confidence interval:
##  0.1921529 0.4371524
## sample estimates:
## mean in group Bic mean in group UTD
##         0.4271481         0.1124954

t.test(Value~Treatment, data=YtgCR)

##
##  Welch Two Sample t-test
##
## data:  Value by Treatment
## t = -0.49111, df = 2.0347, p-value = 0.6712
## alternative hypothesis: true difference in means is not equal to 0
## 95 percent confidence interval:
##  -0.5675118  0.4495186
## sample estimates:
## mean in group Bic mean in group UTD
##         0.8065918         0.8655884

#factor "Treatment" levels
x$Treatment2<-factor(x$Treatment, levels=c("UTD", "Bic"))
#factor "Gene" levels
x$Gene2<-factor(x$Gene, levels=c("GroEL_1", "OmpA", "CTL0174", "YtgD:YtgA",
"YtgCR"))
#barplot faceted by Gene with Welch's t-test for unequal variance
```

```
ggbarplot(x, x="Treatment2", y="Value", add="mean_sd", color="black",
fill="Treatment2", palette=c("firebrick4", "firebrick3"), facet.by="Gene2",
ncol=5) + scale_y_continuous(expand=expand_scale(mult=c(0,0.1))) +
ylab(label="3'-to-5' Ratio") + rremove("legend") +
theme(axis.text.x=element_text(angle=45, hjust=1)) + rremove("xlab") +
geom_exec(geomfunc=geom_point, data=x, x="Treatment2", y="Value",
colour="black") + stat_compare_means(comparisons=list(c("UTD", "Bic")),
label="p.signif", method="t.test", label.y=1.25)
```

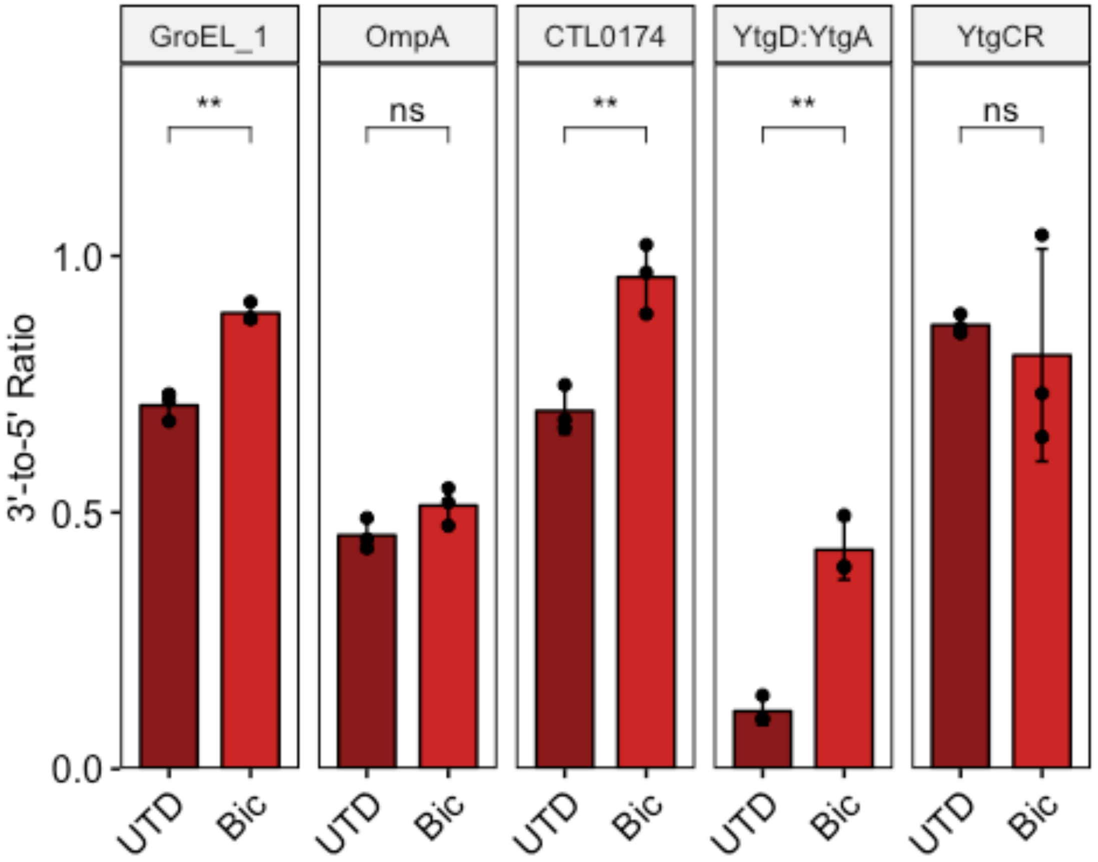
