## Supplementary Table 1 for "Regulation of an iron-dependent repressor by tryptophan availability attenuates transcription of the tryptophan salvage genes in *Chlamydia trachomatis*"

**Supplementary Table 1. Oligonucleotide primers used in this study**

| Primer ID | Nucleotide Sequence (5' to 3') | Application | Figure | Reference |
|---|---|---|---|---|
| qtrpR_Fwd | CGAACGTAAAGATGTCGCTTC | RT-qPCR | Fig. 1D-E, Fig. 5D | Pokorzynski *et al.,* 2019 |
| qtrpR_Rev | AAGGGCATTAGATCCTCTGGTAA | RT-qPCR | Fig. 1D-E, Fig. 5D | Pokorzynski *et al.,* 2019 |
| qtrpB_Fwd | GAGGTGGCTCCAACGCTATTG | RT-qPCR | Fig. 1D-E, Fig. 5D | Brinkworth *et al.,* 2018 |
| qtrpB_Rev | TCCAGCGGAAATGGAGTGAG | RT-qPCR | Fig. 1D-E, Fig. 5D | Brinkworth *et al.,* 2018 |
| qtrpA_Fwd | AGCTCTGATTCAAGGAGGTGTTG | RT-qPCR | Fig. 1D-E, Fig. 5D | Brinkworth *et al.,* 2018 |
| qtrpA_Rev | AGTCCCTTTGTAGAAGCGGATTG | RT-qPCR | Fig. 1D-E, Fig. 5D | Brinkworth *et al.,* 2018 |
| qdnaB_Fwd | TGTGCTGGGGTGTATGTTGACCA | RT-qPCR | Fig. S1 | Pokorzynski *et al.,* 2019 |
| qdnaB_Rev | ATGGGGCGGTCAGACTTGA | RT-qPCR | Fig. S1 | Pokorzynski *et al.,* 2019 |
| qeuo_Fwd | GCTGTTCCTGTTACTTCGCAAA | RT-qPCR; qPCR | Fig. 1B-C, Fig. S2B-C | Thompson & Carabeo, 2011 |
| qeuo_Rev | AACATAGATAGCCTGACGAGTCACA | RT-qPCR; qPCR | Fig. 1B-C, Fig. S2B-C | Thompson & Carabeo, 2011 |
| qomcB_Fwd | CCAAAGCGAAAGACAACACTTCT | RT-qPCR | Fig. 1C, Fig. S2C | Thompson & Carabeo, 2011 |
| qomcB_Rev | AACCGGAGCAACCTCTTTACG | RT-qPCR | Fig. 1C, Fig. S2C | Thompson & Carabeo, 2011 |
| qytgA_Fwd | CTCTTGTGTTTAGCAGGCTGTTTC | RT-qPCR | Fig. S1 | Thompson & Carabeo, 2011 |
| qytgA_Rev | TGCGATTCATAGACAAGACATAGATG | RT-qPCR | Fig. S1 | Thompson & Carabeo, 2011 |
| qahpC_Fwd | CCAGTTAGCTGGACAAACCATTCCG | RT-qPCR | Fig. S1 | Thompson & Carabeo, 2011 |
| qahpC_Rev | CGTTCCATTGACGAGGAATTGCGT | RT-qPCR | Fig. S1 | Thompson & Carabeo, 2011 |
| qdevB_Fwd | ACGAAGATGTAGAAGCTGGAAGTA | RT-qPCR | Fig. S1 | Thompson & Carabeo, 2011 |
| qdevB_Rev | TGCGGTATCCATACGAAAGATTTG | RT-qPCR | Fig. S1 | Thompson & Carabeo, 2011 |
| qNrdA_Fwd | GAGAAGAGGACGGGAGTACAGA | RT-qPCR | Fig. S1 | Brinkworth *et al.,* 2018 |
| qNrdA_Rev | CTACTTCAGACTCTTTGATGCCG | RT-qPCR | Fig. S1 | Brinkworth *et al.,* 2018 |
| qNrdB_Fwd | TCAGCACCGCAGAGAGCTTG | RT-qPCR | Fig. S1 | Brinkworth *et al.,* 2018 |
| qNrdB_Rev | ATCGCAGCACGCTCGTTATAG | RT-qPCR | Fig. S1 | Brinkworth *et al.,* 2018 |
| qYtgCR-5'_Fwd | TTTGGCAGTTTCGCTGATTTG | RT-qPCR | Fig.4B-C, Fig. S5, Fig. S6 | This Study |
| qYtgCR-5'_Rev | GCATGAGAAAGGCTCTCACTTA | RT-qPCR | Fig.4B-C, Fig. S5, Fig. S6 | This Study |
| qYtgCR-3'_Fwd | CCAGACAGCTTTCGGATAGATT | RT-qPCR | Fig.4B-C, Fig. S5, Fig. S6 | This Study |
| qYtgCR-3'_Rev | TAGGCTTCCCTTCCACTACA | RT-qPCR | Fig.4B-C, Fig. S5, Fig. S6 | This Study |
| qCTL0174-5'_Fwd | GATAGGTCGCGATTCAAGGAG | RT-qPCR | Fig. 4B-C, Fig. S6 | This Study |
| qCTL0174-5'_Rev | GCGATTCCGCTATTCTGTCTAT | RT-qPCR | Fig. 4B-C, Fig. S6 | This Study |
| qCTL0174-3'_Fwd | GAATCGCTGAAACCCTAAGTAGA | RT-qPCR | Fig. 4B-C, Fig. S6 | This Study |
| qCTL0174-3'_Rev | CGCCTTCCTTCGTGGATTAT | RT-qPCR | Fig. 4B-C, Fig. S6 | This Study |
| qGroEL_1-5'_Fwd | GAGACGGAACTACAACAGCTAC | RT-qPCR | Fig. 4B-C, Fig. S6 | This Study |
| qGroEL_1-5'_Rev | CGTTTGAGGTCCATTGGATTTG | RT-qPCR | Fig. 4B-C, Fig. S6 | This Study |
| qGroEL_1-3'_Fwd | CGAAGAGTTGGGCATGAAATTAG | RT-qPCR | Fig. 4B-C, Fig. S6 | This Study |
| qGroEL_1-3'_Rev | ACGATGGTCGTGTCTTCTTTAG | RT-qPCR | Fig. 4B-C, Fig. S6 | This Study |
| qOmpA-5'_Fwd | GGTATTAGTGTTTGCCGCTTTG | RT-qPCR | Fig.4B-C, Fig. S5, Fig. S6 | This Study |
| qOmpA-5'_Rev | GCCGAAACCTTCCCATAGAA | RT-qPCR | Fig.4B-C, Fig. S5, Fig. S6 | This Study |
| qOmpA-3'_Fwd | ACGCTCAATCCAAGCCTAAA | RT-qPCR | Fig.4B-C, Fig. S5, Fig. S6 | This Study |
| qOmpA-3'_Rev | AGGGAATTCTTGCCCTACATATC | RT-qPCR | Fig.4B-C, Fig. S5, Fig. S6 | This Study |
| qYtgA-5'_Fwd | CCCGCTGATGAGAGCATCTA | RT-qPCR | Fig. 4C, Fig. S6 | This Study |
| qYtgA-5'_Rev | TCGCTCCATCAATCAGAACAA | RT-qPCR | Fig. 4C, Fig. S6 | This Study |
| qYtgD-3'_Fwd | GAAGAAGCAAGCACAGCTTTAG | RT-qPCR | Fig. 4C, Fig. S6 | This Study |
| qYtgD-3'_Rev | CATCCGCATTACCAATGACAAG | RT-qPCR | Fig. 4C, Fig. S6 | This Study |
| pET151-YtgCR-3xFLAG_Fwd | CACCATGCTGAGTTGTATATTTCAGGACACTATCTTTCT | Cloning | Fig. 3 | This Study |
| pET151-YtgCR-3xFLAG_Rev | CTATTTATCGTCATCATCCTTATAGTCCTTGTCGTCATCGTCCTT | Cloning | Fig. 3 | This Study |
| pBOMBL-YtgCR-FLAG_Fwd | AAAGATCTTCACACAGGACATCTGCATGCTGAGTTGTATATTTCAGGACAC | Cloning | Fig. 2C-D, Fig. S3 | This Study |
| pBOMBL-YtgCR-FLAG_Rev | ACATATTTGAATGGTCGACCGGTACTTACTTATCGTCGTCATCCTTGTAGTCGCAACCATCCGACTTCCTTG | Cloning | Fig. 2C-D, Fig. S3 | This Study |
| RACE_trpB_GSP | GATTACGCCAAGCTTAGCTCGTAACGCCTCTTCATCGGTGGCT | 5'-RACE | Fig. 5A | Pokorzynski *et al.,* 2019 |
| RACE_trpB_nGSP | GATTACGCCAAGCTTCGTGGAATACTCCAGGTCGCCCTGTTGC | 5'-RACE | Fig. 5A | Pokorzynski *et al.,* 2019 |
| M13_Fwd | TGTAAAACGACGGCCAGT | 5'-RACE | Fig. 5B-C | Addgene |
| M13_Rev | CAGGAAACAGCTATGAC | 5'-RACE | Fig. 5B-C | Addgene |
