## Supplementary Table 2 for "Regulation of an iron-dependent repressor by tryptophan availability attenuates transcription of the tryptophan salvage genes in *Chlamydia trachomatis*"

**Supplementary Table 2. Plasmids used in this study**

| Plasmid | Relevant Details | Application | Figure |
|---|---|---|---|
| pET151-EV | Circularized empty vector | Expression of YtgCR in *E. coli* | Fig. 3 |
| pET151-YtgCR(WWW)-3xFLAG | Expression of YtgCR with intact WWW motif and C-terminal 3xFLAG epitope w/ N-terminal 6xHis and V5 tags | Expression of YtgCR in *E. coli* | Fig. 3 |
| pET151-YtgCR(YYF)-3xFLAG | Expression of YtgCR with WWW motif mutated to YYF and C-terminal 3xFLAG epitope w/ N-terminal 6xHis and V5 tags | Expression of YtgCR in *E. coli* | Fig. 3 |
| pBOMBL-YtgCR(WWW)-FLAG | Expression of YtgCR in *C. trachomatis* with intact WWW motif and C-terminal FLAG epitope | Expression of YtgCR in *C. trachomatis* | Fig. 2C-D, Fig. S3 |
| pBOMBL-YtgCR(YYF)-FLAG | Expression of YtgCR in *C. trachomatis* with WWW motif mutated to YYF and C-terminal FLAG epitope | Expression of YtgCR in *C. trachomatis* | Fig. 2C-D, Fig. S3 |
| pRACE | Sequencing vector for RACE products | RACE Sequencing | Fig. 5B-C |
