## Supplementary figures and images for "Regulation of an iron-dependent repressor by tryptophan availability attenuates transcription of the tryptophan salvage genes in *Chlamydia trachomatis*"

### Figure 1

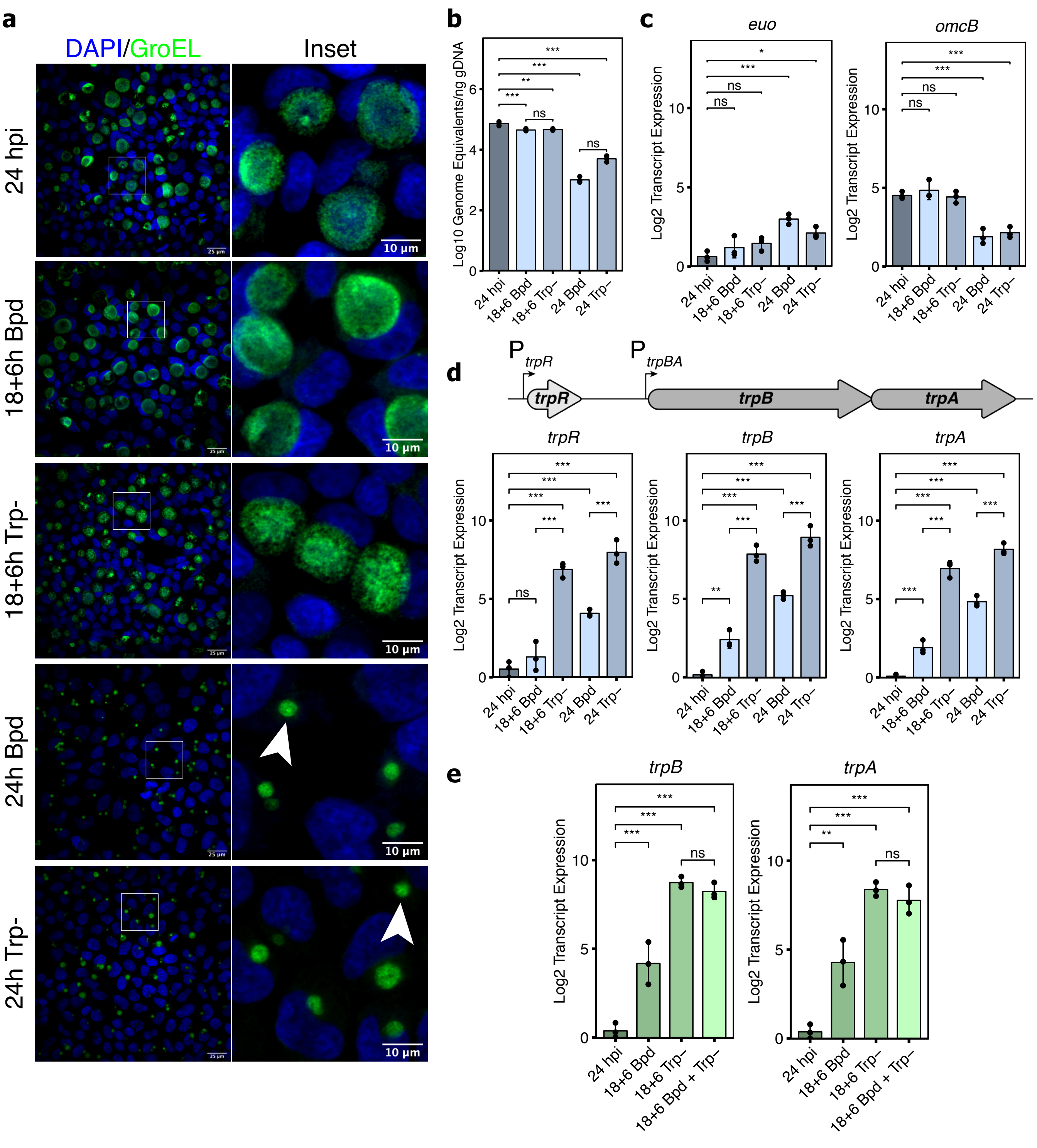

### Figure 2

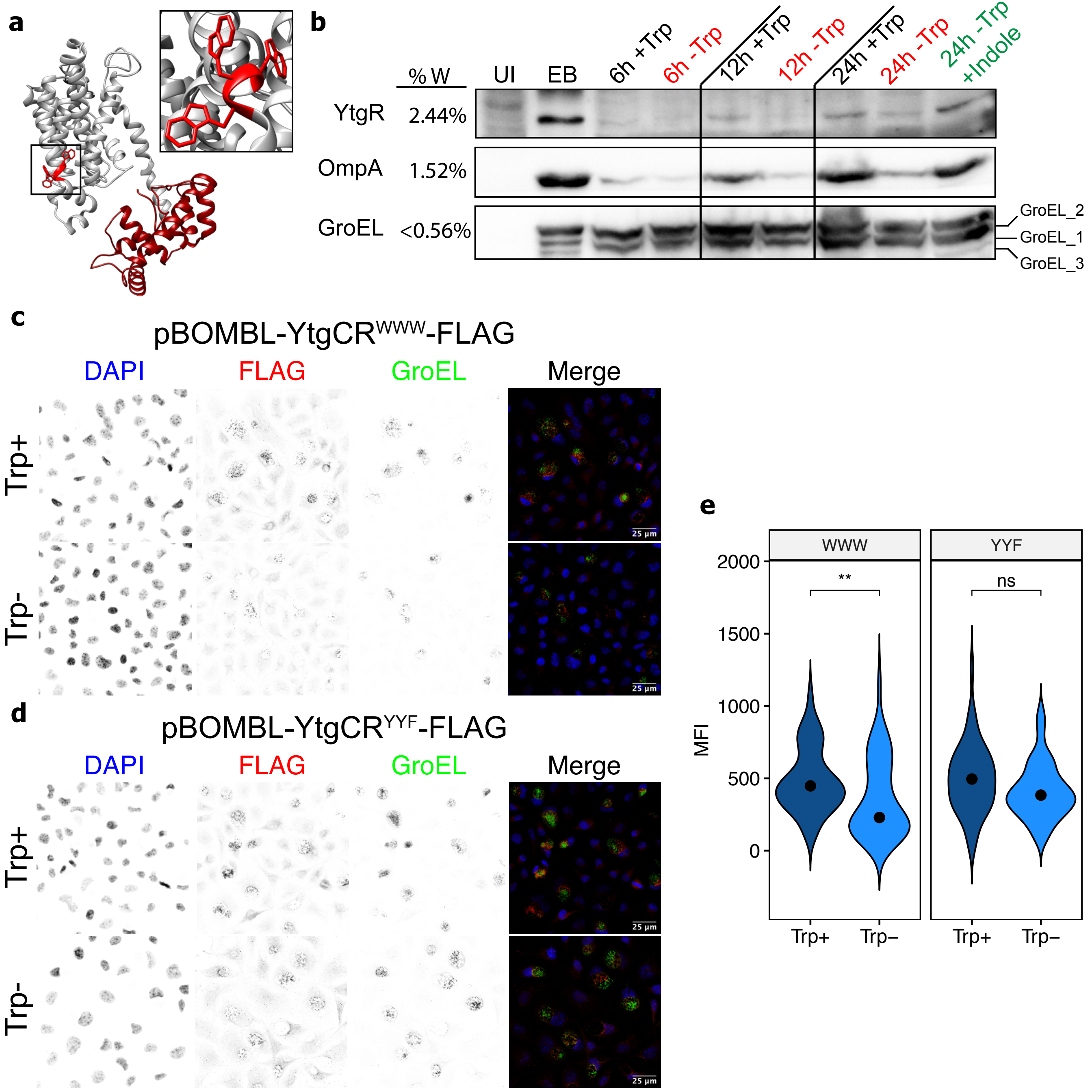

### Figure 3

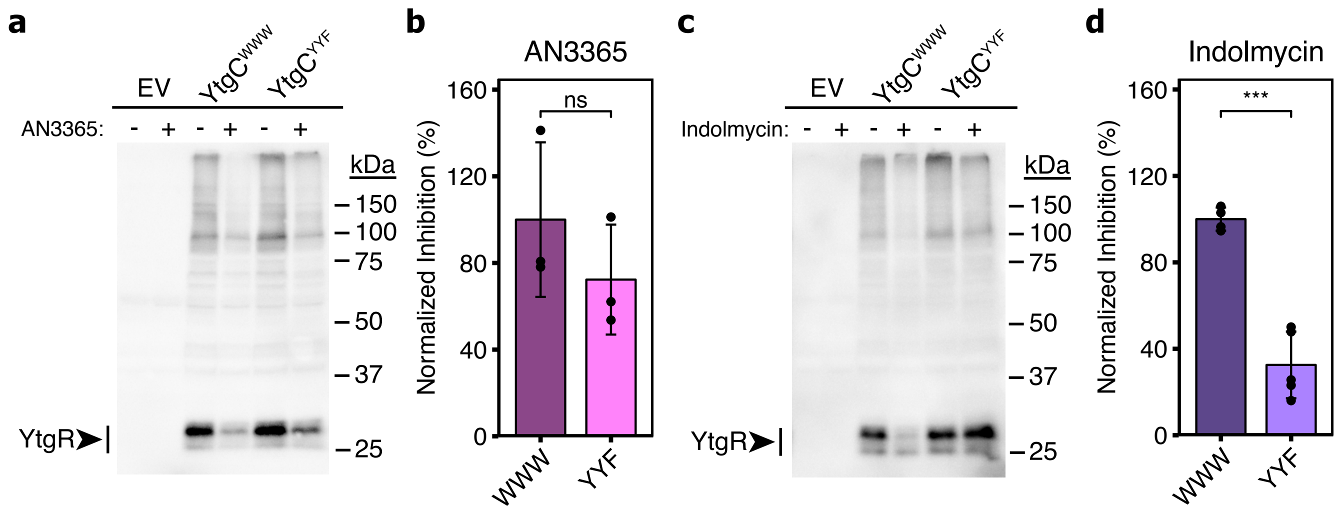

### Figure 4

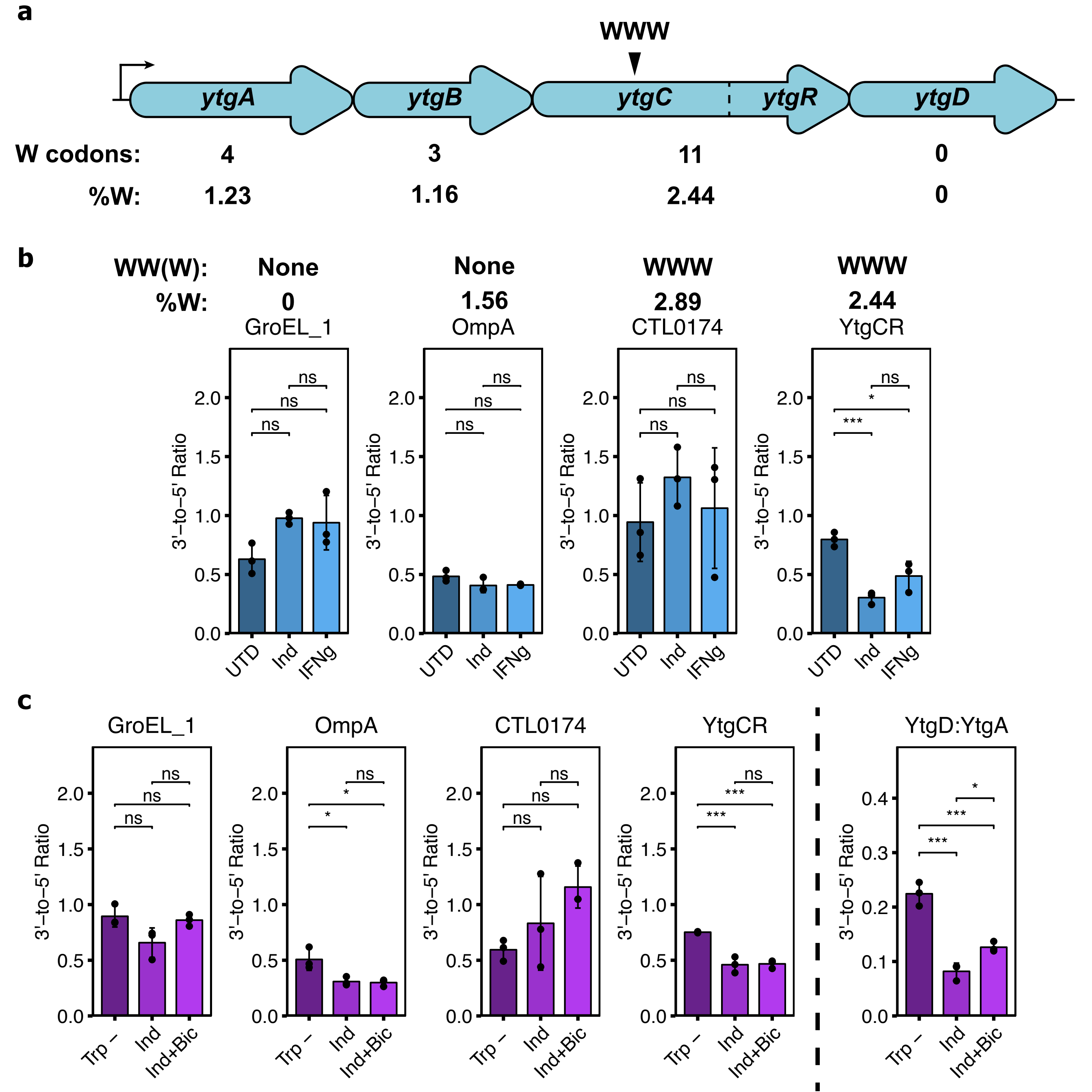

### Figure 5

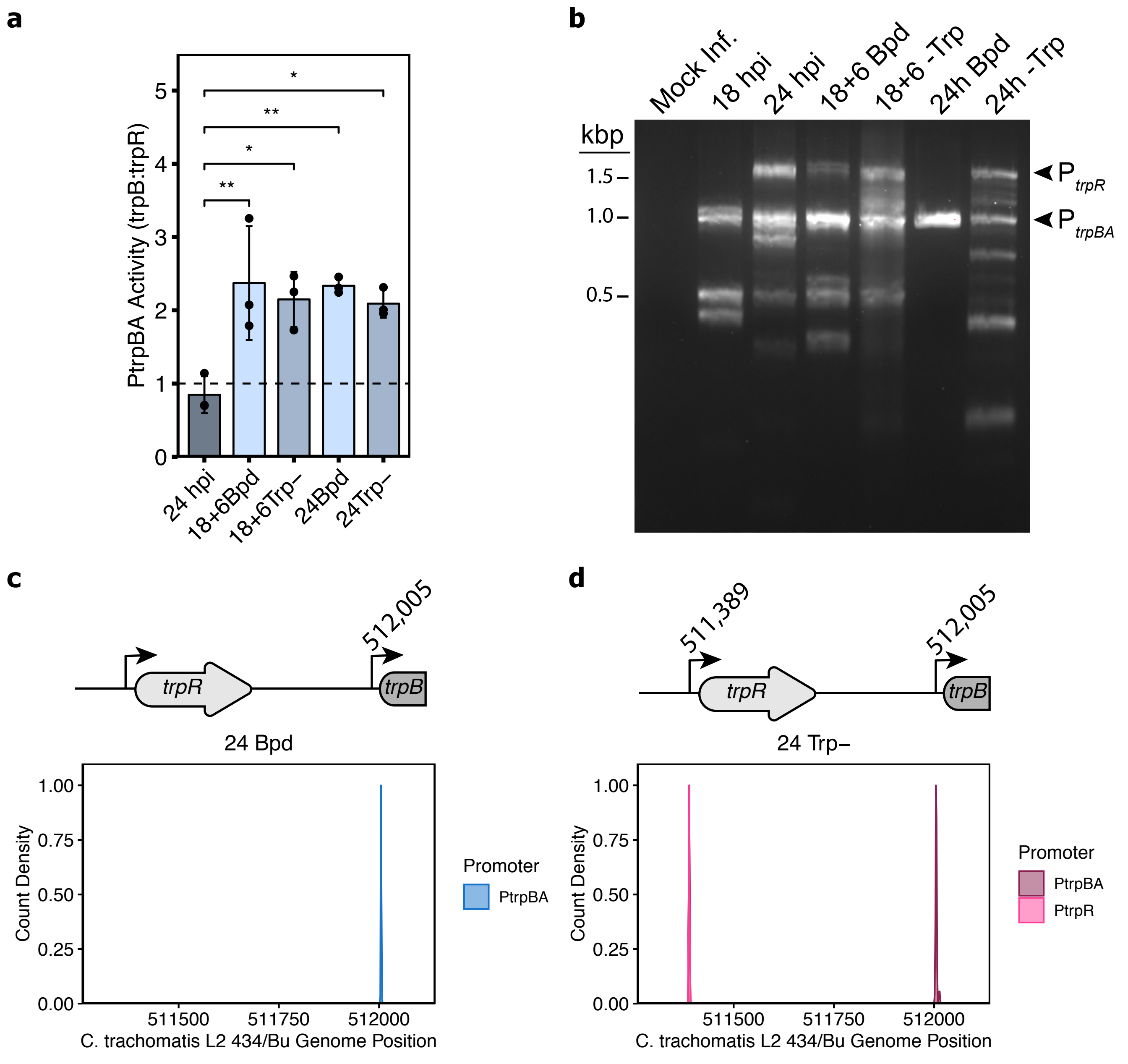

### Figure 6

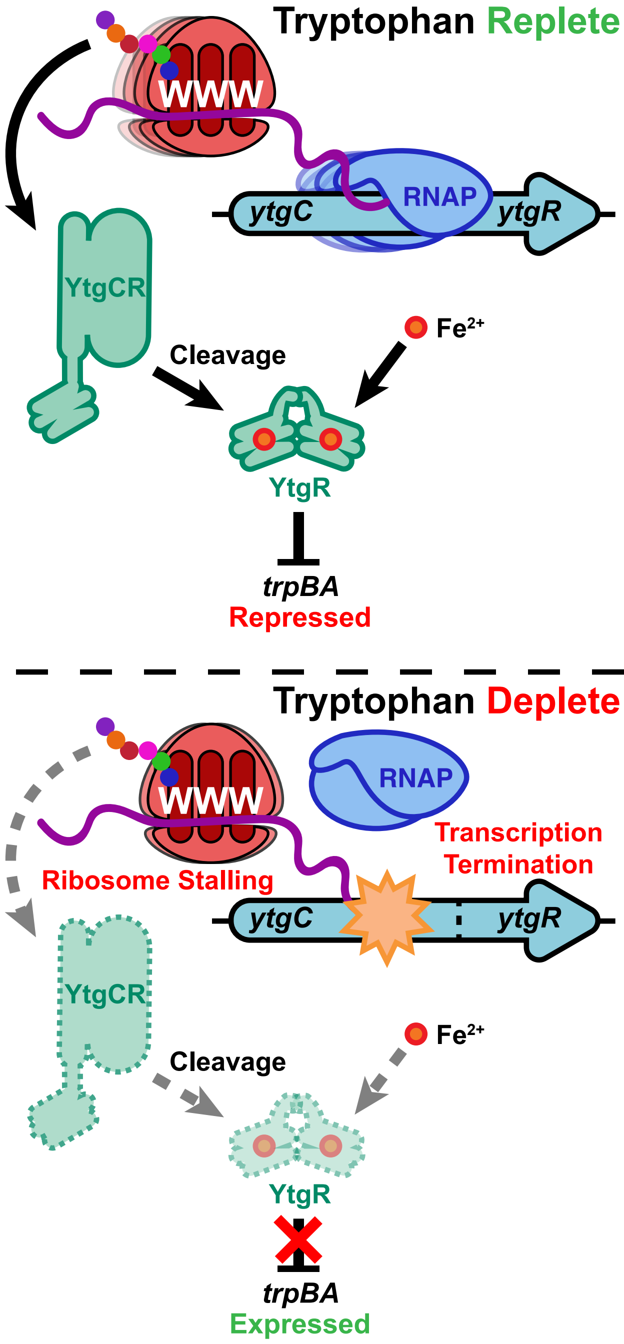

### Supplementary Figure 1

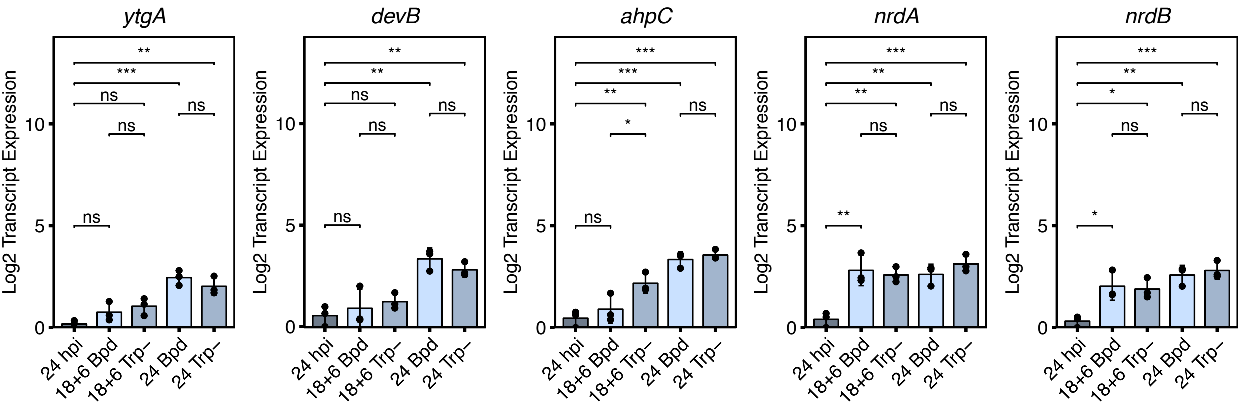

### Supplementary Figure 2

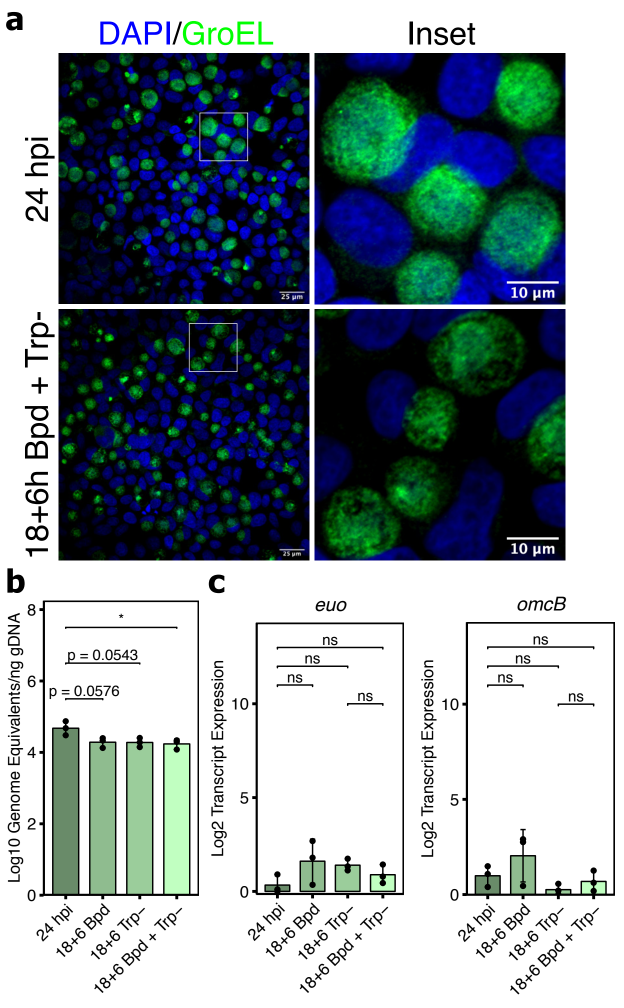

### Supplementary Figure 3

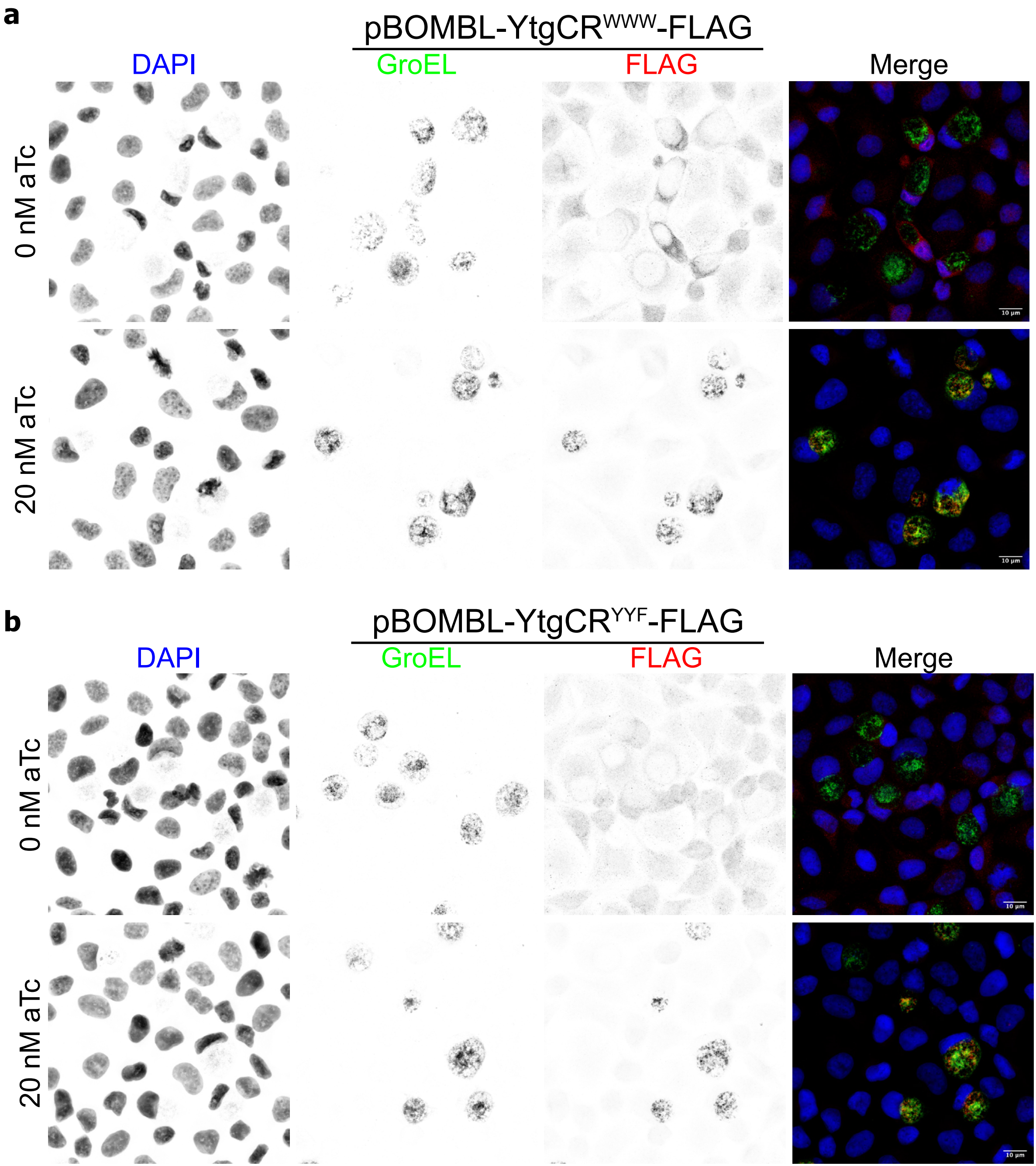

### Supplementary Figure 4

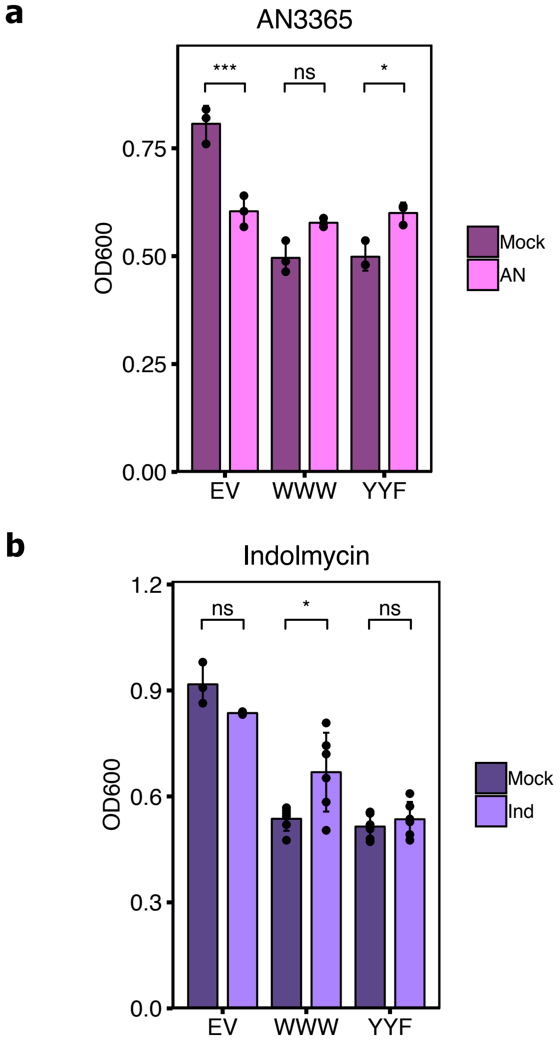

### Supplementary Figure 5

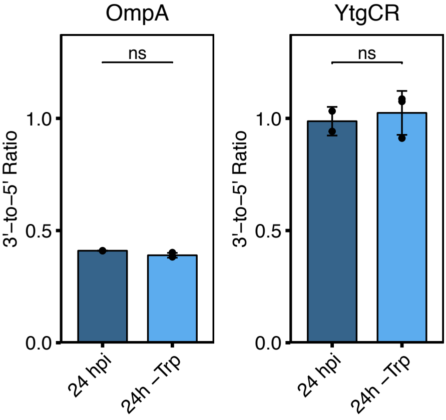

### Supplementary Figure 6

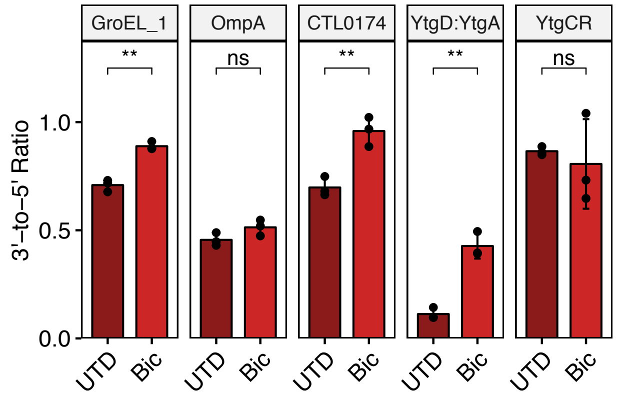

### Supplementary Figure 8

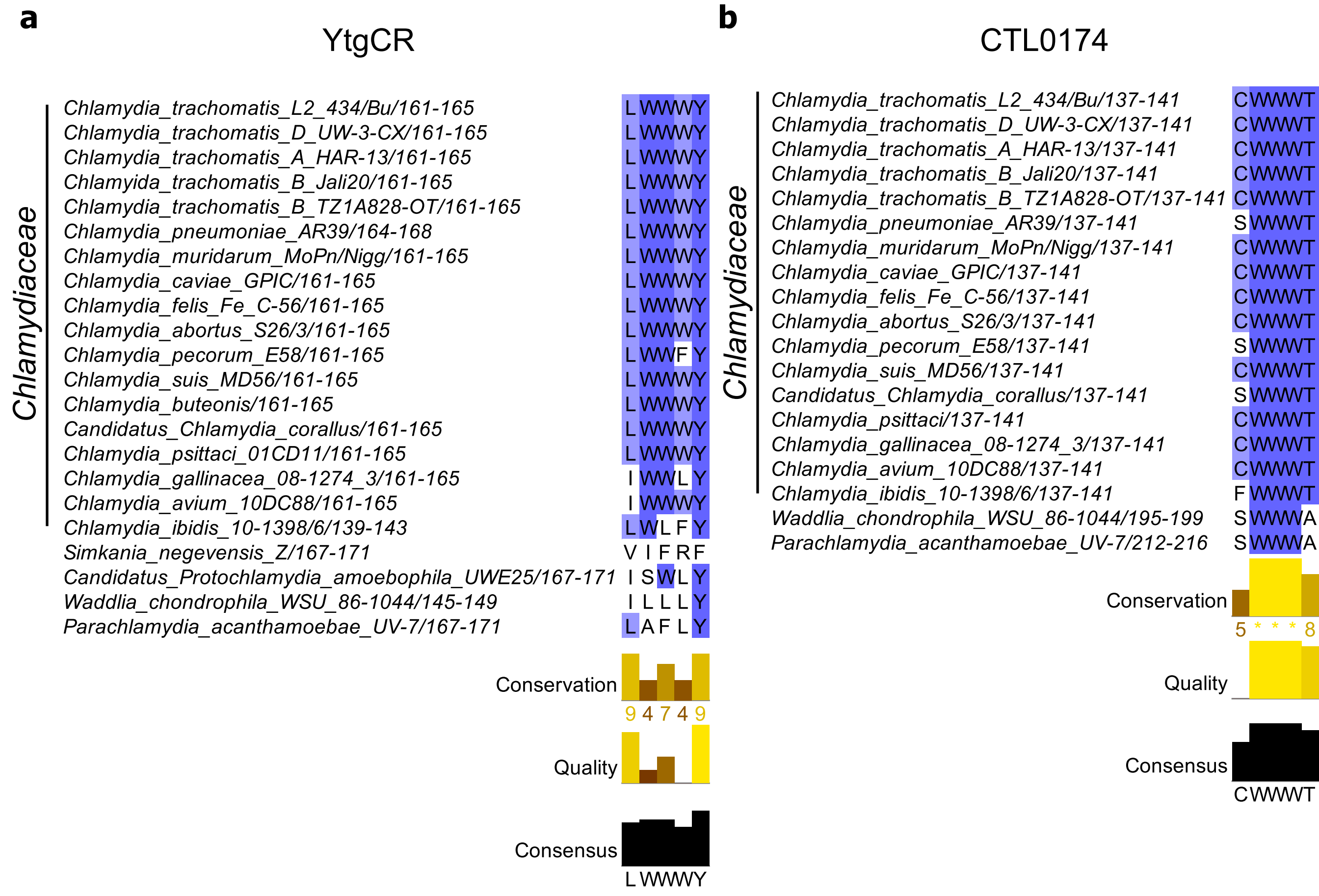
