## Supplementary Figure 7 for "Regulation of an iron-dependent repressor by tryptophan availability attenuates transcription of the tryptophan salvage genes in *Chlamydia trachomatis*"

**Terminator Fold**

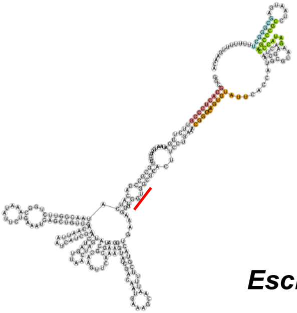

**Anti-Terminator Fold**

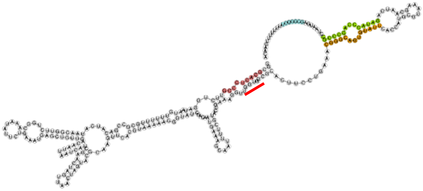

***Escherichia coli* K-12**
***trpL***
Score: **0.63**

**Terminator Fold**

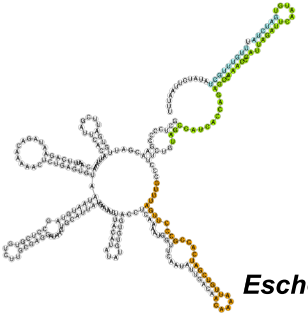

**Anti-Terminator Fold**

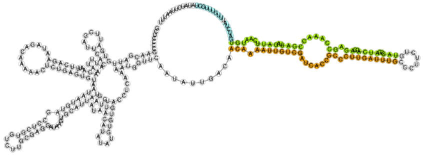

***Escherichia coli* K-12**
***tnaC***
Score: **0.34**

**Terminator Fold**

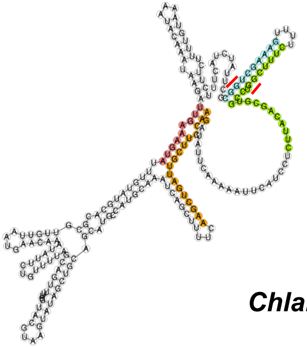

**Anti-Terminator Fold**

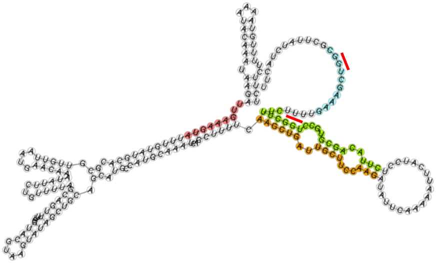

***Chlamydia trachomatis* L2**
***trpL***
Score: **0.37**
